# Epitope order Matters in multi-epitope-based peptide (MEBP) vaccine design: An *in silico* study

**DOI:** 10.1101/2021.06.29.450372

**Authors:** Muthu Raj Salaikumaran, Prasanna Sudharson Kasamuthu, V L S Prasad Burra

## Abstract

With different countries facing multiple waves, with some SARS-CoV-2 variants more deadly and virulent, the COVID-19 pandemic is becoming more dangerous by the day and the world is facing an even more dreadful extended pandemic with exponential positive cases and increasing death rates. There is an urgent need for more efficient and faster methods of vaccine development against SARS-CoV-2. Compared to experimental protocols, the opportunities to innovate are very high in immunoinformatics/*in silico* approaches especially with the recent adoption of structural bioinformatics in peptide vaccine design. In recent times, multi-epitope-based peptide vaccine candidates (MEBPVCs) have shown extraordinarily high humoral and cellular responses to immunization. Most of the publications claim that respective reported MEBPVC(s) assembled using a set of *in silico* predicted epitopes, to be the computationally validated potent vaccine candidate(s) ready for experimental validation. However, in this article, for a given set of predicted epitopes, it is shown that the published MEBPVC is one among the many possible variants and there is high likelihood of finding more potent MEBPVCs than the published candidate. To test the same, a methodology is developed where novel MEBP variants are derived by changing the epitope order of the published MEBPVC. Further, to overcome the limitations of current qualitative methods of assessment of MEBPVC, to enable quantitative comparison, ranking, and the discovery of more potent MEBPVCs, novel predictors, Percent Epitope Accessibility (PEA), Receptor specific MEBP vaccine potency(RMVP), MEBP vaccine potency(MVP) are introduced. The MEBP variants indeed showed varied MVP scores indicating varied immunogenicity. When the MEBP variants were ranked in descending order of their MVP scores, the published MEBPVC had the least MVP score. Further, the MEBP variants with IDs, SPVC_387 and SPVC_206, had the highest MVP scores indicating these variants to be more potent MEBPVCs than the published MEBPVC and hence should be prioritized for experimental testing and validation. Through this method, more vaccine candidates will be available for experimental validation and testing. This study also opens the opportunity to develop new software tools for designing more potent MEBPVCs in less time. The computationally validated top-ranked MEBPVCs must be experimentally tested, validated, and verified. The differences and deviations between experimental results and computational predictions provide an opportunity for improving and developing more efficient algorithms and reliable scoring schemes and software.

## 1. Introduction

The World Health Organization (WHO) announced SARS-CoV-2 as a pandemic in January 2020. Globally, as of date, there have been over 159 million confirmed cases of COVID-19 with the second COVID-19 wave kicking in around the 1st week of March 2021 and registering over 3.3 million deaths. According to a report, *preventive and treatment options* is one of the top two scenarios, the other being *digitalization drive*, which needs global adoption by the post-COVID-19 world to bounce back and get on to the revival path [1].

Since December 2019 the global COVID-19 pandemic has reached unprecedented deaths and death rates, currently continuing its more dreaded second wave. The world has called for preparedness for COVID-19 third wave as well. As of date, there are 287 COVID19 candidate vaccines undergoing trials with 102 candidates under the clinical phase and 185 candidates under the preclinical stage[2]. These vaccine candidates belong to at least fourteen different vaccine platforms [3]. Currently, 133 vaccine candidates of 287 that are undergoing trials belong to the protein subunit platform (~34%) which emphasizes the importance and future potential of peptide-based vaccine candidates. There are 85 protein subunit-based vaccine candidates (~64%) that are under preclinical trials indicating there is a huge opportunity for peptide-based vaccine candidates [4]. Out of a total of nineteen vaccines approved for use, by at least one national regulatory authority(NRA), only one vaccine: EpiVacCorona (Russia NRA), is a peptide-antigen-based vaccine. There are seven vaccines currently approved and listed under WHO Emergency Use Listing (EUL) but no approved vaccine belongs to the protein subunit platform[5].

One of the first efforts of *in-silico* vaccine design was published in 1999, formally launching immunoinformatics [6–8]. One of the first attempts to use multi-epitope-based polypeptides (MEBPs) for HIV-1 vaccine development dates back to 1999 [9]. In recent times MEBPVCs have shown extraordinarily high humoral and cellular responses to immunization. One of the first computational methods for designing MEBP vaccine constructs was published in 2010 [10,11]. Since then numerous articles related to *in silico* design of MEBP vaccine development have been published addressing vaccines against cancer, Chagas disease, filarial diseases, multi-drug resistance, Malaria, TB, and others [12–19]. MEBPVCs are gaining preference as they provide better control over the immunogenic components of the pathogen responsible for causing diseases, better reproducibility, and experimental control [20].

Recently, many novel **multi-epitope-based peptide (MEBP)** vaccine constructs (MEBPVCs) against Spike Protein and or multi-targets of SARS-CoV-2 that causes COVID-19 disease have been designed using *in silico* approaches. Rahmani et. al., 2021, proposed a trivalent (multi-target) MEBPVC that contains new components such as an intracellular delivery agent (TAT) and synthetic epitope (PADRE) in addition to conventional components such as adjuvants (β-defensin 2), the predicted epitopes, and linkers to boost the immune response [21]. The novelty in Saha et. al., 2021 publication is dual-purpose epitopes i.e., each epitope predicted is a B-Cell derived T-cell epitope with a fixed size of ten amino acids. In other words, each predicted epitope triggers a response from both B-Cells and T-Cells simultaneously. This approach keeps the size of MEBPVC small yet efficient, keeping the titers high from both B-cells and T-cells [22]. Similarly, Khairkhah et. al., 2020, in their design, have not used any adjuvants[23]. Table 1 gives a detailed summary of the most recent *in silico* approaches in MEBPVCs against spike protein or multi-targets of SARS-CoV-2.

**Table 1:** The common components, i.e., adjuvants, linkers and predicted epitopes, used in a MEBPV construction. The data provides the components which are commonly being used in design of MEBPVCs with relevant references.

| S.no | Molecular Weight (kDa) | Sequence Length | VCs | Epitope Size (Fixed(F) / Varying (V)) | adjuvants | linkers | special motifs | TLRs | Targets | Reference |
| --- | --- | --- | --- | --- | --- | --- | --- | --- | --- | --- |
| 1 | 75 | 700 | 1 | V | β-defensin 2 | GPGPG, EAAAK, AAY, GGGS, KK | PADRE, TAT, | TLR3,4,8 | S, M, N, E, Nsp8, Nsp3 | [21] |
| 2 | 20 | 183 | 1 | F | β-defensin 2 | EAAAK, AAY, GPGPG | - | TLR8 | S | [22] |
| 3 | 50 | 460 | 5 | V | HSP70, TR-433, RS09 | GPGPG, EAAAK, AAY | - | TLR4 | S | [24] |
| 4 | 17 | 146 | 1 | V | HBD-2 | EAAAK, AAY | -- | TLR-2,3,4 | S | [25] |
| 5 | 30, 37, 50 | 254, 330, 475 | 3 | V | - | KK, AA, GPGPG | - | TLR-2,3,4 | S, M, N | [23] |
| 6 | 50 | 485 | 1 | F | HP-91, HBD-3 | - | - | TLR3,5,8 | S, N, M, E | [26] |

It is a fundamental and proven fact in molecular biology that a change (modification/mutation) in the composition and or order of an individual or a group of amino/nucleic acids in the protein/DNA/RNA sequence could bring substantial change to the fold/3D structure and hence alter the function. In the context of peptide subunit vaccine platforms, this would mean that any change in the order or composition of amino acids of the peptide-based immunogen (especially large sized immunogens), may undergo changes in the fold and 3D structure. This in turn is expected to alter the biophysical and immunological properties of the MEBPVC such as surface accessibility of the epitopes in the MEBP. Any change in the epitope accessibility to the host immune system influences the immunogenicity/antigenicity of the MEBPVCs.

With increasing adoption, strong interdependence of structural biology and immunoinformatics, and with a trend for designing bigger and larger vaccines with sizes ranging from 10 - 100kDa, the emphasis should be to utilize the protein structural biology knowledge for effective vaccine design. The opportunity for structure-based vaccine design leads to next-generation immunogen development [27,28]. In this article, we present a novel methodology to computationally design and validate MEBPVCs. This study indeed provided interesting insights that will help in developing novel MEBP specific design tools and also speedup and improve the current vaccine design methodologies and protocols which is not only the need of the hour but also a need for global preparedness for the future pandemics supporting novel platforms[29].

## 2. Materials and Methods

In general, a MEBP sequence is constructed with three broad components (peptide sequence patterns), namely, Linkers (for example AAY, GPGPG, EAAAK), Adjuvants (beta-defensin 2, HSP70), and predicted target specific MHCI and MHC II Epitopes (oligopeptides with size ranging from 8AA - 20AA). The MEBP sequence (REF_SEQ) published earlier is taken as a reference for this study [22]. Table 5 gives the properties of REF_SEQ MEBPVC.

### 2.1. Construction of MEBP and its variants

The construction of the REF_SEQ is described elsewhere [22]. Using the REF_SEQ another nine MEBPVCs were generated by shuffling the epitope-linker set from REF_SEQ (Fig. 1). The generated MEBP variants are given unique IDs following the format as: SPVC_NNN, where SPVC stands for shuffled peptide vaccine candidate and NNN stands for a unique number. The ten sequences are provided as supplementary data (Supp. 1 file).

**Figure 1:**
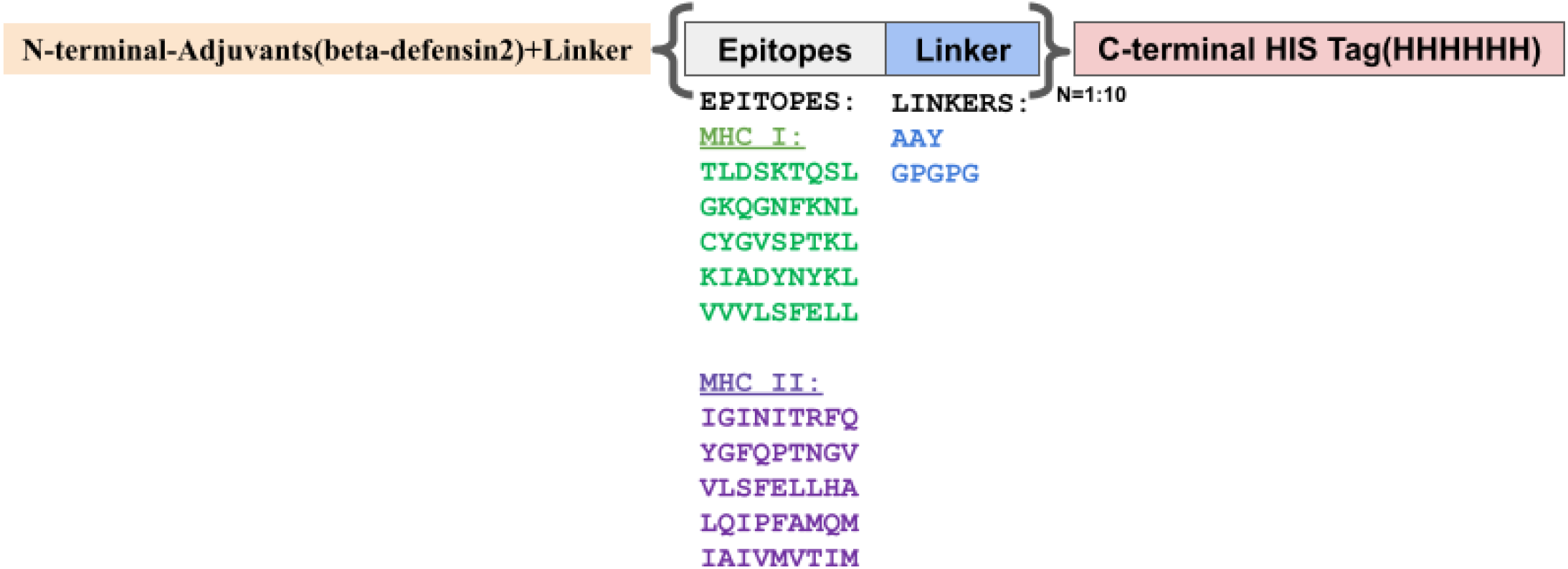
Schematic template of a typical MEBPVC to generate variants from REF_SEQ. The MEBPVC contains five major components: 1. Adjuvant (yellow color), 2. Predicted MHC I Epitopes (Green color) 3. Predicted MHC II Epitopes (Magenta) 4. Linkers(AAY, GPGPG) (Blue color) 5. HIStag(Light red color). The MEBP variants were generated by shuffling the epitopes at the ten (N=1:10) positions.

The following restraints were applied while generating the variants: a) The position and order of N-terminal adjuvant (β-defensin 2) followed by EAAAK (linker) and the C-terminal HIS-tag were kept unchanged. b) The length of the construct was kept unchanged at 183AA only. c) The B-cell-derived T-cell epitope (9AA long) plus the linker (AAY / GPGPG), together, were rearranged manually to create nine MEBPVC variants. d) No distinction was made between MHC-I (green) and MHC-II (magenta) epitopes. The molecular weight, the sequence length, the isoelectric point (pI), the aliphatic index, the number of atoms, half-life, and the chemical formula remained identical for all the variants.

### 2.2. Sequence alignment

The formatting code, all-to-all sequence alignment among the members of the MEBP dataset was done using BioInt [30], Biobhasha (www.biobhasha.org), and BOSv2.0 (Biological Object-Based Software (BOS): An Integrative Biological Programming Environment).

### 2.3. Prediction of immunological and biophysical properties of MEBPVCs

The following relevant immuno- and biophysical properties were predicted using the online/offline tools to compare and develop a rationale for identifying the more potent ones. Table 2 gives the biophysical and immunological properties employed for analysis and their respective servers and tools.

**Table 2:**
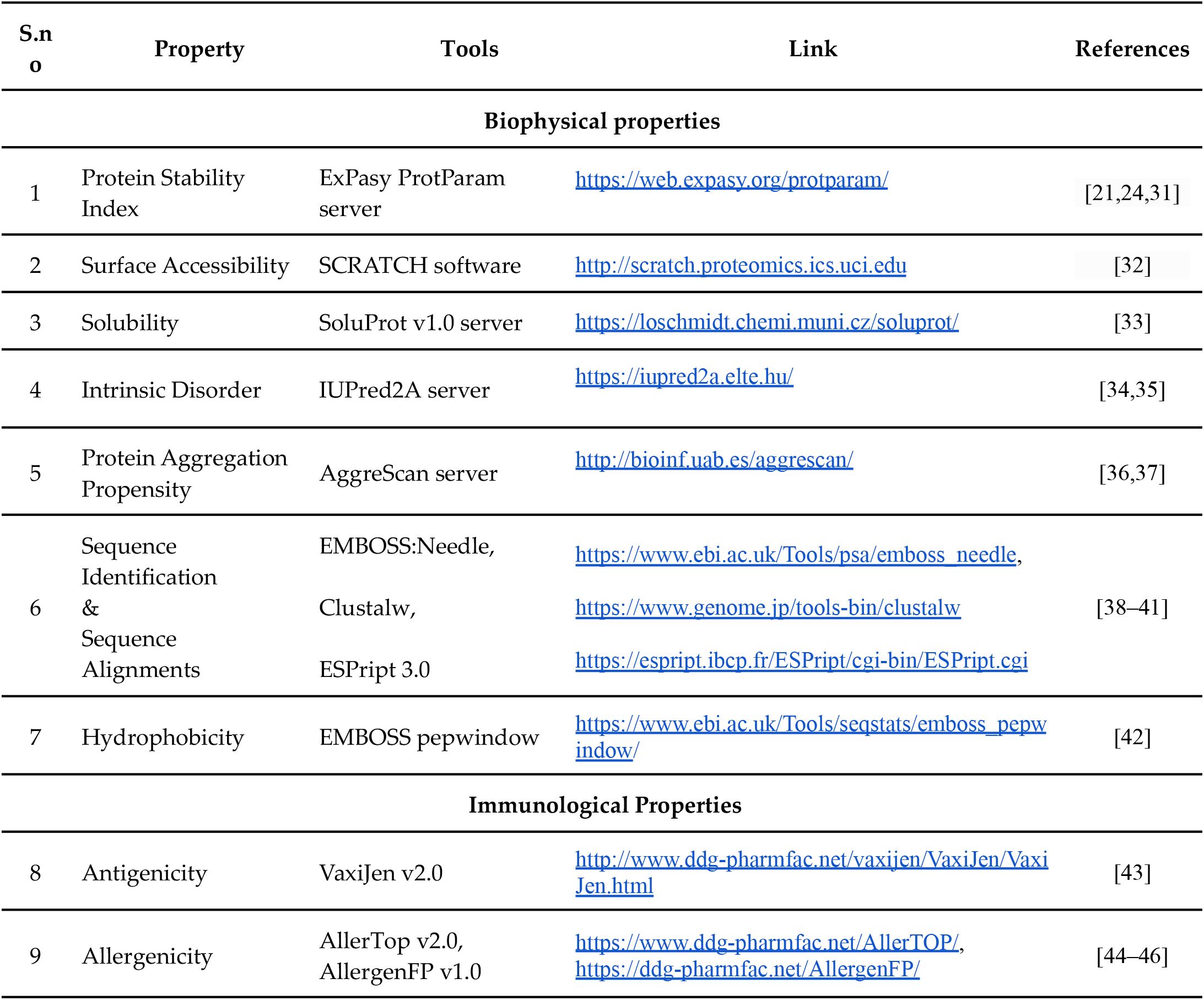
Details about the various biophysical and immunological properties, respective tools and web server links that were used for generating the data.

#### 2.3.1. Antigenicity

Antigenicity is the extent to which the host immune system responds to the antigen (foreign body) triggering both humoral and cellular responses. VaxiJen2.0 server was used to predict the antigenicity of the MEBP variants. The result from VaxiJen 2.0 categorizes the peptide input into either Probable ANTIGEN or Probable NON-ANTIGEN. Only those variants which were categorized as Probable ANTIGEN were selected for further processing.

#### 2.3.2. Allergenicity

Allergenicity is the extent to which an immunogen or antigen induces allergic reactions in the host system resulting in discomfort and or inconvenience or any allergic conditions such as asthma, diarrhea, skin rashes, and others. AllerTop v2.0 server was used to predict the allergenicity of the MEBP variants. The output from AllerTop v2.0 indicates if the input sequence is an allergen or not using the following restricted text values i.e. PROBABLE ALLERGEN or PROBABLE NON-ALLERGEN. Only those variants were selected for further processing which were categorized PROBABLE NON-ALLERGEN.

#### 2.3.3. Peptide/protein stability index

Protein stability is an important property especially to understand the structure-function and activity relationships of a protein. The EXPASY ProtParam server was used for predicting the Instability index of the nine MEBPVCs. The instability index was modified to the stability index by subtracting the score from 100. A score of more than 60 henceforth indicates the input protein to have better stability.

#### 2.3.4. Solvent accessibility

Solvent accessibility is an important feature for understanding and interpreting the structure, activity relationship. The solvent accessibility or the surface exposed residues provide data that helps in various predictions such as protein-protein interactions, receptor-ligand interactions, drug designing, protein folding, and others. Scratch Protein Predictor (http://scratch.proteomics.ics.uci.edu) [47] was used to predict the solvent accessibility of the variants. The output from Scratch Protein Predictor contained residue level accessibility with higher values indicating higher accessibility and vice versa. In the context of peptide vaccine design, higher accessibility and especially the residues with high accessibility in the epitope regions of MEBP is preferred since to elicit an immune response, the epitopes in the vaccine construct should be accessible, be exposed, and be on the surface. The accessibility predictor gives a string output, equal to the length of the sequence, with ‘e’(for each exposed/accessible residue) and ‘b’(for each buried residue). A more meaningful accessibility score in the context of MEBP vaccine design is percent epitope accessibility (PEA) defined and calculated as per the formula given below:

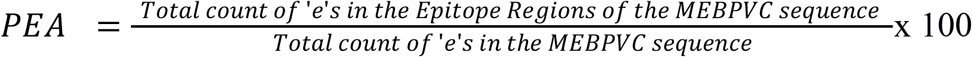

The higher epitope accessibility value is considered favorable.

#### 2.3.5. Solubility

Solubility of protein is an important biophysical property that depends on the amino acid composition and the 3D structure. Solubility influences the production of a protein and its half-life in the cell. Less soluble proteins are a major concern since the proteins synthesized may precipitate out or form inclusion bodies which lead to various disease states. SoluProt v1.0 server (https://loschmidt.chemi.muni.cz/soluprot/) was used to predict the solubility of all the MEBP variants [33]. For each input sequence, i.e. an MEBP variant in the current context, the SoluProt v1.0 server gives a score in the range of 0 −1.0 where > 0.5 score indicates soluble and < 0.5 indicates insoluble peptide. The vaccine constructs with higher solubility (>0.5) were selected for further processing.

#### 2.3.6. Inherent intrinsic disorder

Intrinsically disordered proteins (IDPs) (a.k.a intrinsically unstructured proteins (IUPs)) are proteins that deviate from the dogma that every protein has a rigid 3D structure. IDPs or intrinsically disordered regions (IDRs) regulate many important biological functions such as transcription regulation, tissue-specific expression, and other signal transduction pathways. IDRs show high conformational changes, influence the stability of the protein, and affect the binding modes during ligand-receptor interactions. These regions play an important role in protein-protein interactions. IUPred2A server (https://iupred2a.elte.hu/) was used to predict the disorder of the MEBP variants [35]. The output has residue-wise disorder values and contiguous IDRs. A value >0.5 is considered disordered and the value <0.5 is considered ordered. An average disorder for each MEBP variant was calculated using the residue-wise disorder. The variants with the low average disorder are considered for further analysis.

#### 2.3.7. Protein aggregation

Protein aggregation is a biological process in which protein/peptide subunits instead of forming regular and functional assemblages, misfold, aggregate (intra- or extracellularly), and precipitate. Protein aggregation is one of the important phenomena implicated in diseases such as Parkinson’s, Alzheimer’s, and prion diseases. To predict the aggregation for the variants, we used the AGGRESCAN server (http://bioinf.uab.es/aggrescan/). Variants with lower predicted aggregation were considered for further analysis.

#### 2.3.9. Hydrophobicity

Hydrophobicity is the ‘water-hating/avoiding/repelling’ property of molecules. Hydrophobic amino acids tend to fold and shrink together to minimize contact with the solvent water or hydrophilic surroundings. The hydrophobic effect is a well-known important property to understand the 3D structure of a protein[48]. Hydrophobic interactions are an important driving force in protein folding hence the overall 3D structure. The shape determines the function of the protein. In the context of MEBPs which are the potential immunogens, higher hydrophobicity indicates more globularity, rigid 3D structure, and associated accessible surface residues. Kyte and Doolittle’s method was used to generate the Hydrophobicity values by using EMBOSS pepwindow server (https://www.ebi.ac.uk/Tools/seqstats/emboss_pepwindow/). The outputs were screened and filtered for higher hydrophobicity and used for further processing.

### 2.4. Dosage versus immune response simulation

C-IMMSIM (http://150.146.2.1/C-IMMSIM/index.php) is one of the most commonly used in *in silico* immune simulation servers. Given the vaccine candidate, the dosage volume, and dosage intervals, C-IMMSIM predicts the humoral and cellular immune responses in the host system over a period of time. The server uses machine learning techniques to predict the immune responses of immunogens. The time step of injection, 3 doses, was set at 1 hr, 84 hrs (3.5 days), and 168 hrs (7 days). The simulation steps and simulation volume were kept at 1100 and 10.

### 2.5. Structural Studies

#### 2.5.1. Ab initio 3D structure prediction

A local installation of I-TASSER-suite was used to predict the tertiary structures of the MEBP variants. The suite uses the Local Meta-Threading server (LOMET) to find out the suitable template structure from the input sequence. In the second step, the server performs the template-based fragment assembly simulation to create a full structure and the final step is it will produce the top five predicted models with TM-score (Quantitative assessment of similarity between protein structures) and C-score (confidence score for estimating the quality of predicted models).

#### 2.5.2. Molecular docking

The protein-protein docking was performed in Discovery studio (Z_DOCK module). The receptors (TLR4 and TLR8) were downloaded from RCSB with PDBIDs:4G8A, 3W3M, and the MEBP variants were used as ligands. Other parameters were kept unchanged (default values). The docking resulted in ~2000 clusters with each cluster containing ~70 to 80 poses of ligands (MEBP variants) with the receptor (TLR4 and TLR8). The top poses with the highest Z_RANK-SCORE were selected to find R_DOCK-SCORES (Refine docked protein). The R_DOCK uses CHARMm energy to optimize docked poses produced by the Z_DOCK module. The top hits from the R_DOCK module were selected for further simulation studies.

#### 2.5.3. Molecular dynamics simulation

The top-ranked poses of MEBP variants with TLR4 and TLR8 were selected for running MD simulations. iMODS server was used to run the simulations. The server uses the Cα coarse grain model for complexes to calculate the mobility of the proteins based on ENM(elastic network model). It also calculates the mode variance, B-factor plot, eigenvalue, and covariance. It also uses NMA(Normal mode analysis) for finding out the protein motion in the internal coordinates.

#### 2.5.4. Implementation of a scoring scheme for ranking and discovering the best MEBPVC from the library of variants

The values obtained for the ten MEBPVC variants for the above discussed properties were grouped into a) positive influencers (Stability, Accessibility, Solubility, Hydrophobicity, Antigenicity, ZrankScore, binding Affinity) b) negative influencers (disorder and aggregation). All the values of a property were normalized using normalization by averaging i.e.

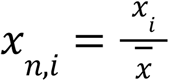

where

x = original value of the property
n = normalized value
i = index of the variant
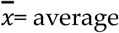, all the variants.

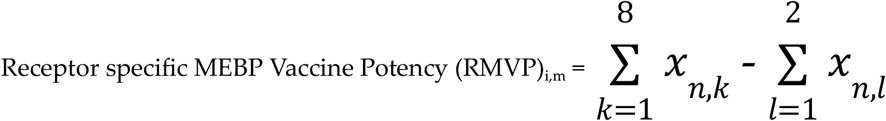

- where k, l are the normalized values of each positive and negative influencers (property of the variant) respectively.
- where m is the receptor i.e. TLR4(1) or TLR8(2).

Final MVP for each variant, i, was calculated by summing individual RMVPs.

## 3. Results

The purpose of a vaccine (MEBPVC in the current context) is to elicit a strong response from the host immune system against a disease (COVID-19) releasing various neutralizing antibodies which stayput in the body to protect the host from any repeat infection henceforth. Toll-like receptors (TLRs) are the common receptors that interact with the immunogen (MEBPVC) and trigger the downstream response and release of neutralizing antibodies. From the informatics point of view, to correlate the properties, to derive relationship with immunogenicity and changed epitope positions of the MEBPVC, the necessary data was generated at four levels: a) sequence level, b) 3D structure level, c) receptor-ligand interaction level and d) dosage versus immune response level. The data generated thus, is analysed to understand if the changed epitope positions influenced the various properties and eventually the immunogenicity. Prior to the analysis, to establish the fact that the change in the order/positions of the epitopes in a MEBPVC changes the immunogenicity, one has to first assess the diversity among the MEBP variants. To assess the same, pairwise alignment, multiple sequence alignment (MSA), and structure alignment were performed on the ten variants. The results are discussed below.

### 3.1. Sequence & structural similarities between the MEBP variants

The sequence identity among the variants varied between 39.3 - 74% with the root mean squared deviations (RMSDs) ranging between 3.38 and 17.3 Å (Table 3) where, in the domain of structural biology, RMSD is the common quantitative descriptor of structural similarity between two 3D structures. The average sequence identity between the variants is around 49% which is understandable considering the common adjuvants (45AA), linkers(AAY:3×5(copies)=15AA, GPGPG:5×5(copies)=25AA), and HIS tags(6AA) which make up to 49% (91AA) of the 183 AA long MEBPVC.

**Table 3:** For ready comparison between the MEBP variants, the data is presented as Percent sequence identity (lower triangle: below 100.00 diagonal) and Root Mean Square Deviation (RMSD) (upper triangle: above 100.00 diagonal). Lighter shades of a color indicate lower RMSD (higher structural similarity) and lower sequence identity whereas darker shades of the colors indicate higher RMSD (lower structural similarity) and higher sequence identity.

|  | SPVC_R<br>EF | SPVC_20<br>6 | SPVC_21<br>4 | SPVC_32 | SPVC_35<br>7 | SPVC_53<br>7 | SPVC_38<br>3 | SPVC_56<br>5 | SPVC_44<br>6 | SPVC_38<br>7 |
| --- | --- | --- | --- | --- | --- | --- | --- | --- | --- | --- |
| SPVC_R<br>EF | 100.00 | 13.83 | 17.30 | 13.30 | 6.01 | 14.48 | 16.20 | 13.34 | 7.14 | 12.60 |
| SPVC_20<br>6 | 46.24 | 100.00 | 16.32 | 10.27 | 14.01 | 8.75 | 12.88 | 8.82 | 13.15 | 14.79 |
| SPVC_21<br>4 | 64.32 | 56.28 | 100.00 | 11.77 | 14.87 | 13.07 | 16.60 | 11.79 | 7.90 | 4.92 |
| SPVC_32 | 54.10 | 49.47 | 57.30 | 100.00 | 7.19 | 13.24 | 15.21 | 12.75 | 3.38 | 15.08 |
| SPVC_35<br>7 | 59.89 | 49.47 | 50.27 | 49.20 | 100.00 | 16.06 | 6.67 | 13.95 | 6.38 | 15.52 |
| SPVC_53<br>7 | 55.68 | 43.09 | 51.34 | 70.27 | 50.81 | 100.00 | 11.60 | 14.23 | 3.72 | 7.18 |
| SPVC_38<br>3 | 57.30 | 60.64 | 73.51 | 53.55 | 48.13 | 52.97 | 100.00 | 13.77 | 4.04 | 15.76 |
| SPVC_56<br>5 | 50.54 | 49.47 | 53.19 | 71.89 | 51.35 | 65.95 | 52.97 | 100.00 | 13.63 | 13.66 |
| SPVC_44<br>6 | 46.77 | 54.64 | 46.24 | 46.28 | 41.08 | 48.68 | 47.59 | 47.85 | 100.00 | 15.51 |
| SPVC_38<br>7 | 58.12 | 54.64 | 39.13 | 47.06 | 56.15 | 42.47 | 50.27 | 50.27 | 55.14 | 100.00 |

Interestingly, the MEBP variants with IDs, SPVC_214 and SPVC_383, have 73.51% (highest) sequence identity(Fig. 2a). When the epitope positions between SPVC_214 and SPVC_383 were studied, the same epitopes were present at the 2nd, 6th, 8th, 9th and 10th positions (Table 4).

**Figure 2:**
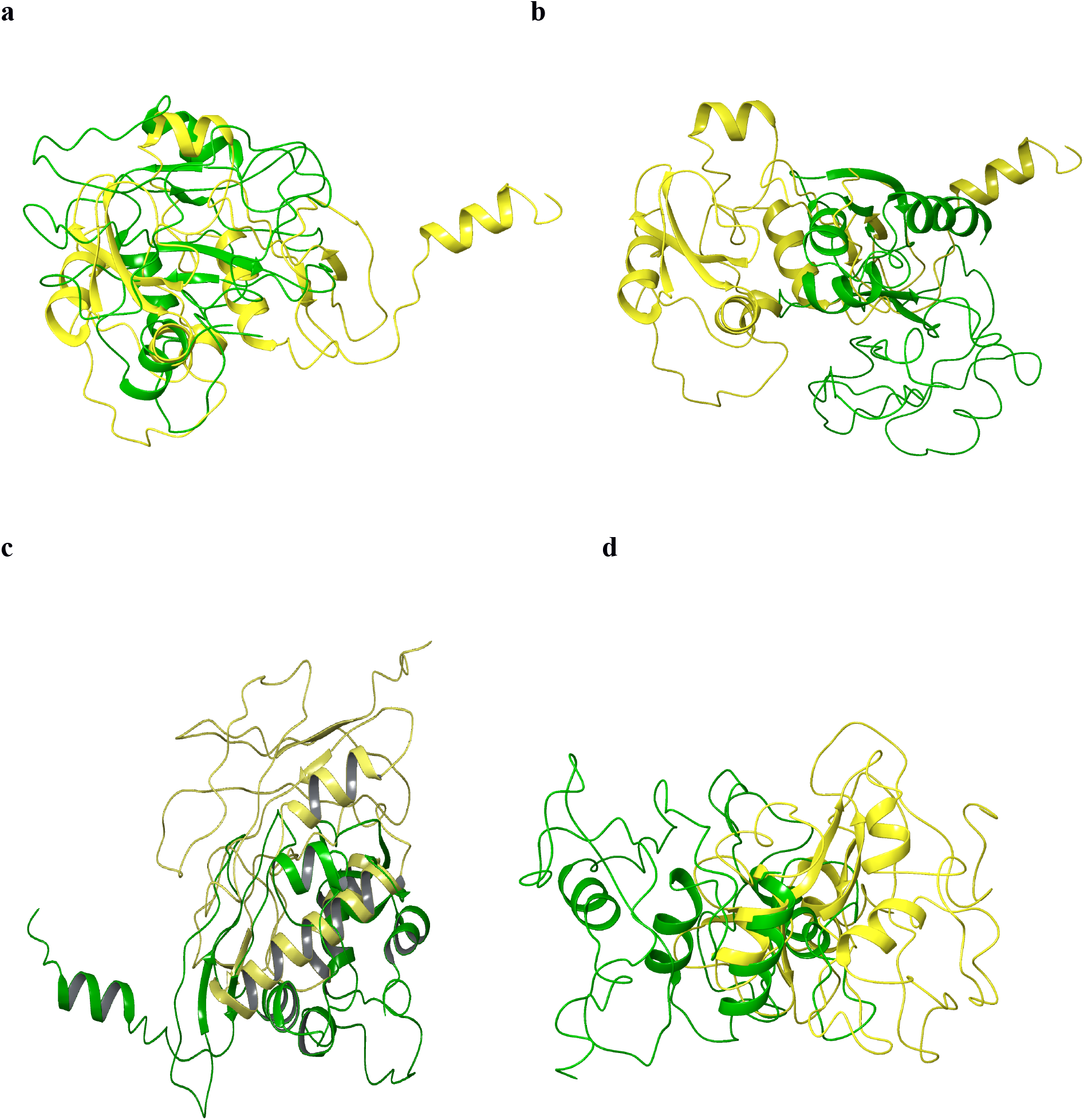
Structure alignment: a) SPVC_214(yellow) and SPVC_383(green), 73.51% (SeqId) b) SPVC_214(yellow) and SPVC_387(green), 39.13% c) REF_SEQ(yellow) and SPVC_214(green), 17.3 Å d) SPVC_32(yellow) and SPVC_446(green) 3.38 Å.

**Figure 3.**
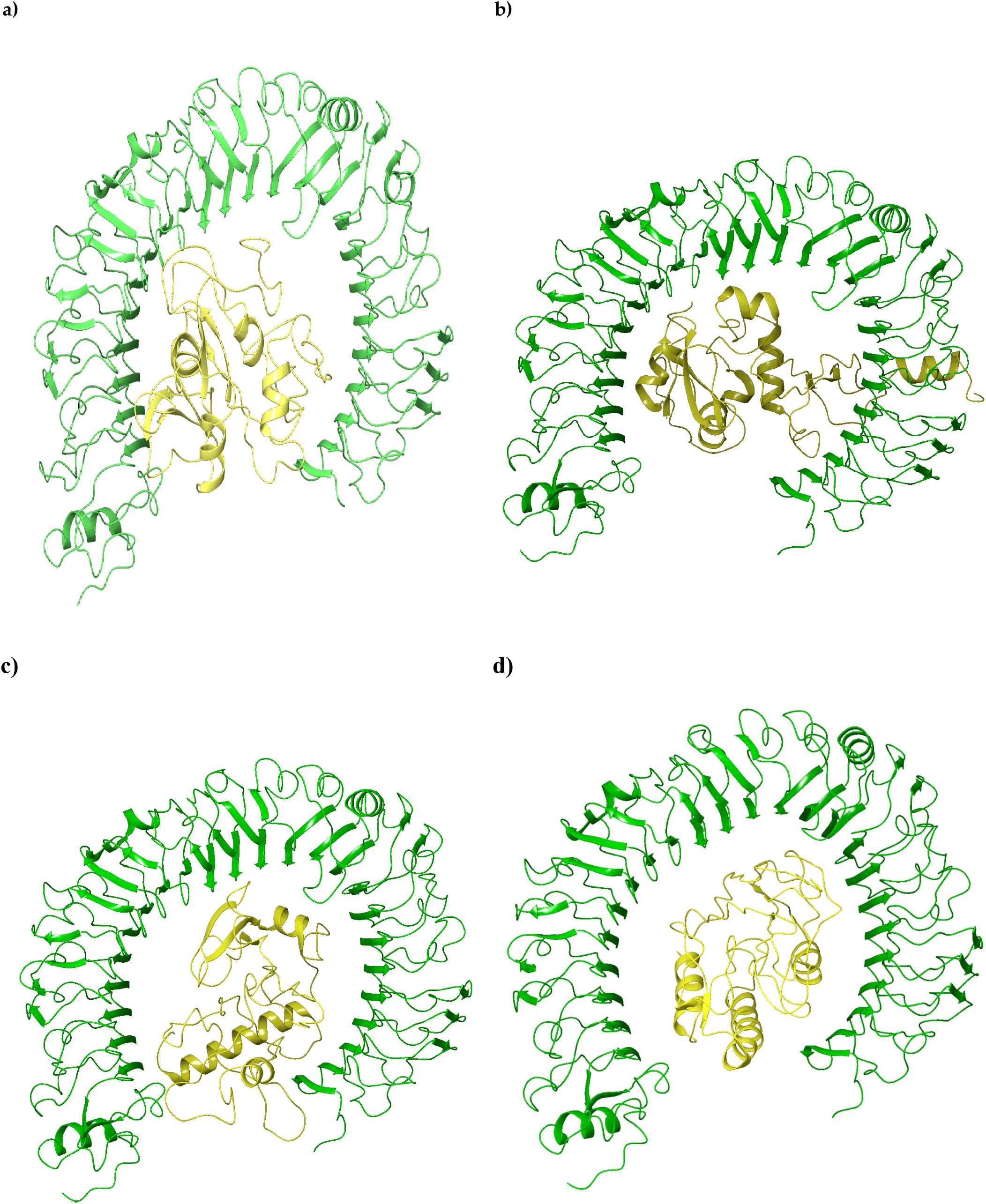
Docked ligand of a)SPVC_32, b)SPVC_214, c)SPVC_565, d)REF_SEQ(yellow) with TLR4(green).

**Table 4 :**
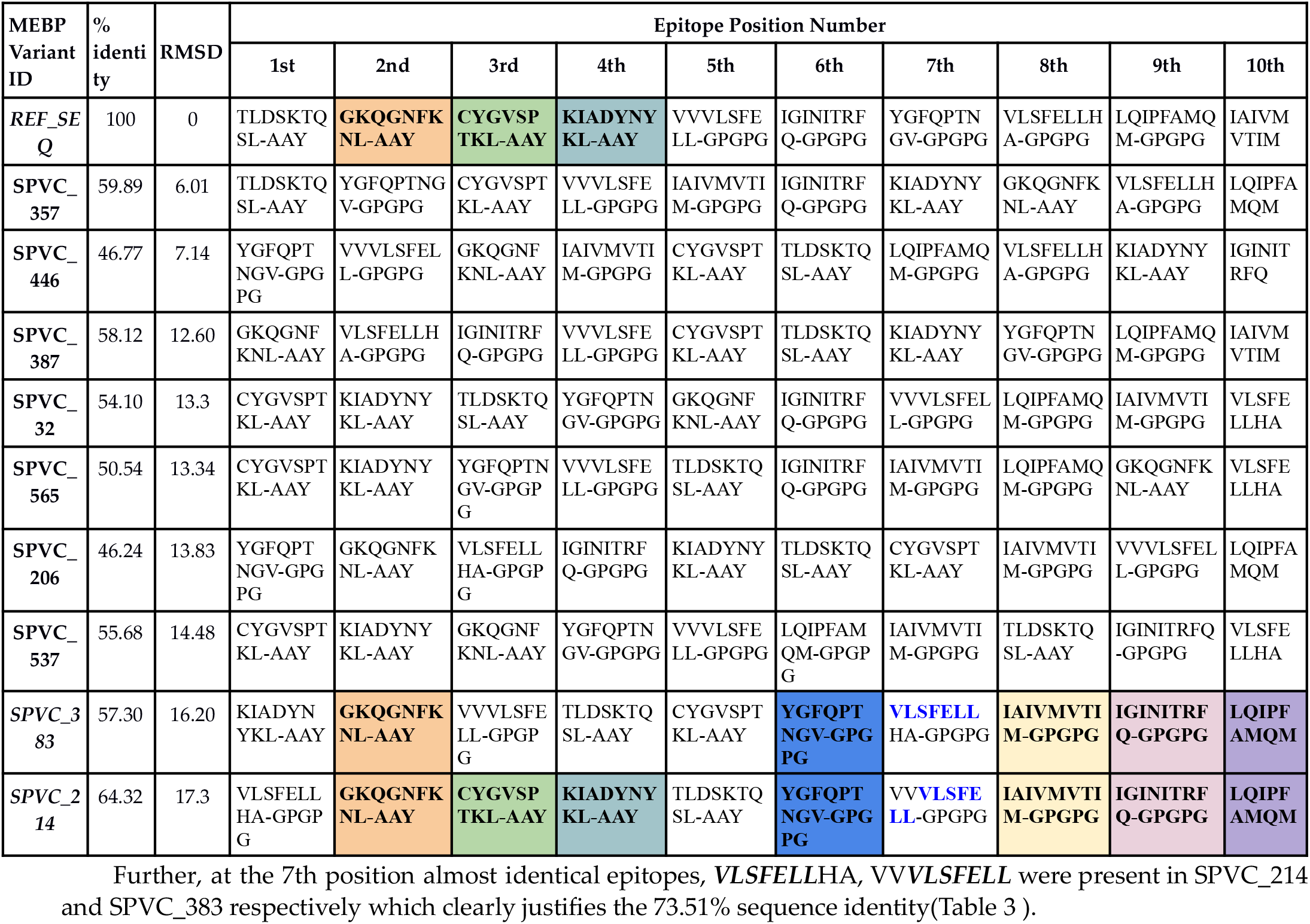
Comparative epitope positions in the ten MEBP variants with % identity and RMSD with reference to REF_SEQ MEBPVC. Three comparisons are highlighted in this table. 1) REF_SEQ vs SPVC_383, 2) REF_SEQ vs SPVC_214 and 3) SPVC_383 vs SPVC_214. Epitopes that are found at same position across the variants are highlighted (Same color indicates same epitopes found at same position)

However, their RMSD is one among the highest, i.e. 16.60 Å (Table 3) indicating very dissimilar structures which is contrary to the notion in homology modeling that is, high sequence identity (>30%) indicates high structural similarity and same function.

The pair with the least sequence identity (39.13%) is SPVC_214 and SPVC_387(Fig. 2b). Their RMSD is 4.92 Å indicating reasonable structural similarity though sequences are not very similar. The pair with the highest RMSD i.e.17.3 Å are REF_SEQ and SPVC_214. With such high RMSD, it is commonly expected to have very little sequence similarity between the two sequences. However, they have a sequence identity of around 65%. Further analysis revealed that the same epitopes were seen at 2nd, 3rd, and 4th positions in the two MEBP constructs (REF_SEQ and SPVC_214)(Table 3) justifying the above-average sequence identity. Similarly, the pair with the lowest RMSD i.e. 3.38 Å are SPVC_32 and SPVC_446 with their sequence identity of only 46.24% only. Fig. 2c,d show structure alignments of the MEBP variant-pair with lowest (3.38 Å, SPVC_32 and SPVC_446) and highest (17.3 Å, REF_SEQ and SPVC_214) RMSDs. These typical cases are clearly indicating that there are changes in the 3D structures of the MEBP variants on changing the positions/order of the epitopes in the MEBPVCs.

### 3.2. Immunological and Biophysical Properties of the MEBP Dataset

#### 3.2.1. Immunological Properties

##### Antigenicity, allergenicity

The antigenicity scores from Vaxijen 2.0 server indicate that all the MEBPVCs are probable antigens with a range between 0.62 to 0.78 (Table 5).

**Table 5:** The raw data of the Immuno- and Biophysical properties of the ten MEBP variants for ready reference.

| MEBP Variant ID | Stability | Accessibility | Solubility | Disorder | Aggregation | Hydrophobicity | Antigenicity | Sequence Identity (%) | RMSD (Str. Similarity Score) |
| --- | --- | --- | --- | --- | --- | --- | --- | --- | --- |
| REF_SEQ | 74.52 | 36.36 | 0.82 | 0.17 | 3.30 | 0.20 | 0.62 | 0.00 | 0.00 |
| SPVC_206 | 73.38 | 35.87 | 0.82 | 0.17 | 2.80 | 0.22 | 0.65 | 55.60 | 13.83 (0.31) |
| SPVC_214 | 73.17 | 36.56 | 0.82 | 0.15 | 2.80 | 0.19 | 0.62 | 64.70 | 17.3 (0.14) |
| SPVC_32 | 75.83 | 35.87 | 0.83 | 0.16 | 2.70 | 0.21 | 0.66 | 61.10 | 13.3 (0.34) |
| SPVC_357 | 73.84 | 35.79 | 0.82 | 0.17 | 2.80 | 0.22 | 0.63 | 64.10 | 6.01 (0.70) |
| SPVC_537 | 76.97 | 36.84 | 0.82 | 0.16 | 2.70 | 0.21 | 0.68 | 58.20 | 14.48 (0.28) |
| SPVC_383 | 73.84 | 37.23 | 0.82 | 0.16 | 2.80 | 0.22 | 0.65 | 60.00 | 16.20 (0.19) |
| SPVC_565 | 75.83 | 35.79 | 0.82 | 0.17 | 2.70 | 0.21 | 0.66 | 57.90 | 13.34 (0.33) |
| SPVC_446 | 76.05 | 36.56 | 0.73 | 0.16 | 3.10 | 0.24 | 0.78 | 50.70 | 7.14 (0.64) |
| SPVC_387 | 73.84 | 35.63 | 0.81 | 0.15 | 3.30 | 0.22 | 0.67 | 55.70 | 12.60 (0.37) |

The sequence, SPVC\_446 has the highest antigenicity(0.78) and SPVC\_214 (0.62) has the lowest antigenicity. All the allergenicity scores predicted using AllerTop v2.0 and AllergenFP v1.0 servers indicate that all the variants are non-allergens hence the allergen column was omitted from Table 5.

#### 3.2.2. Biophysical Properties

##### Stability

The stability scores of MEBPVC variants fall between 73.17 to 76.97 which is the first parameter used as a filter. As a rule, all the MEBPVC variants must be predicted as stable which is found to be true for all the variants in the dataset. SPVC_214 is predicted to have the lowest and SPVC_537, to have the highest stability scores. The variations in the stability scores, though not very dispersed, indicate that change in the order of epitopes changes the stability of the vaccine construct.

##### Solubility

The next important biophysical property is solubility. Less soluble proteins are a major concern since the proteins synthesized may not fold to the right structure and hence lose the activity and function and are observed that they precipitate out or form inclusion bodies which leads to various disease states. The solubility scores range from 0 - 1.0 where > 0.5 score indicates soluble and < 0.5 indicates insoluble peptide. In our case, all the MEBP variants had solubility scores ranging from 0.73 to 0.83 indicating all are soluble. The SPVC_446 variant has the lowest solubility and the SPVC_32 variant has the highest solubility. These solubility scores also indicate that the order of epitopes in the MEBPVC is important and crucial in the design of a good vaccine candidate.

##### Accessibility

Solvent accessibility is an important feature which, in the current context, has direct implications in eliciting the immune response in the host. The higher the epitope accessibility the more immunogenic the vaccine candidate. For the MEBP variants, the percent epitope accessibility ranged between 35 to 37. The variant SPVC_387 has the lowest accessibility and SPVC_383 has the highest accessibility.

##### Disorder

In our MEBP variants disorder ranges between 0.15 to 0.16 and hence all variants are considered ordered. The MEBPVC sequence SPVC_357 has high disorder and SPVC_387 has low disorder among the variants. Low disorder is considered favourable for better vaccine design.

##### Aggregation

The predicted aggregation propensities ranged from 2.7 - 3.3 with lower values considered favorable. The sequences REF_SEQ and SPVC_387 have the highest aggregation propensity and SPVC_32, SPVC_565, SPVC_537 have the lowest propensity. Table 5 gives further details.

##### Hydrophobicity

Higher hydrophobicity shows better globularity, better accessible surface residues, and rigid 3D structure. The predicted hydrophobicity values ranged from 0.194 - 0.239, higher hydrophobicity values are considered more favorable. The sequence SPVC_446 has the highest hydrophobicity and SPVC_214 has the lowest hydrophobicity.

### 3.3. Comparative Docking analysis of MEBP Variants

The properties compared in the previous sections were sequence-based. The analysis proved that the order of the epitopes indeed influenced the solubility, accessibility, disorder, and aggregation properties. To make the analysis more complete and comprehensive, the following sections explore the docking and MD simulation studies using the ab initio modeled 3D structures of the MEBP variants. In the previous section, the MEBP models (variants) and their RMS deviations were discussed.

Among the family of Toll-Like Receptors, TLR8 and TLR4 are the most common receptors interacting with antigens/immunogens triggering an immune response from the host system to fight the immunogen. TLR8 plays an important role in the generation of effective immune responses in humans. TLR8 also senses the single-stranded RNA of viruses in the endosome and is predominantly expressed in the lungs. TLR4 plays an important role in the regulation of myocardial function, fibroblast activation, and acute inflammation by immune cells. Both the receptors are implicated in COVID-19. Table 6 shows the docking scores, binding affinities, minimization energies for both the receptors (TLR8 and TLR4) in complex with the MEBP variants.

**Table 6.**
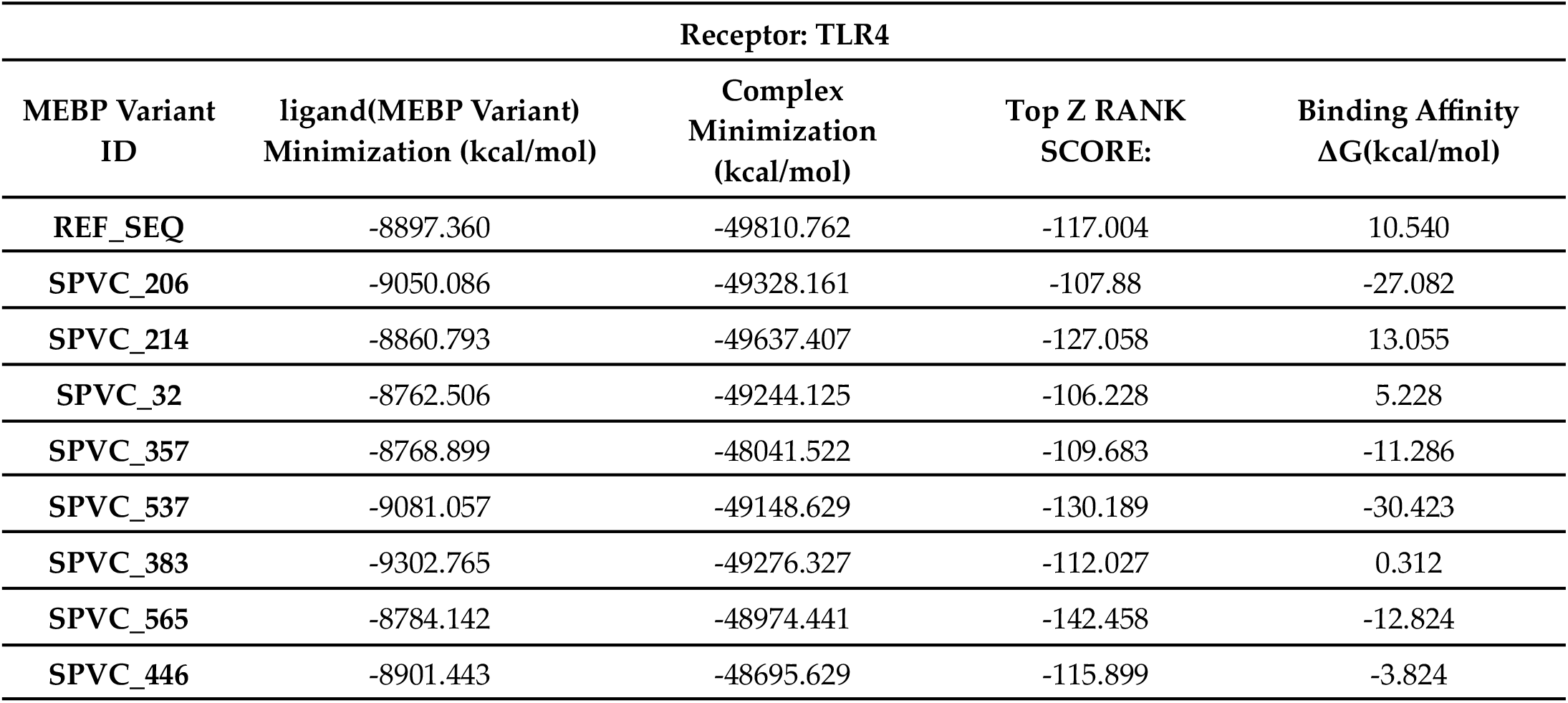
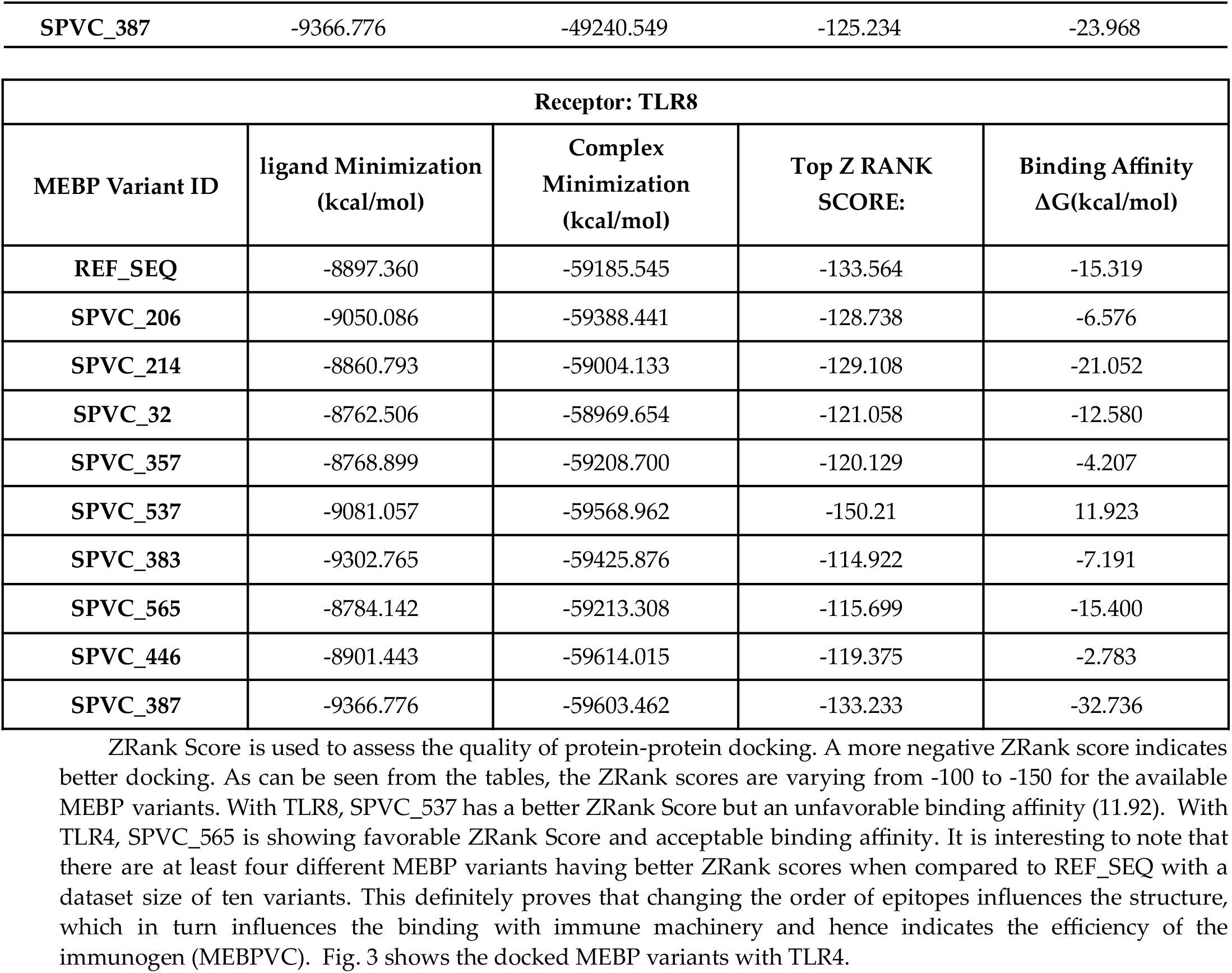
Docking scores, binding affinities, minimization energies of MEBP variants with both the receptors (TLR4 and TLR8).

| Receptor: TLR4 |  |  |  |  |
| --- | --- | --- | --- | --- |
| MEBP Variant ID | ligand(MEBP Variant) Minimization (kcal/mol) | Complex Minimization (kcal/mol) | Top Z RANK SCORE: | Binding Affinity $\Delta G$(kcal/mol) |
| REF_SEQ | -8897.360 | -49810.762 | -117.004 | 10.540 |
| SPVC_206 | -9050.086 | -49328.161 | -107.88 | -27.082 |
| SPVC_214 | -8860.793 | -49637.407 | -127.058 | 13.055 |
| SPVC_32 | -8762.506 | -49244.125 | -106.228 | 5.228 |
| SPVC_357 | -8768.899 | -48041.522 | -109.683 | -11.286 |
| SPVC_537 | -9081.057 | -49148.629 | -130.189 | -30.423 |
| SPVC_383 | -9302.765 | -49276.327 | -112.027 | 0.312 |
| SPVC_565 | -8784.142 | -48974.441 | -142.458 | -12.824 |
| SPVC_446 | -8901.443 | -48695.629 | -115.899 | -3.824 |
| SPVC_387 | -9366.776 | -49240.549 | -125.234 | -23.968 |

| Receptor: TLR8 |  |  |  |  |
| --- | --- | --- | --- | --- |
| MEBP Variant ID | ligand Minimization<br>(kcal/mol) | Complex<br>Minimization<br>(kcal/mol) | Top Z RANK<br>SCORE: | Binding Affinity<br>$\Delta G$(kcal/mol) |
| REF_SEQ | -8897.360 | -59185.545 | -133.564 | -15.319 |
| SPVC_206 | -9050.086 | -59388.441 | -128.738 | -6.576 |
| SPVC_214 | -8860.793 | -59004.133 | -129.108 | -21.052 |
| SPVC_32 | -8762.506 | -58969.654 | -121.058 | -12.580 |
| SPVC_357 | -8768.899 | -59208.700 | -120.129 | -4.207 |
| SPVC_537 | -9081.057 | -59568.962 | -150.21 | 11.923 |
| SPVC_383 | -9302.765 | -59425.876 | -114.922 | -7.191 |
| SPVC_565 | -8784.142 | -59213.308 | -115.699 | -15.400 |
| SPVC_446 | -8901.443 | -59614.015 | -119.375 | -2.783 |
| SPVC_387 | -9366.776 | -59603.462 | -133.233 | -32.736 |

### 3.4. Simulation analysis of MEBPVC variants complexed with TLRs

The objective has been to see if there is any influence of change in the order of epitopes on binding. The earlier sections covered the results of sequence- and structure-based analysis. This section discusses the results of MD simulations to understand the flexibility, stability, and binding between the MEBP variants and the receptors. the iMODS simulation server was used for running fast and coarse simulation studies to understand the differences in the flexibility and stability of the complexes of MEBP variants with receptors, TLR8 and TLR4, of the host immune system. Simulations were run with the two receptors, in complex with each MEBP variant (Supp. 2 Fig. S1-20). The deformability plots show normal flexibilities in both receptors, TLR8 and TLR4. The MEBP variants with TLR4 seem more stable as compared to TLR8 complexes. Further examination shows that some MEBP variants formed more favorable complexes with TLR8 compared to others. For example, SPVC_214 and SPVC_32 have less overall deformability compared to other variants. SPVC_206, SPVC_357, SPVC_565 show high deformability compared to REF_SEQ. With TLR4, SPVC_32, SPVC_206, SPVC_565 show high deformability compared to REF_SEQ. However, SPVC_214, SPVC_537, and SPVC_446 show relatively less deformability. Deformability also indicates the possible hinge regions. TLR8 shows a possible hinge region at around 400 - 420 AAs. There are no such hinge region indications seen in TLR4. Overall, the simulation studies also prove that the complexes are stable however, the variants are unique indicating that the epitope positions alter the immunogenicity.

### 3.5. Dosage versus immune response simulation analysis

As the last of tasks, we performed dosage vs immune response simulation for the ten vaccine constructs (MEBP variants) using the C-ImmSim server with the same objective, to see if the variants trigger different responses than REF_SEQ, if so will they be more potent or less. Two simulation experiments were done: a) with adjuvant and HIS-tag and b) without adjuvant and HIS-tag. The MEBPVC variants had all the parameters within the optimal and recommended ranges for them to be considered as a potent vaccine candidate individually with the exception of SPVC_387(Supp. 3 Fig. S19) (without adjuvants+HIS-tag). A common observation has been that a repeated exposure led to an overall increase in the immune response and a decrease in the antigenic load (Supp. 3 Fig. S1-20).

Few observations are presented here. When compared to REF_SEQ, all other variants trigger strong antibody (especially IgM or IgM+IgG) responses with their 1st dose (exposure). Of all the variants, SPVC_214 is seen to trigger the highest titers of IgG+IgM. Of all the constructs, REF_SEQ triggers the weakest. The titers reach ~650000 counts per ml for SPVC_214 and others but only ~580000 counts per ml for REF_SEQ.

It is interesting to note that variants without adjuvants+HIS-tag seem to trigger more strongly than with adjuvants+HIS-tag. The antibody titers reach 90000 counts per ml without adjuvants+HIS-tag as compared to only 20000 counts per ml with adjuvants+HIS-tag on exposure to 1st dose of SPVC_214. The IgM+IgG titers reach ~760000 counts per ml on the last (third) exposure. A similar trend is seen for all other variants as well where without adjuvants+HIS-tag are triggering a better immune response. Fig. 4(a,b,c) & Fig. 5(a,b,c) shows the level of immunoglobulins (with and without adjuvant+HIS-tag) at two different doses(1st, 2nd, and 3rd).

**Figure 4a:**
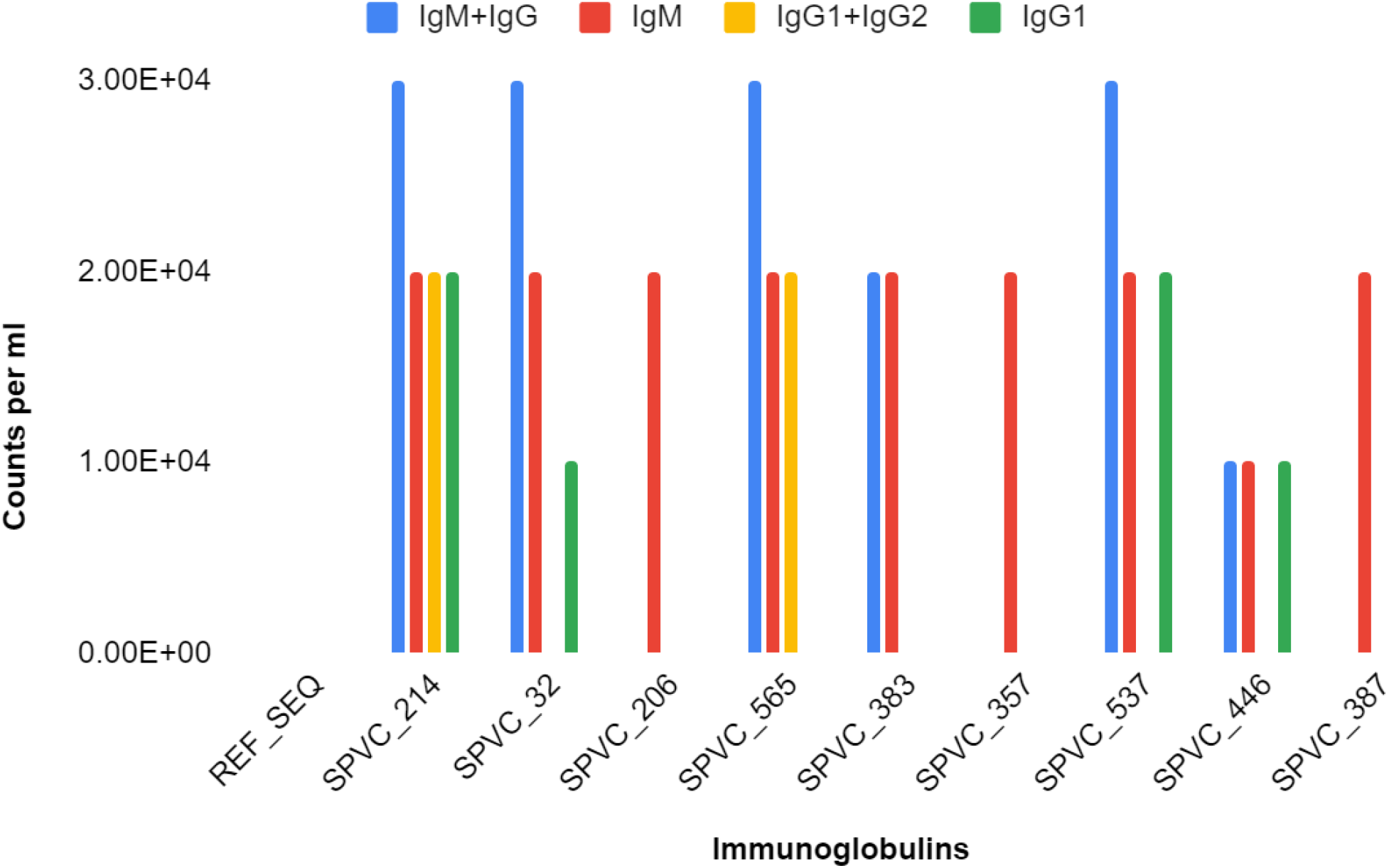
Immunoglobulin counts of MEBP variants (with adjuvants+HIS-tag) after 1st Dose(X axis-Vaccine constructs, Y axis - counts per ml)

**Figure 4b:**
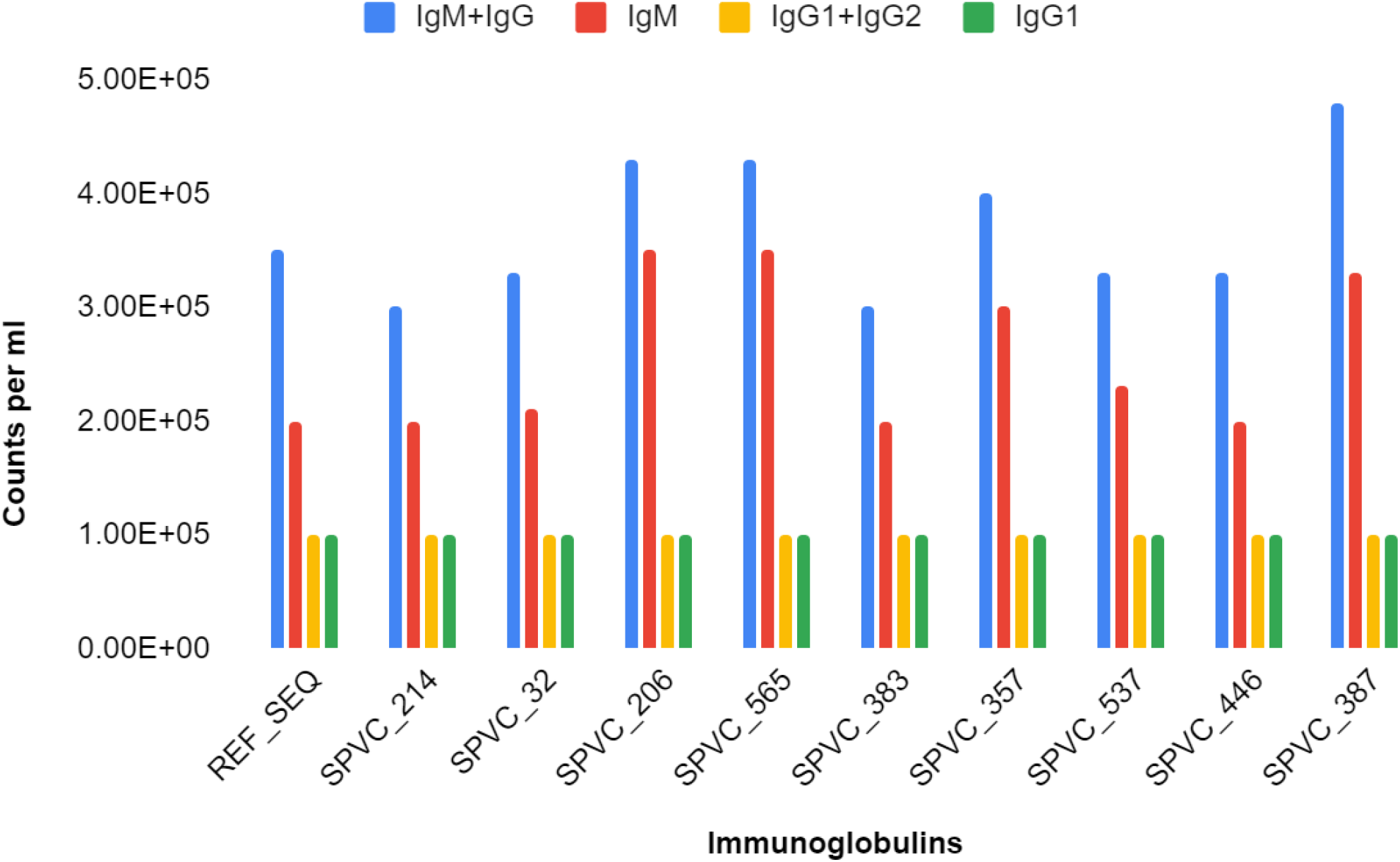
Immunoglobulin counts of MEBP variants (with adjuvants+HIS-tag) after 2nd Dose (X axis-Vaccine constructs, Y axis - counts per ml)

**Figure 4c:**
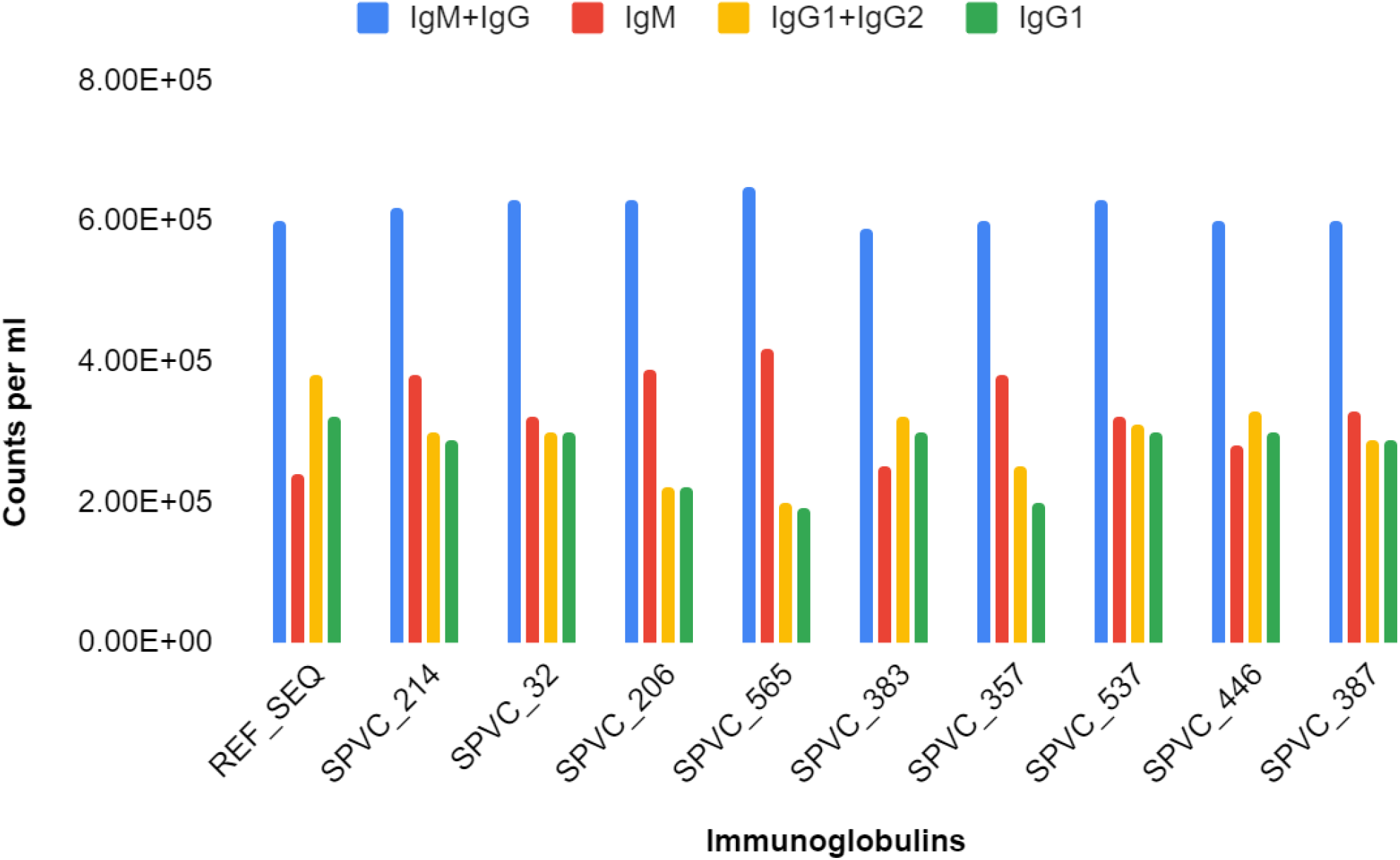
Immunoglobulin counts of MEBP variants (with adjuvants+HIS-tag) after 3rd Dose (X axis-Vaccine constructs, Y axis - counts per ml)

**Figure 5a:**
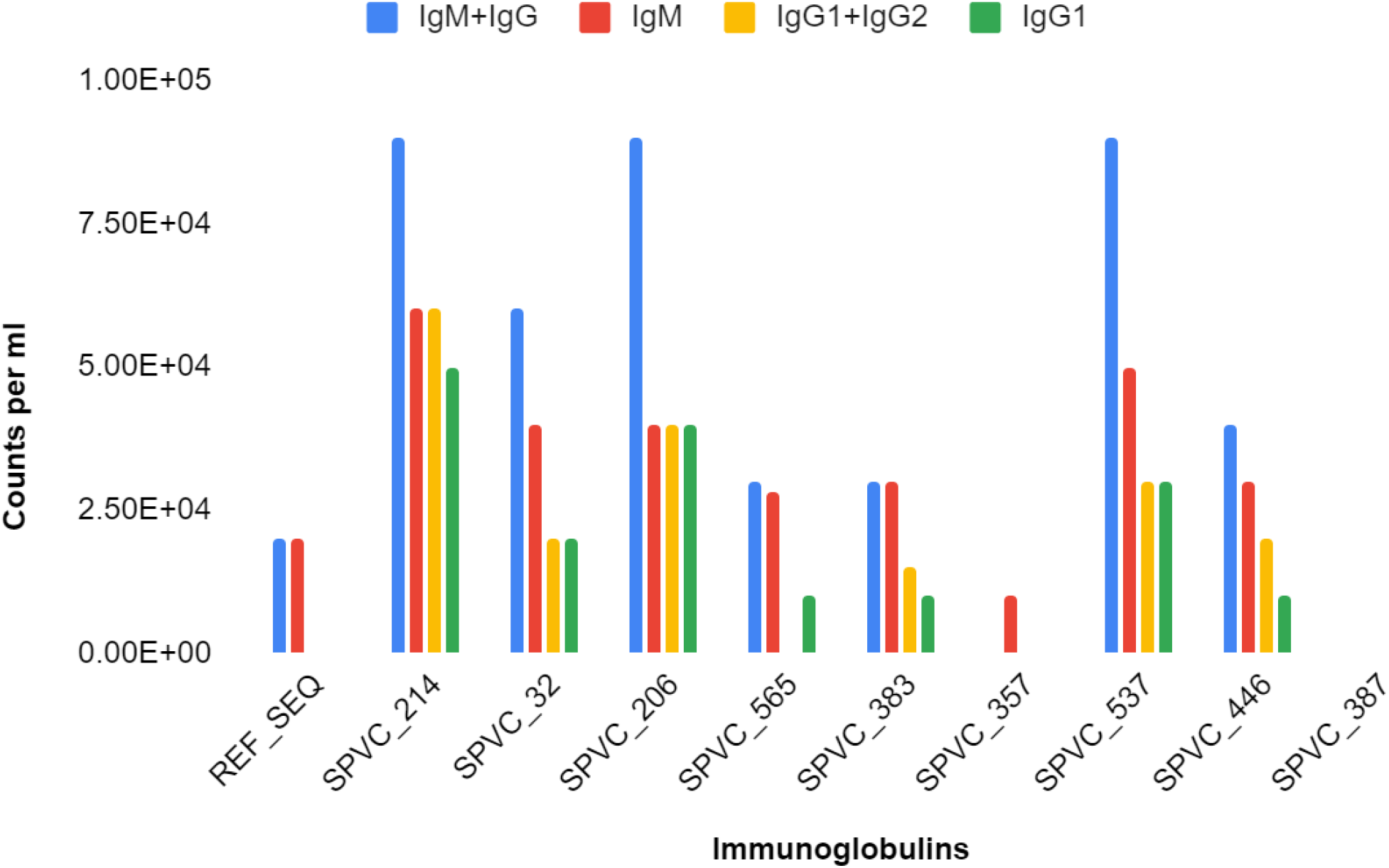
Immunoglobulin counts of MEBP variants (without adjuvants+HIS-tag) after 1st Dose (X axis-Vaccine constructs, Y axis - counts per ml)

**Figure 5b:**
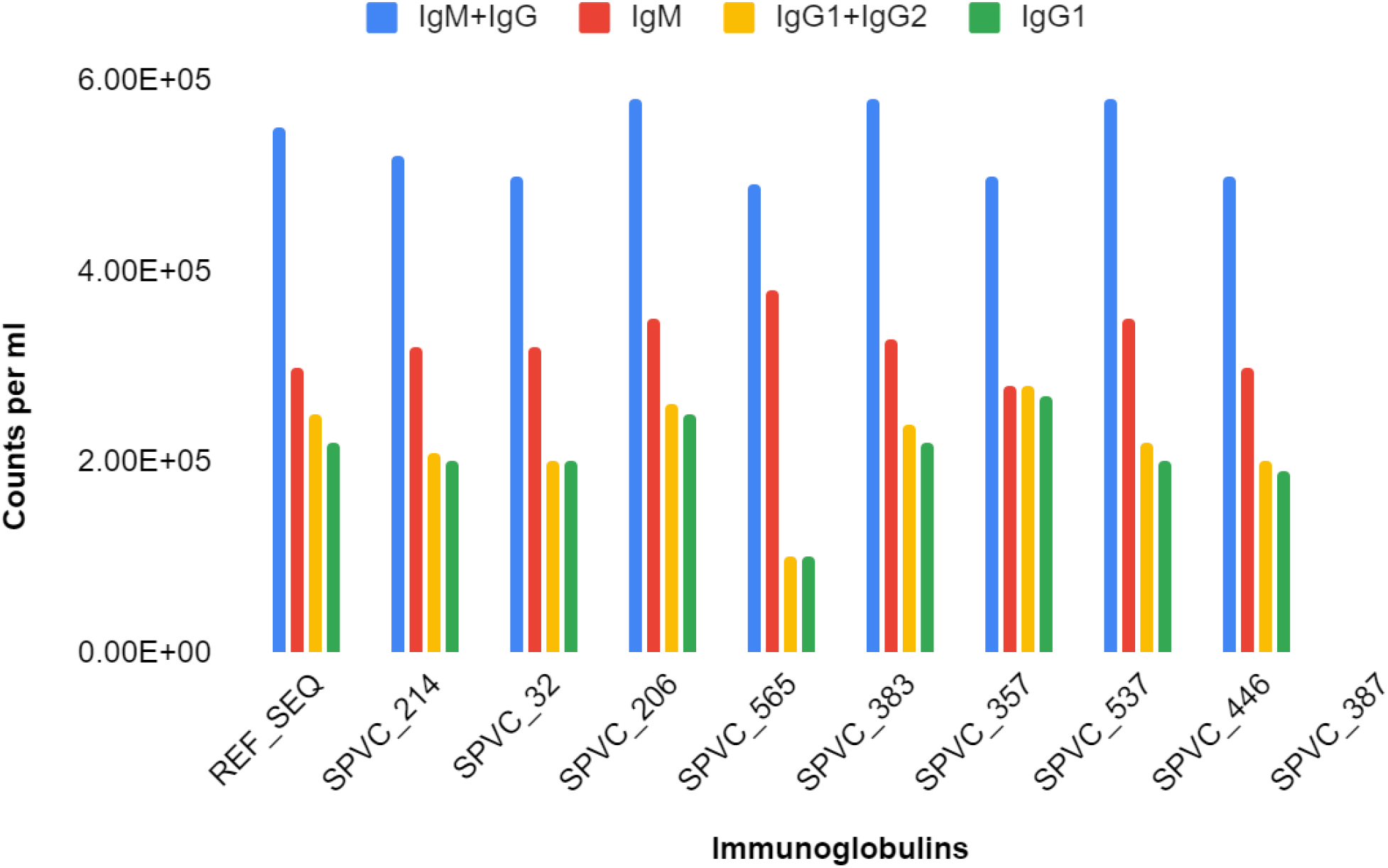
Immunoglobulin counts of MEBP variants (without adjuvants+HIS-tag) after 2nd Dose (X axis-Vaccine constructs, Y axis - counts per ml)

**Figure 5c:**
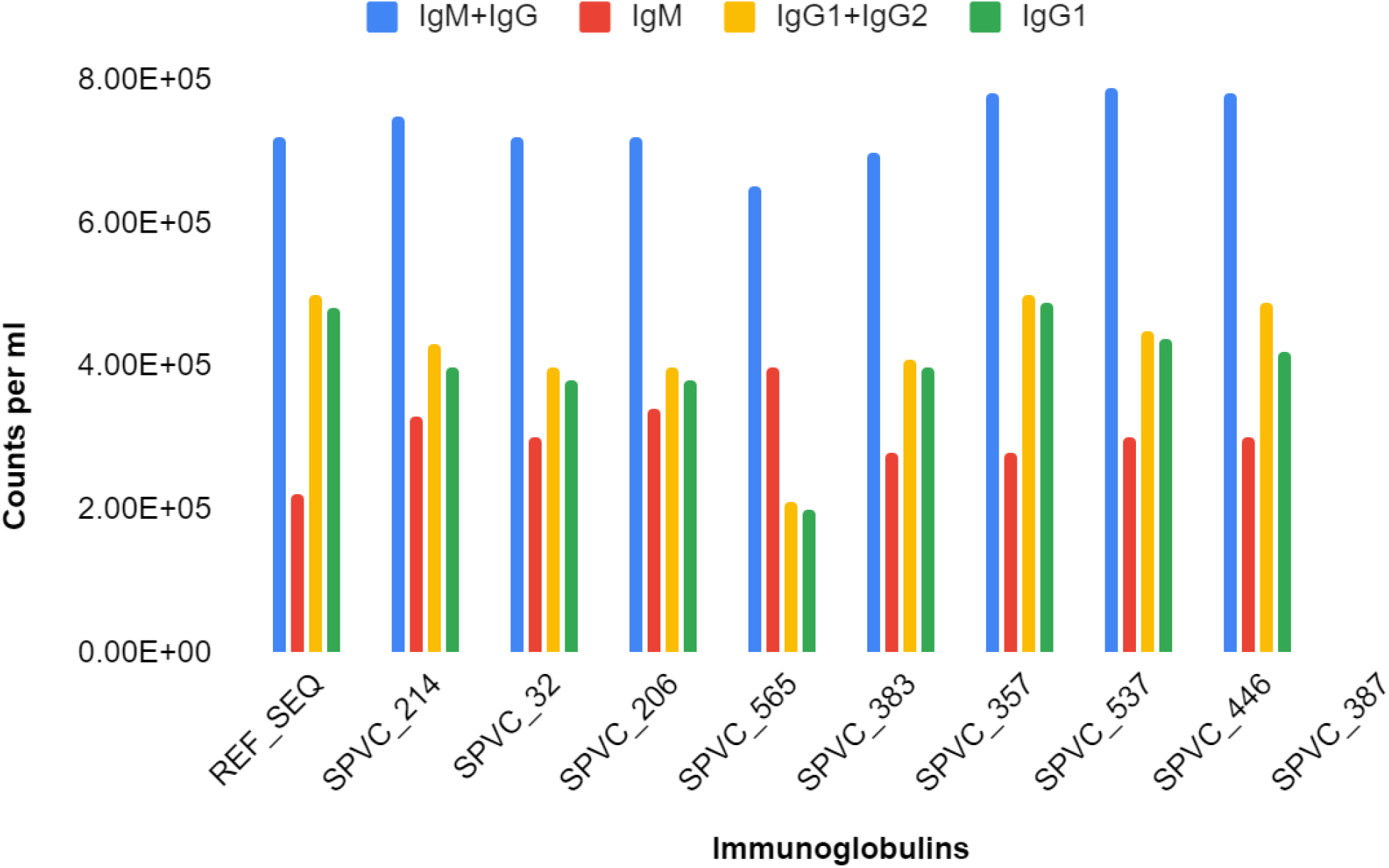
Immunoglobulin counts of MEBP variants (without adjuvants+HIS-tag) after 3rd Dose (X axis-Vaccine constructs, Y axis - counts per ml)

It is also observed that some variants such as SPVC_383 trigger high TH cell populations per state with counts reaching around 8200 per mm3 with an average antibody (IgG+IgM) response of around 570000 counts. When compared to others, the same variant also shows the best B cell population counts (800 per mm3).

In IFN-γ, it is interesting to note that variants with adjuvant+HIS-tag and without adjuvant+HIS-tag have totally different trends in bar graphs Fig. 6(a,b). For example, in (with adjuvant+HIS-tag) SPVC_446 triggers the highest concentration of IFN-γ and REF_SEQ has the lowest concentration of IFN-γ in the first dose.

**Figure 6a:**
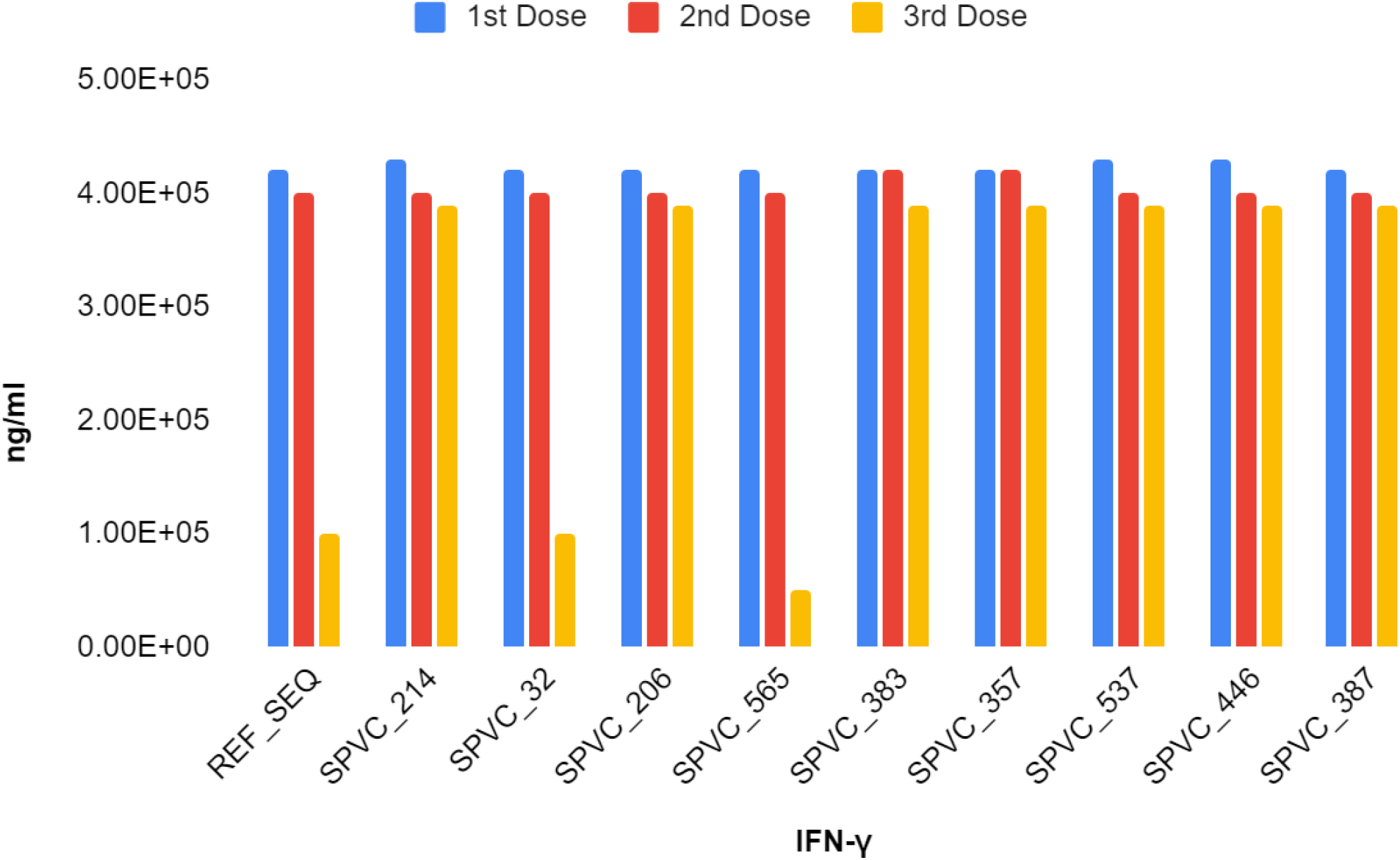
Concentration of IFN-γ for all MEBP variants (with adjuvant+HIS-tag) after the 1st, 2nd, and 3rd Doses. (X axis-Vaccine constructs, Y axis - ng/ml)

**Figure 6b:**
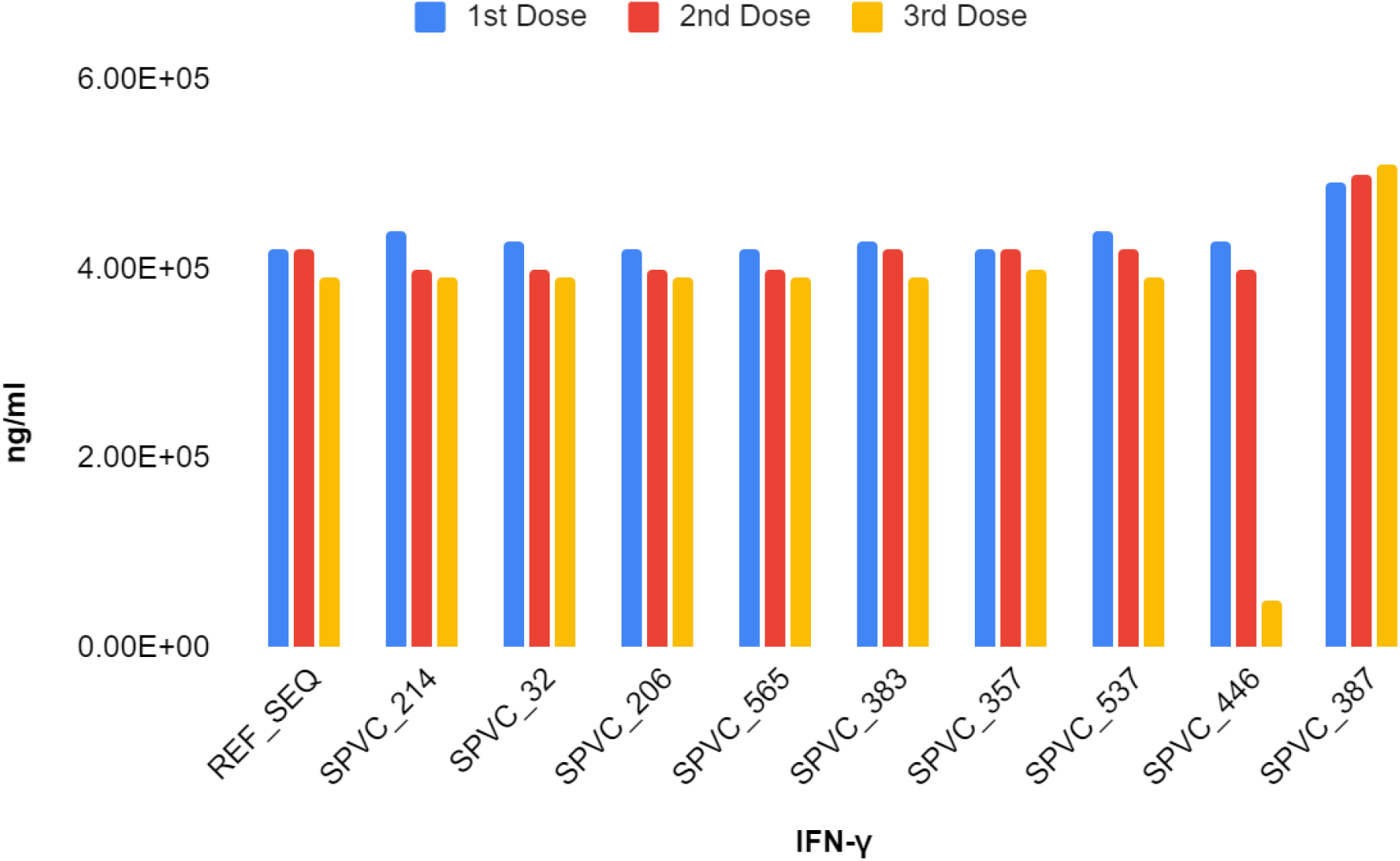
Concentration of IFN-γ for all MEBP variants (without adjuvant+HIS-tag) after the 1st, 2nd, and 3rd Doses(X axis-Vaccine constructs, Y axis - ng/ml).

The above observations clearly demonstrate that change in the epitope order in a MEBP vaccine candidate influences immunogenicity.

### 3.6. Ranking the ten variants and identifying the most potent MEBPVC

Table 7 summarizes the receptor specific scores and final MVP score for each variant. From the MVP score, SPVC_387 is predicted to be the most potent MEBPVC followed by SPVC_206. As can be seen, the least potent is REF_SEQ clearly proving that better and more potent MEBPVCs are possible by changing the epitope order and that epitope order influences immunogenicity. (Details about the normalized data is provided in Supp. 4)

**Table 7:** MVP of each variant is calculated by adding Receptor associated MVP (RMVP)s. RMVPs were calculated for the two receptors, TLR4 and TLR8 using the normalized values as described in the methods section..

| MEBP Vaccine Potency (MVP) |  |  |  |
| --- | --- | --- | --- |
| MEBP Variant ID | RMVP(TLR4) | RMVP(TLR8) | MVP (TLR4+TLR8) |
| SPVC_387 | 6.960 | 7.068 | 14.029 |
| SPVC_206 | 7.302 | 4.662 | 11.964 |
| SPVC_565 | 5.801 | 5.378 | 11.179 |
| SPVC_537 | 7.998 | 3.178 | 11.176 |
| SPVC_357 | 5.299 | 4.321 | 9.620 |
| SPVC_446 | 4.586 | 4.345 | 8.931 |
| SPVC_383 | 3.991 | 4.678 | 8.669 |
| SPVC_32 | 3.299 | 5.204 | 8.503 |
| SPVC_214 | 2.371 | 5.941 | 8.312 |
| REF_SEQ | 2.392 | 5.226 | 7.618 |

## 4. Discussion

Vaccine development typically takes 10 years. In the pre-COVID-19 world, the fastest vaccine development time recorded was four years against mumps. It is no small feat to develop a vaccine against COVID-19 in a span of 9 - 10 months and vaccinate nearly 1.5 Billion people. This shall be the new benchmark and reference for future vaccine development strategies and preparedness for future pandemics. This has become possible because of global cooperation for vaccine research and distribution[49].

The current COVID-19 vaccines listed under EUL have respective advantages and disadvantages[50]. The major disadvantage of Pfizer/BioNtech Comirnaty vaccine is its stringent cold chain requirement though it has shown very good titers[51]. The adenovector-based vaccines show relatively less effective neutralizing antibody response[52]. The inactivated vaccines seem to show inferior immunogenicity and low T Cell response, though have shown lower adverse reactions[53]. Similar to inactivated vaccines, the protein subunit vaccines show low immunogenicity.

However, the possible advantages and potential benefits attracted the pharma companies to invest in protein subunit platforms. More than 30% of the total COVID-19 vaccine candidates undergoing trials are protein subunit vaccines with 65% under preclinical trials. Peptide-based vaccines have many unique benefits such as 1. fully defined composition, 2. affordable large scale production 3. stable in storage and freeze dryable 4. absence of biological contamination 5. minimum allergic and autoimmune responses 6. customizable multipurpose therapeutic, and 7. standard rDNA technology-based production and manufacturing protocols in place. MEBPVC has further advantages of designing multi-epitope, multi-target, multi-copy, multi-disease, and custom-size (molecular weight) vaccine constructs. The MEBP subunit vaccine platforms are in the initial phases of development.

Applications of *in silico* approach to design a MEBP vaccine is one of the pragmatic opportunities that can reduce the time in developing vaccines and reach the market in shorter time duration. The *in silico* methodology presented in this article shall further reduce the time to identify potential new vaccine candidates under the protein subunit vaccine platform.

It is known that peptide vaccines are weakly immunogenic. Considering the advantages offered by peptide-based vaccines that include MEBP vaccines, it is worth addressing the peptide vaccine-specific issues, where the major issue seems to be lower immunogenicity. This limitation is being effectively addressed through a) combining with adjuvants such as b-defensin 2, HSP70, HBD-2, Matrix-M1, nanoparticles, b) altering the size (molecular weight), and others. Adjuvants have shown to significantly boost immunogenicity but have not matched the current platforms such as RNA, adenovirus vector, and inactivated virus-based platforms [54–56].

It is the fundamental phenomenon that changes in the amino acid order change the structure and function, giving the clue that the earlier reported MEBPVC (REF_SEQ) could have variants if the epitope order changed. A set of ten variants were generated manually to explore if the variants thus generated shall have altered immunogenicity.

The variants were analyzed at the sequence, structure, and complex interaction and dosage versus immune response level. In homology modeling, it is a common rule of thumb that for any two sequences, if the sequence identity is > 30%, it is assumed that their 3D structures shall be similar and likely to have identical function[57]. Further, it is also believed that with the increase in the sequence identity, the structural similarity also increases, i.e, RMSD decreases. However, it is interesting to note that the variants show deviations from the rule of thumb. As can be seen from Table 3 and Table 4, there are many pairs that show deviations. There have been studies that proved that 3D structures of 100% identical sequences were having natural conformations that have RMSDs as large as 24 Å[58]. There have been studies where an all-**α** helix protein(Protein G) was engineered and transformed into an all-**β** protein(Protein Rop) by changing only 50% of the amino acid composition [59–61]. There is a need to experimentally verify the MEBP variant 3D structures through experimental structure determination techniques such as X-Ray Crystallography, NMR and or Electron microscopy.

The analysis, indeed, strongly suggests that changing the epitope order in MEBPVC changes the structure and hence the various associated properties resulting in the alteration of immunogenicity of the variant. Hence more potent MEBPVCs can be identified if the protocol described in this article is followed. The step to generate a dataset of shuffled variants is key in the analysis as this step enables the comparative study which otherwise has not been reported till now in MEBP vaccine design protocols. MVP score has also been developed which provides an opportunity to rank and identify the most potent MEBPVC from the dataset. Further, the data generated becomes the necessary input for developing better scoring schemes and algorithms.

It is pertinent to mention that the number of variants was restricted to ten out of a possible 3,628,800 (10! possibilities) unique variants of similar length. The peptide length was restricted to 183 amino acids. The scope of the work was to test if the immunogenicity changes with the epitope order in the MEBPVC. For some parameters, it was also observed that the changes in the biophysical and immune parameters were not significant. This can be argued that the small sample size, restriction on the length of the construct, and manual shuffling of epitopes could be the reasons for such small changes. The next challenges are to work with a bigger dataset of variants, optimize the parameters influencing the MEBP vaccine design and gain deeper insights into the mechanisms behind the mutations and their virulence and improvise the MVP score incorporating the knowledge and the decision making systems such as AI and ML[62].

## 5. Conclusions

In pursuit to discover more potent MEBPVCs, given the list of predicted epitopes, a methodology to generate variants and three predictors percent epitope accessibility (PEA), receptor-specific vaccine potency (RMVP) and MEBP vaccine potency (MVP) scores have been introduced to quantify and enable opportunity for the development of efficient MEBP vaccine design platform. This enabled not only ranking but also picking up the best MEBPVC. Thus, the results prove that the reported MEBPVC (REF_SEQ) is not the most potent candidate after all[22]. In this article, only the *order* of epitopes has been used for generating variants. However, other parameters such as length of epitopes, length of the sequence, copy numbers, and or other parameters should also be explored in the future to design and discover more potent MEBPVCs. In conclusion, our *in silico* analysis and results indeed prove that changes in the position or order of the epitopes change the properties of the MEBPVC. Henceforth, the dataset generation method, PEA, RMVP, and MVP score should enable generating novel MEBPVCs and may be adopted in all MEBPVC design pursuits. The method enables the discovery of the computationally validated most immunogenic MEBPVC, each having a unique epitope order. Experimental validation and verification has no substitute hence, the computationally validated vaccine constructs, with IDs, SPVC_387 and SPVC_206, need to be validated through *in vitro*, and *in vivo* experimental studies. The experimental validation provides important insights and inputs for improvising and developing a more efficient and more reliable scoring function and the improved versions of the software.

## Supplementary Materials

Supplementary Material 1: MEBP variants in FASTA Format; Supplementary Material 2: Figure S1 to 20(Simulation analysis of MEBP Variants); Supplementary Material 3: Figure S1 to 20(Immune response simulation results of MEBP Variants); Supplementary Material 4: Normalized data for MEBP variants

## Author Contributions

**n**Conceptualization: BVLS. Investigation: BVLS, SMR. Methodology: BVLS, SMR, KPS. Software: SMR, KPS. Supervision: BVLS. Validation: BVLS. Writing – original draft: BVLS. Writing – review & editing: BVLS, SMR

## Institutional Review Board Statement

Not applicable

## Informed Consent Statement

Not applicable

## Data Availability Statement

Data is contained within the article.

## Acknowledgments

Burra V L S Prasad acknowledges DST-NSM (DST/NSM/R&D_HPC _Applications/Sanction/2021/16) ICMR (ISRM/12(7)/2019) and SERB (CRG/2018/003276) for supporting the COVID-19 Vaccine development initiative.

