## Supplementary Material 1 for "Epitope order Matters in multi-epitope-based peptide (MEBP) vaccine design: An *in silico* study"

1) **MEBP Variants in FASTA format**

### >REF_SEQ

### GIINTLQKYYCRVRGGRCAVLSCLPKEEQIGKCSTRGRKCCRRKKEAAAKTLDSKTQSLAAYGKQGNFKNLAAYCYGV

### SPTKLAAYKIADYNYKLAAYVVVLSFELLGPGPGIGINITRFQGPGPGYGFQPTNGVGPGPGVLSFELLHAGPGPGLQ

### IPFAMQMGPGPGIAIVMVTIMHHHHHH

### >SPVC_206

### GIINTLQKYYCRVRGGRCAVLSCLPKEEQIGKCSTRGRKCCRRKKEAAAKYGFQPTNGVGPGPGGKQGNFKNLAAYVL

### SFELLHAGPGPGIGINITRFQGPGPGKIADYNYKLAAYTLDSKTQSLAAYCYGVSPTKLAAYIAIVMVTIMGPGPGVV

### VLSFELLGPGPGLQIPFAMQMHHHHHH

### >SPVC_214

### GIINTLQKYYCRVRGGRCAVLSCLPKEEQIGKCSTRGRKCCRRKKEAAAKVLSFELLHAGPGPGGKQGNFKNLAAYCY

### GVSPTKLAAYKIADYNYKLAAYTLDSKTQSLAAYYGFQPTNGVGPGPGVVVLSFELLGPGPGIAIVMVTIMGPGPGIG

### INITRFQGPGPGLQIPFAMQMHHHHHH

### >SPVC_32

### GIINTLQKYYCRVRGGRCAVLSCLPKEEQIGKCSTRGRKCCRRKKEAAAKCYGVSPTKLAAYKIADYNYKLAAYTLDS

### KTQSLAAYYGFQPTNGVGPGPGGKQGNFKNLAAYIGINITRFQGPGPGVVVLSFELLGPGPGLQIPFAMQMGPGPGIA

### IVMVTIMGPGPGVLSFELLHAHHHHHH

### >SPVC_357

### GIINTLQKYYCRVRGGRCAVLSCLPKEEQIGKCSTRGRKCCRRKKEAAAKTLDSKTQSLAAYYGFQPTNGVGPGPGCY

### GVSPTKLAAYVVVLSFELLGPGPGIAIVMVTIMGPGPGIGINITRFQGPGPGKIADYNYKLAAYGKQGNFKNLAAYVL

### SFELLHAGPGPGLQIPFAMQMHHHHHH

### >SPVC_537

### GIINTLQKYYCRVRGGRCAVLSCLPKEEQIGKCSTRGRKCCRRKKEAAAKCYGVSPTKLAAYKIADYNYKLAAYGKQG

### NFKNLAAYYGFQPTNGVGPGPGVVVLSFELLGPGPGLQIPFAMQMGPGPGIAIVMVTIMGPGPGTLDSKTQSLAAYIG

### INITRFQGPGPGVLSFELLHAHHHHHH

### >SPVC_383

### GIINTLQKYYCRVRGGRCAVLSCLPKEEQIGKCSTRGRKCCRRKKEAAAKKIADYNYKLAAYGKQGNFKNLAAYVVVL

### SFELLGPGPGTLDSKTQSLAAYCYGVSPTKLAAYYGFQPTNGVGPGPGVLSFELLHAGPGPGIAIVMVTIMGPGPGIG

### INITRFQGPGPGLQIPFAMQMHHHHHH

### >SPVC_565

### GIINTLQKYYCRVRGGRCAVLSCLPKEEQIGKCSTRGRKCCRRKKEAAAKCYGVSPTKLAAYKIADYNYKLAAYYGFQ

### PTNGVGPGPGVVVLSFELLGPGPGTLDSKTQSLAAYIGINITRFQGPGPGIAIVMVTIMGPGPGLQIPFAMQMGPGPG

### GKQGNFKNLAAYVLSFELLHAHHHHHH

### >SPVC_446

### GIINTLQKYYCRVRGGRCAVLSCLPKEEQIGKCSTRGRKCCRRKKEAAAKYGFQPTNGVGPGPGVVVLSFELLGPGPG

### GKQGNFKNLAAYIAIVMVTIMGPGPGCYGVSPTKLAAYTLDSKTQSLAAYLQIPFAMQMGPGPGVLSFELLHAGPGPG

### KIADYNYKLAAYIGINITRFQHHHHHH

### >SPVC_387

### GIINTLQKYYCRVRGGRCAVLSCLPKEEQIGKCSTRGRKCCRRKKEAAAKGKQGNFKNLAAYVLSFELLHAGPGPGIG

### INITRFQGPGPGVVVLSFELLGPGPGCYGVSPTKLAAYTLDSKTQSLAAYKIADYNYKLAAYYGFQPTNGVGPGPGLQ

### IPFAMQMGPGPGIAIVMVTIMHHHHHH
