## Supplementary Material 2 for "Epitope order Matters in multi-epitope-based peptide (MEBP) vaccine design: An *in silico* study"

### 1) **Simulation analysis of MEBP Variants**

**A**	**B**

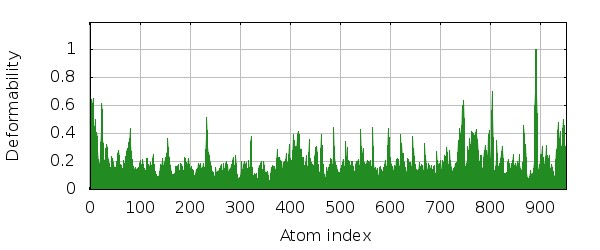

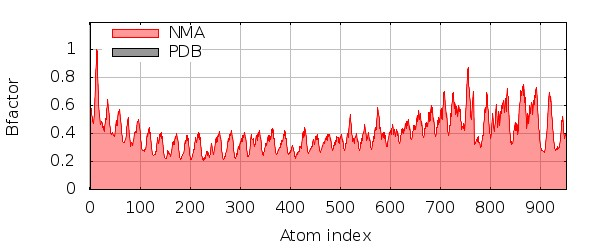

**C**

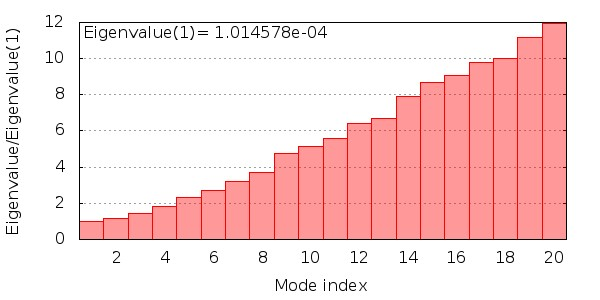

**D	E**

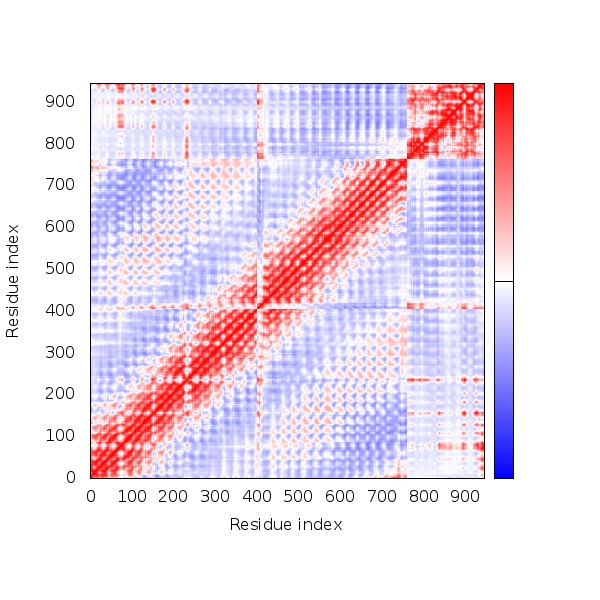

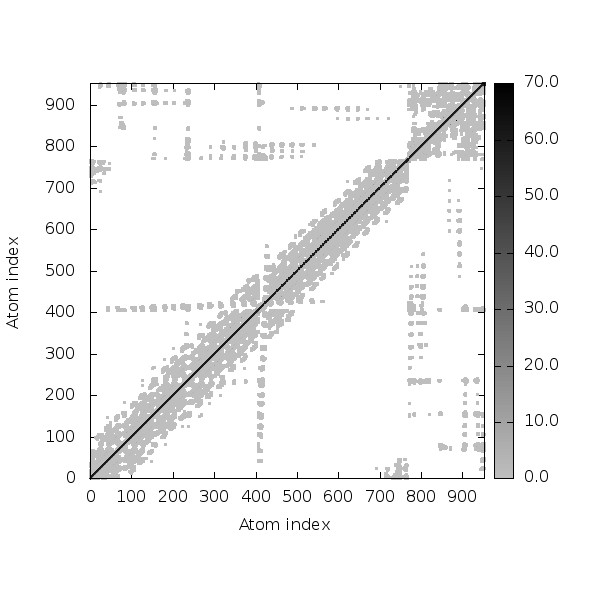

**Figure S1: SPVC_32(TLR8) simulation results [A) Deformability plot, B) B-factor plot, C) Eigenvalue plot , D) Covariance map [correlated (red), uncorrelated (white) or anti-correlated (blue) motions] , E) Elastic network (darker gray regions indicate more stiffer regions) of the complex]**

**A	B**

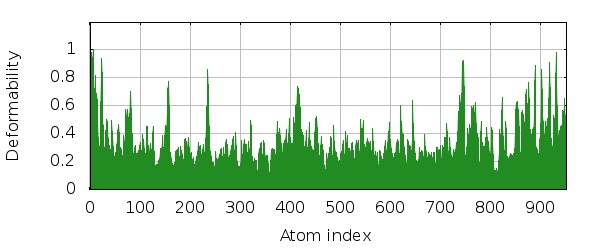

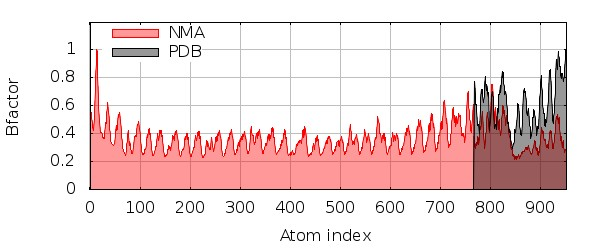

**C**

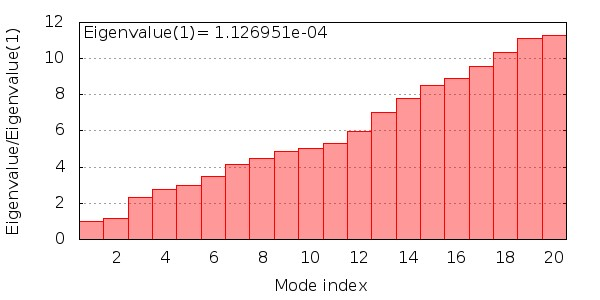

**D	E**

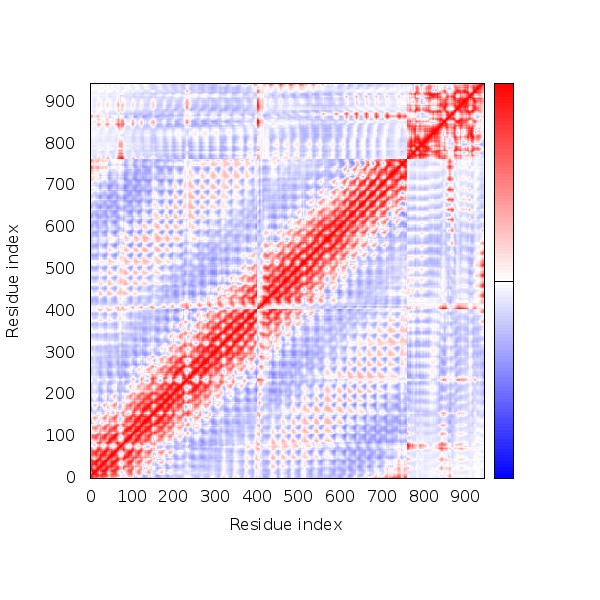

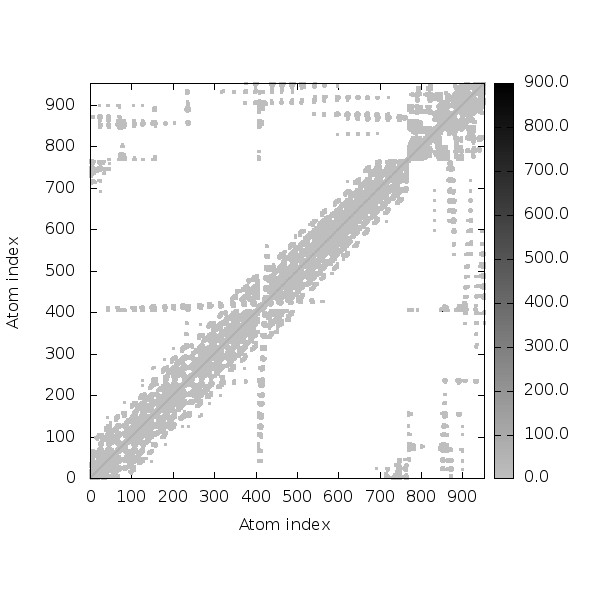
**Figure S2: SPVC_206(TLR8) simulation results [A) Deformability plot, B) B-factor plot, C) Eigenvalue plot , D) Covariance map [correlated (red), uncorrelated (white) or anti-correlated (blue) motions] , E) Elastic network (darker gray regions indicate more stiffer regions) of the complex]**

**A	B**

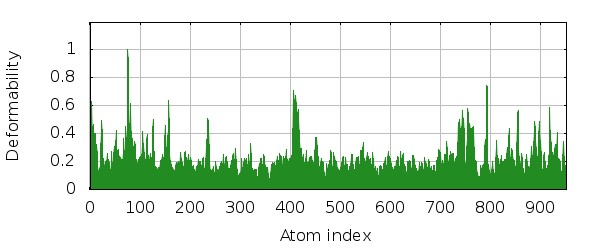

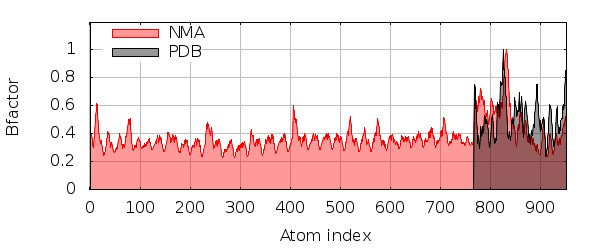

**C**

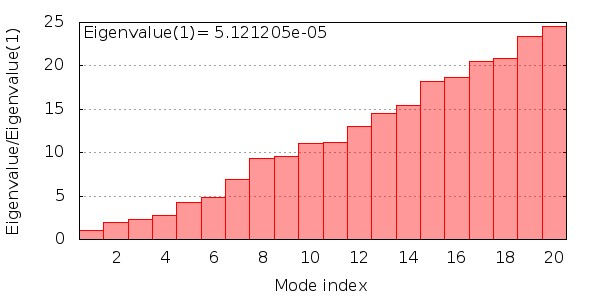

**D	E**

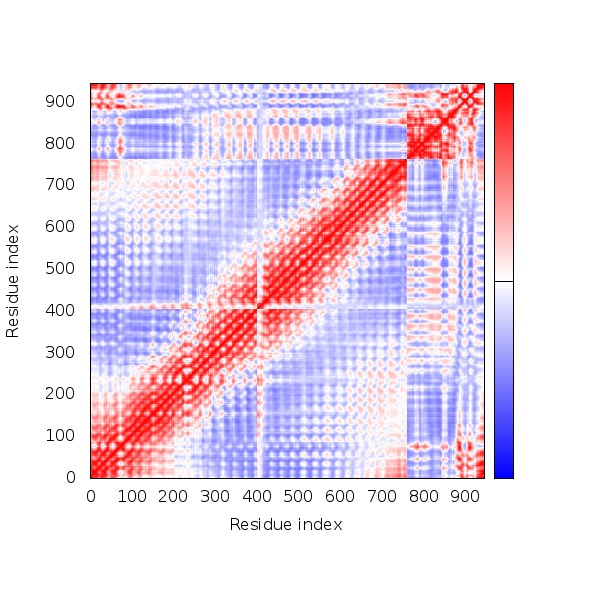

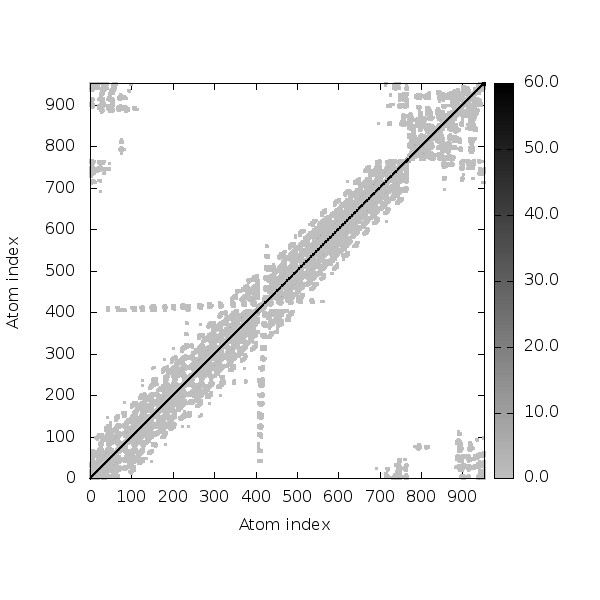

**Figure S3: SPVC_214(TLR8) simulation results [A) Deformability plot, B) B-factor plot, C) Eigenvalue plot , D) Covariance map [correlated (red), uncorrelated (white) or anti-correlated (blue) motions] , E) Elastic network (darker gray regions indicate more stiffer regions) of the complex]**

**A	B**

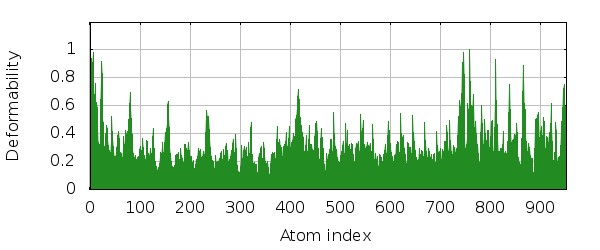

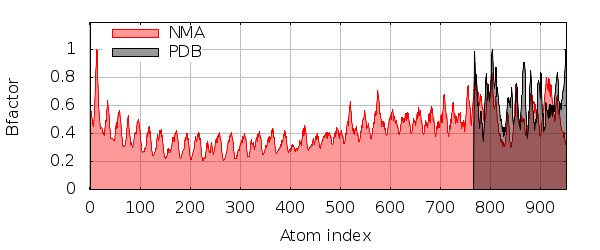

**C**

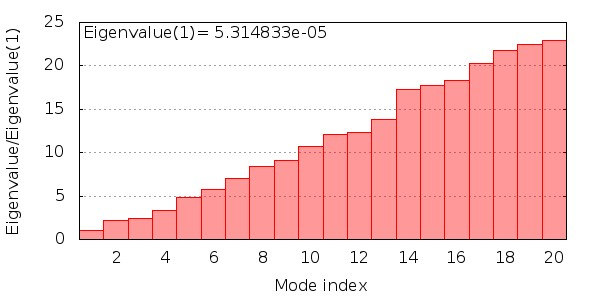

**D	E**

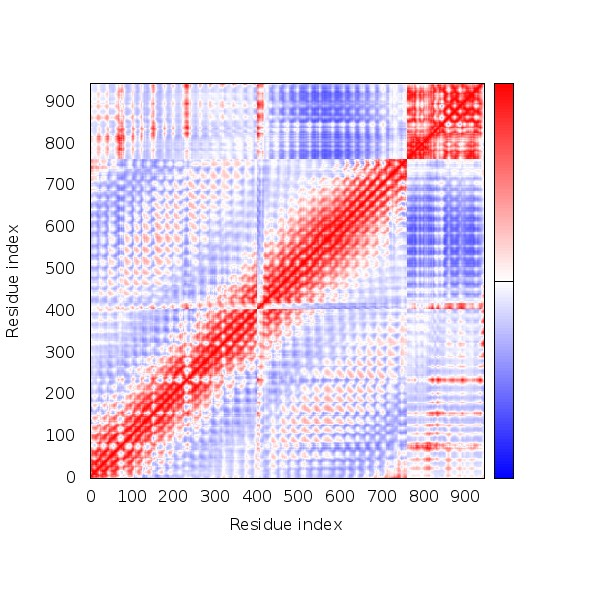

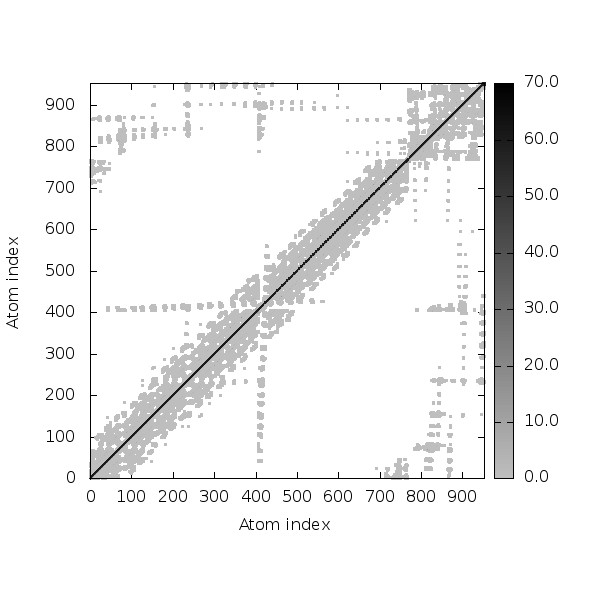

**Figure S4: SPVC_357(TLR8) simulation results [A) Deformability plot, B) B-factor plot, C) Eigenvalue plot , D) Covariance map [correlated (red), uncorrelated (white) or anti-correlated (blue) motions] , E) Elastic network (darker gray regions indicate more stiffer regions) of the complex]**

**A	B**

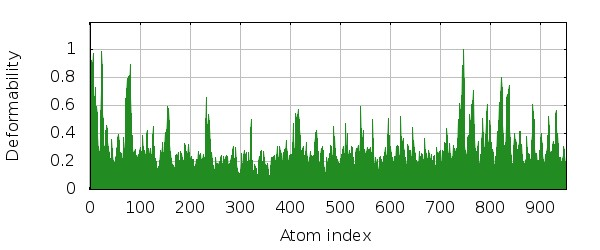

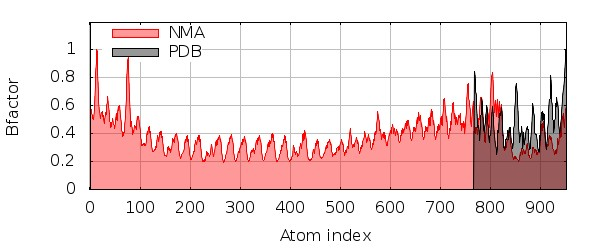

**C**

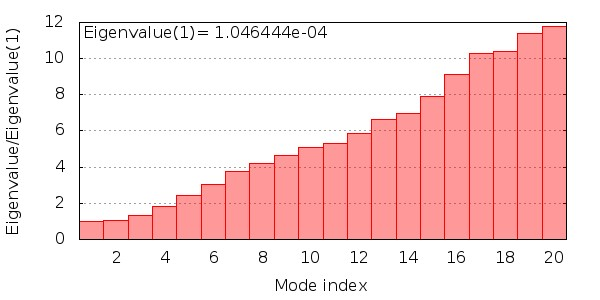

**D	E**

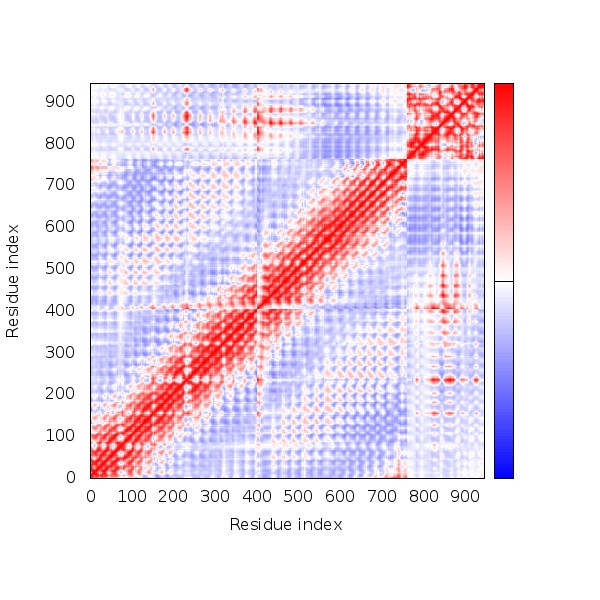

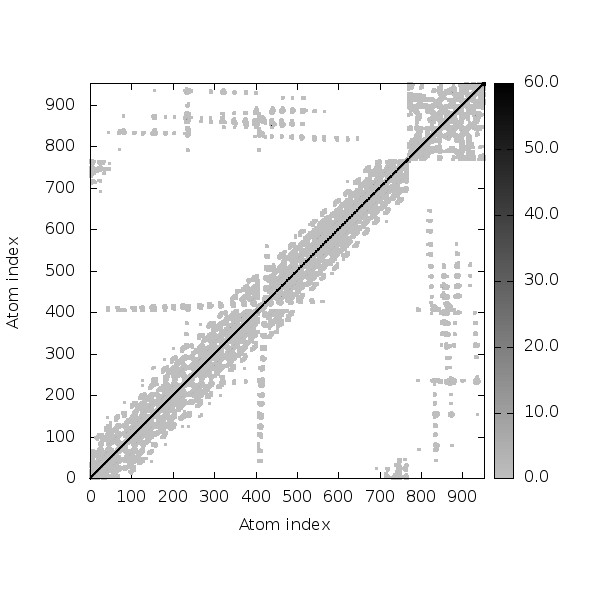

**Figure S5: SPVC_383(TLR8) simulation results [A) Deformability plot, B) B-factor plot, C) Eigenvalue plot , D) Covariance map [correlated (red), uncorrelated (white) or anti-correlated (blue) motions] , E) Elastic network (darker gray regions indicate more stiffer regions) of the complex]**

**A	B**

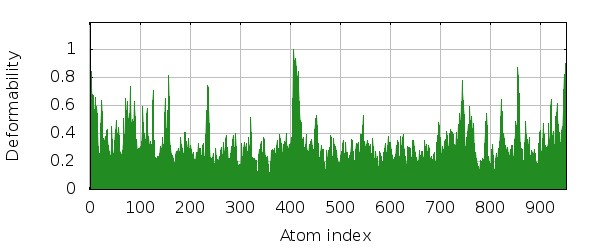

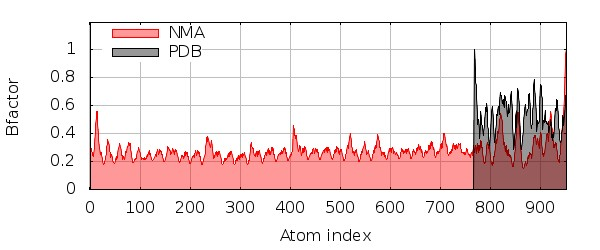

**C**

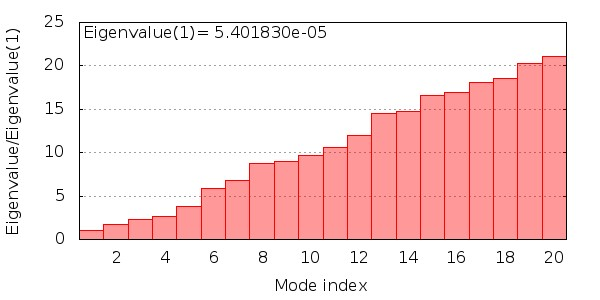

**D	E**

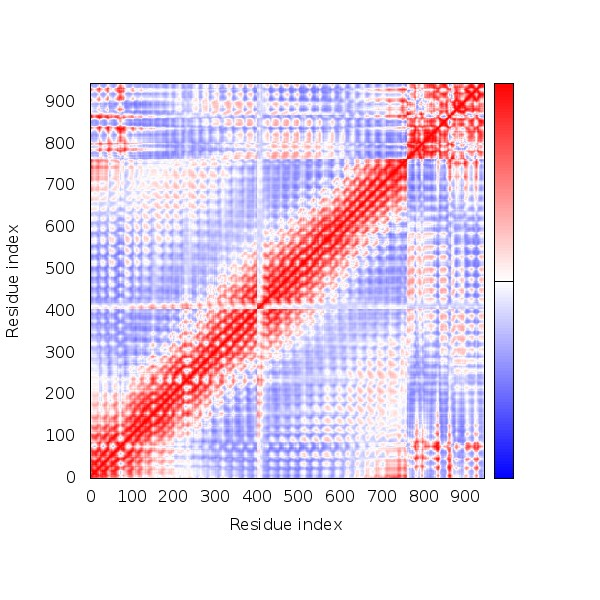

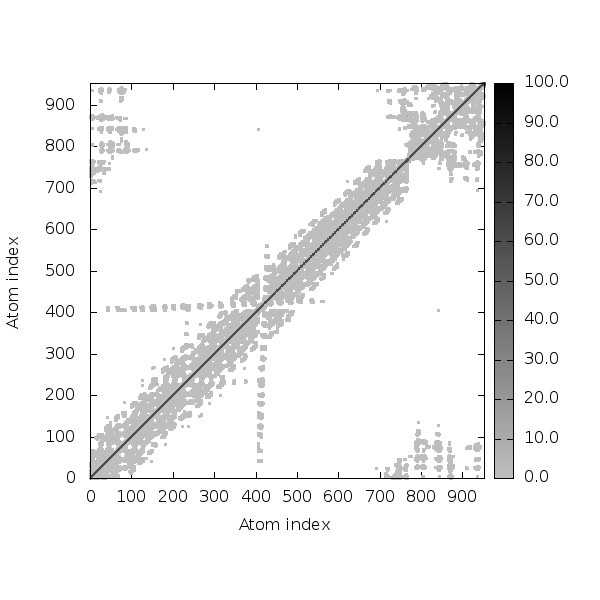

**Figure S6: SPVC_387(TLR8) simulation results [A) Deformability plot, B) B-factor plot, C) Eigenvalue plot , D) Covariance map [correlated (red), uncorrelated (white) or anti-correlated (blue) motions] , E) Elastic network (darker gray regions indicate more stiffer regions) of the complex]**

**A	B**

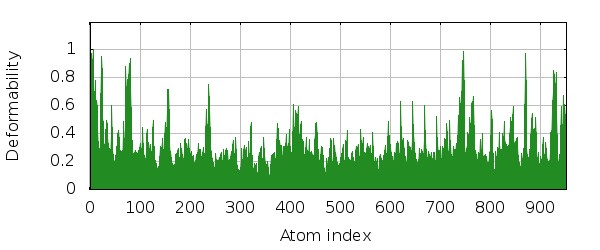

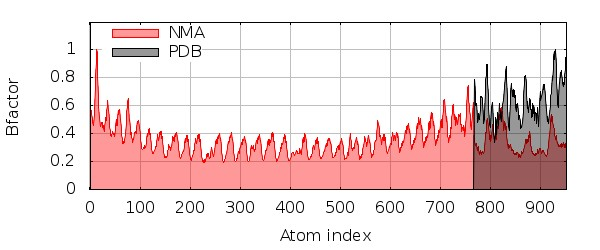

**C**

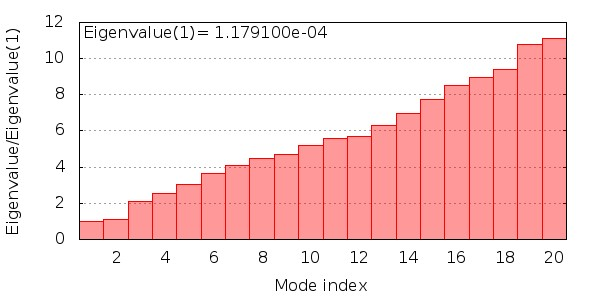

**D	E**

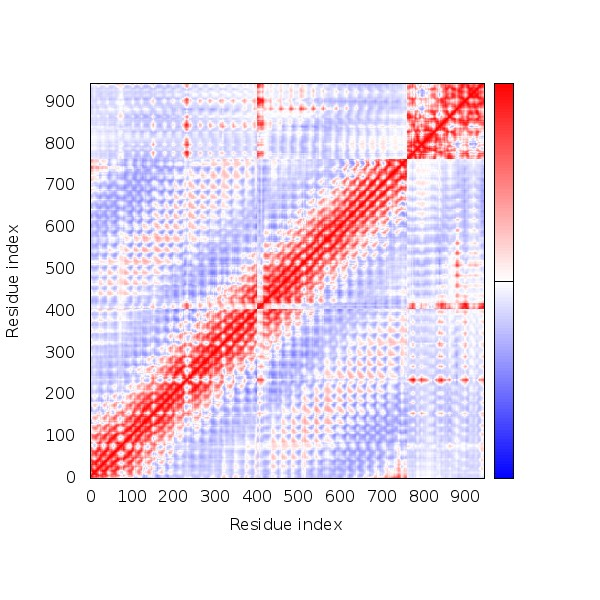

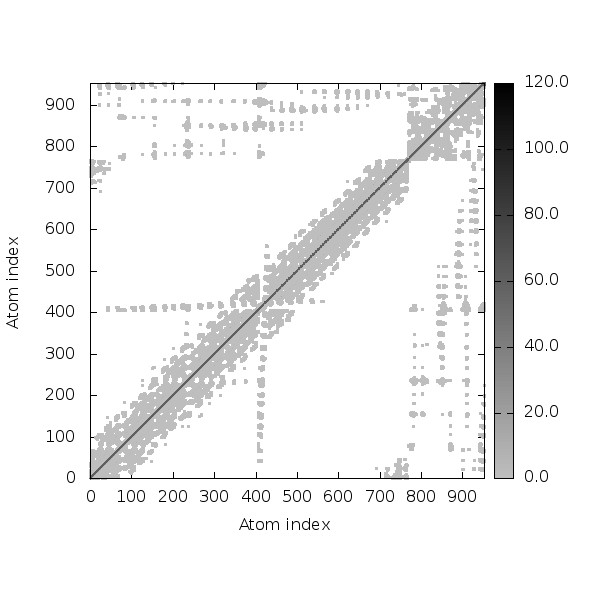

**Figure S7: SPVC_446(TLR8) simulation results [A) Deformability plot, B) B-factor plot, C) Eigenvalue plot , D) Covariance map [correlated (red), uncorrelated (white) or anti-correlated (blue) motions] , E) Elastic network (darker gray regions indicate more stiffer regions) of the complex]**

**A	B**

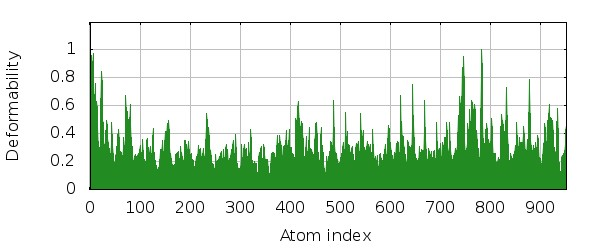

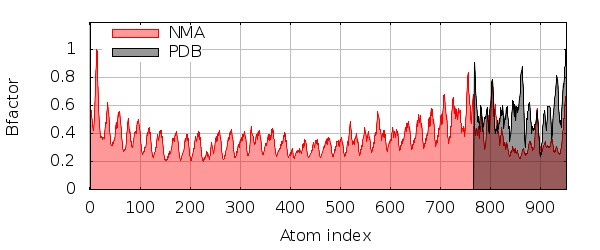

**C**

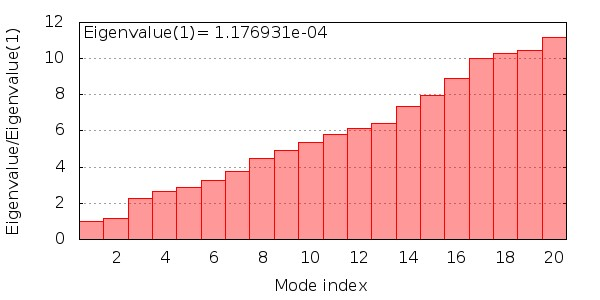

**D	E**

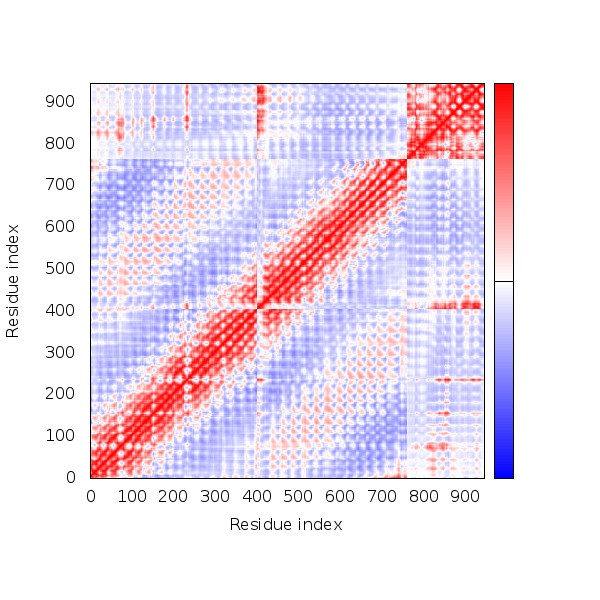

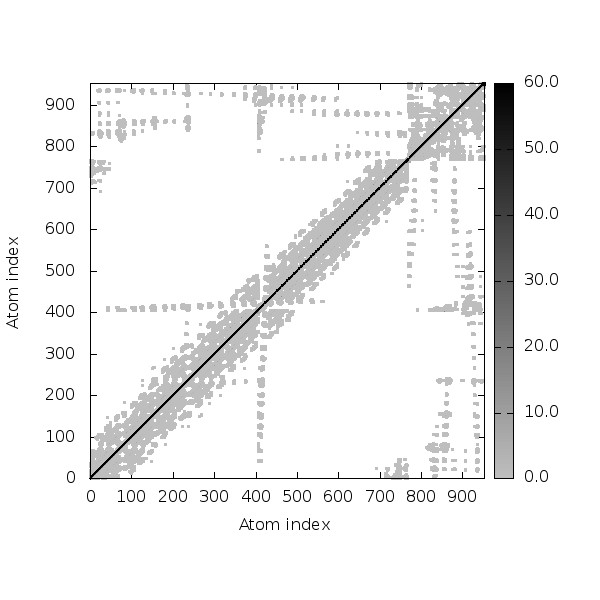

**Figure S8: SPVC_537(TLR8) simulation results [A) Deformability plot, B) B-factor plot, C) Eigenvalue plot , D) Covariance map [correlated (red), uncorrelated (white) or anti-correlated (blue) motions] , E) Elastic network (darker gray regions indicate more stiffer regions) of the complex]**

**A	B**

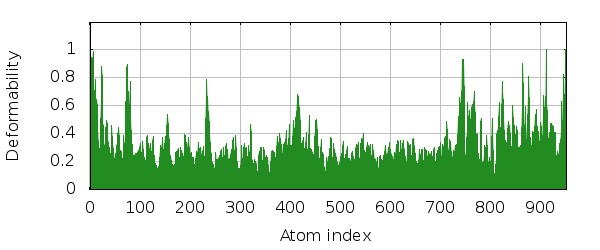

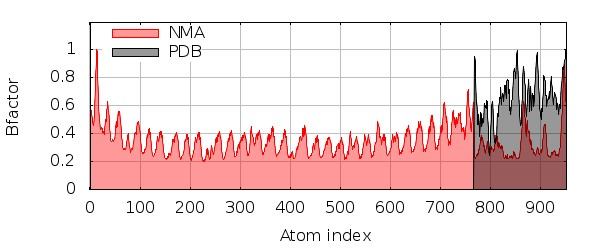

**C**

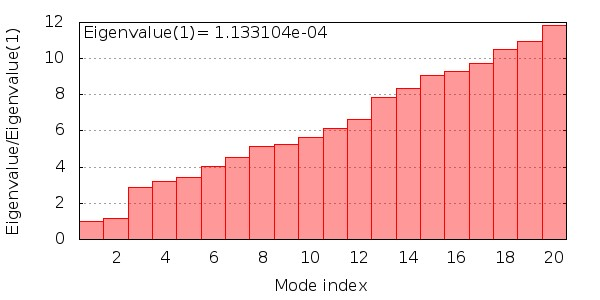

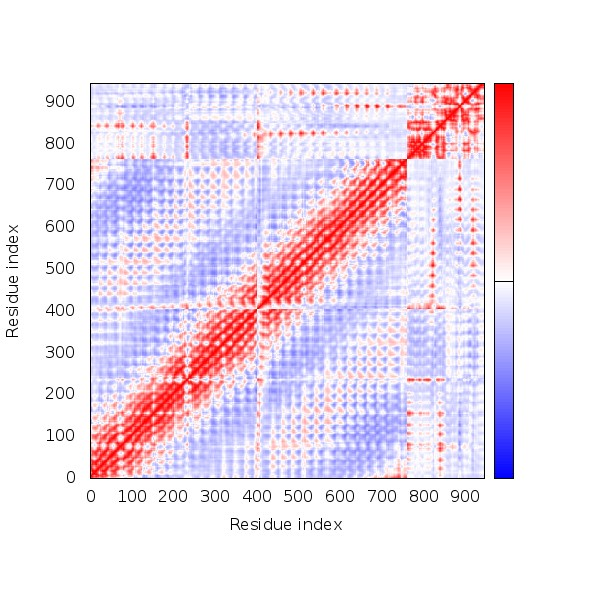

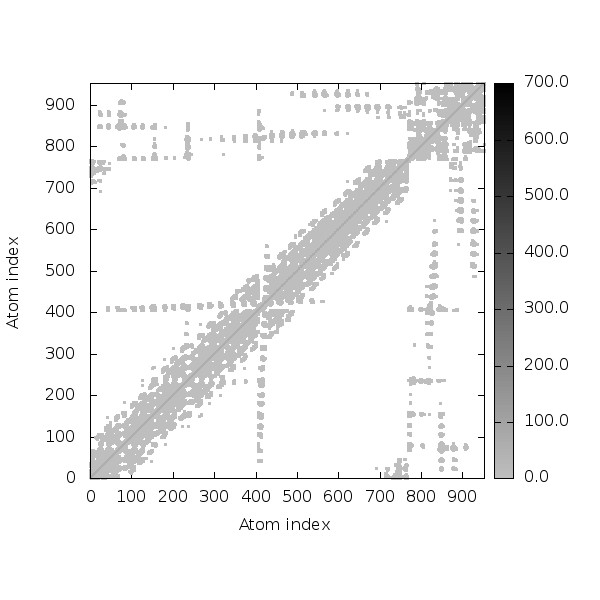

**Figure S9: SPVC_565(TLR8) simulation results [A) Deformability plot, B) B-factor plot, C) Eigenvalue plot , D) Covariance map [correlated (red), uncorrelated (white) or anti-correlated (blue) motions] , E) Elastic network (darker gray regions indicate more stiffer regions) of the complex]**

**A	B**

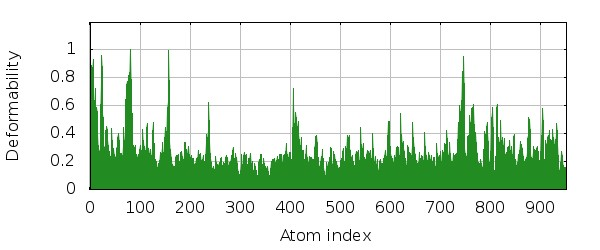

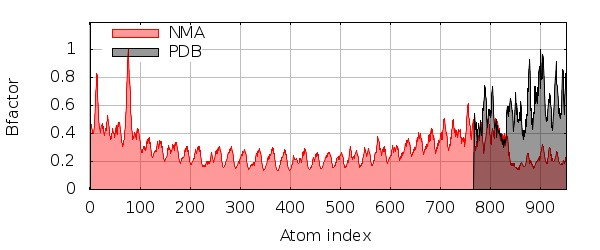

**C**

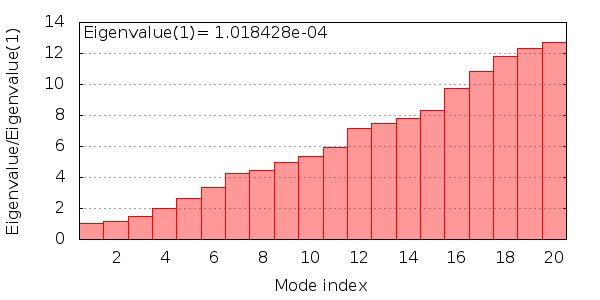

**D	E**

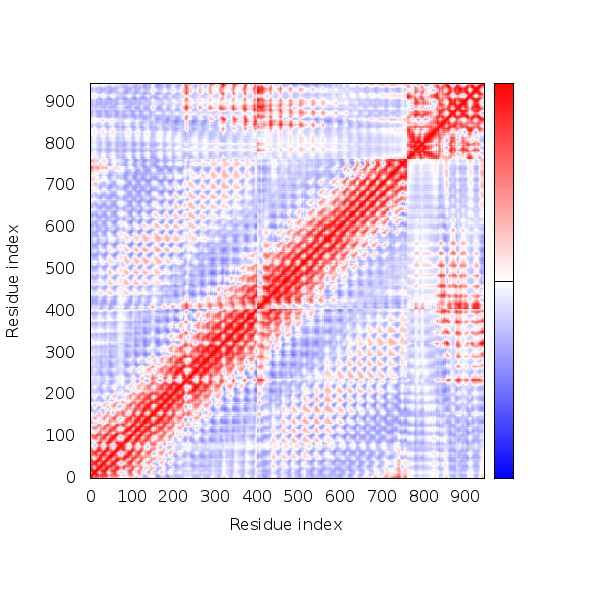

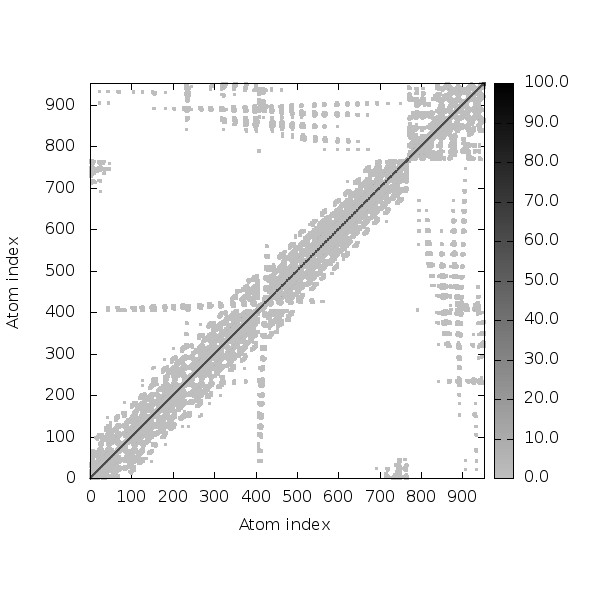

**Figure S10: REF_SEQ(TLR8) simulation results [A) Deformability plot, B) B-factor plot, C) Eigenvalue plot , D) Covariance map [correlated (red), uncorrelated (white) or anti-correlated (blue) motions] , E) Elastic network (darker gray regions indicate more stiffer regions) of the complex]**

**A	B**

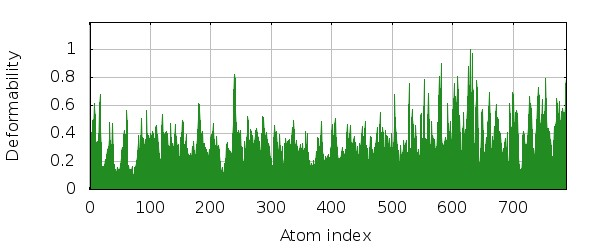

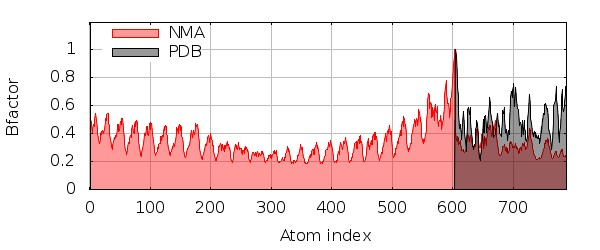

**C**

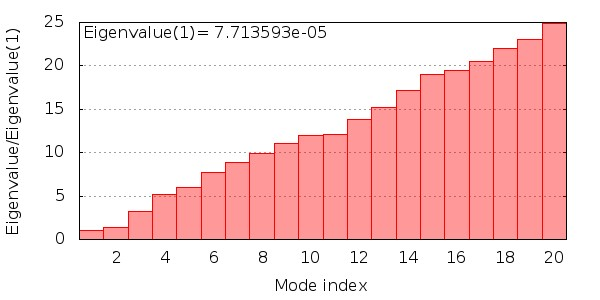

**D	E**

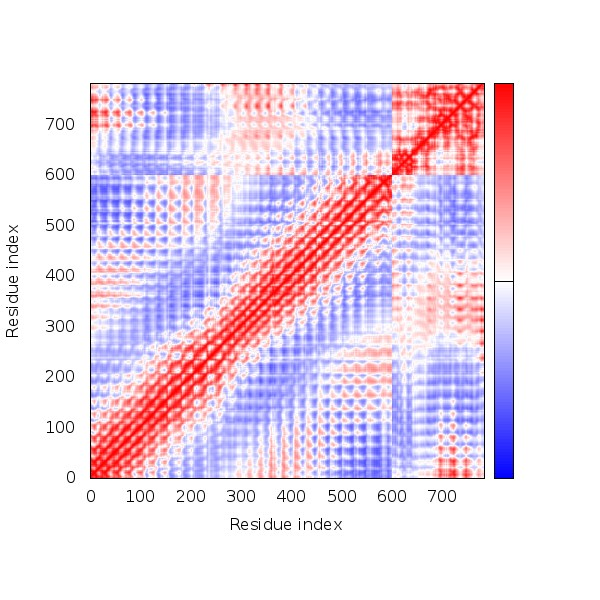

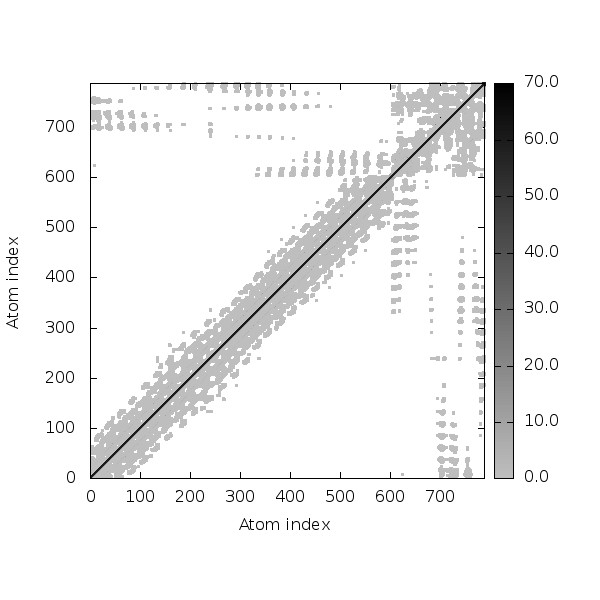

**Figure S11: SPVC_32(TLR4) simulation results [A) Deformability plot, B) B-factor plot, C) Eigenvalue plot , D) Covariance map [correlated (red), uncorrelated (white) or anti-correlated (blue) motions] , E) Elastic network (darker gray regions indicate more stiffer regions) of the complex]**

**A	B**

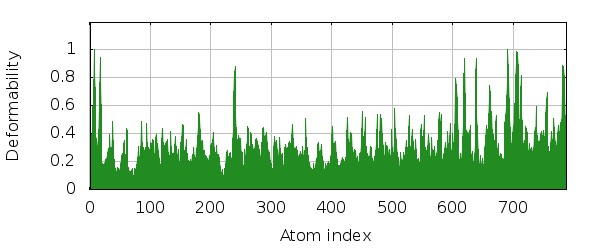

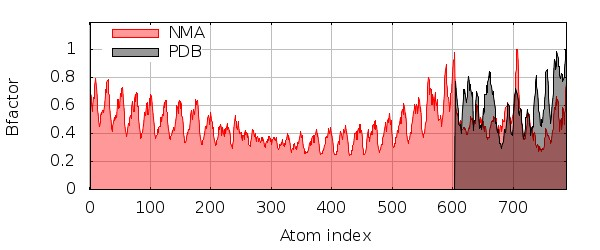

**C**

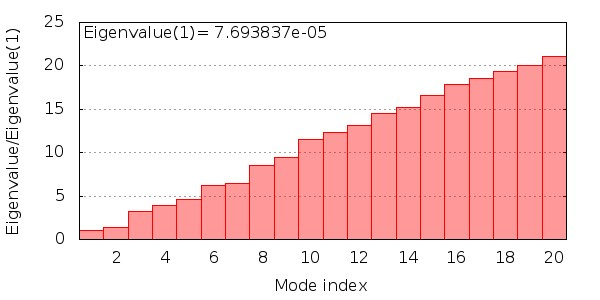

**D	E**

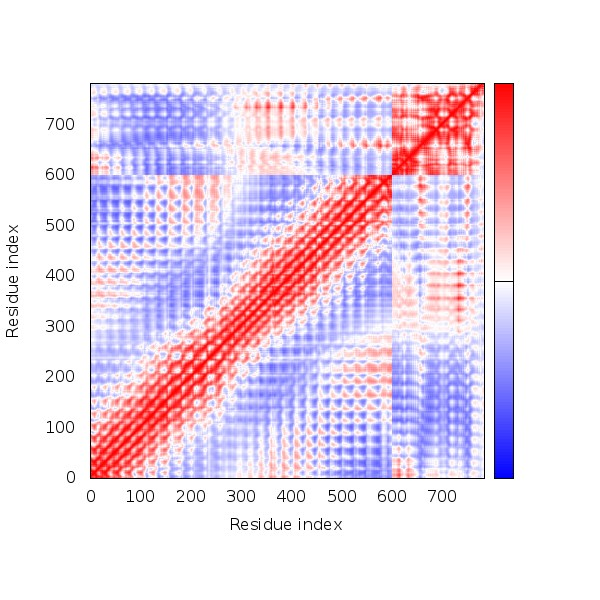

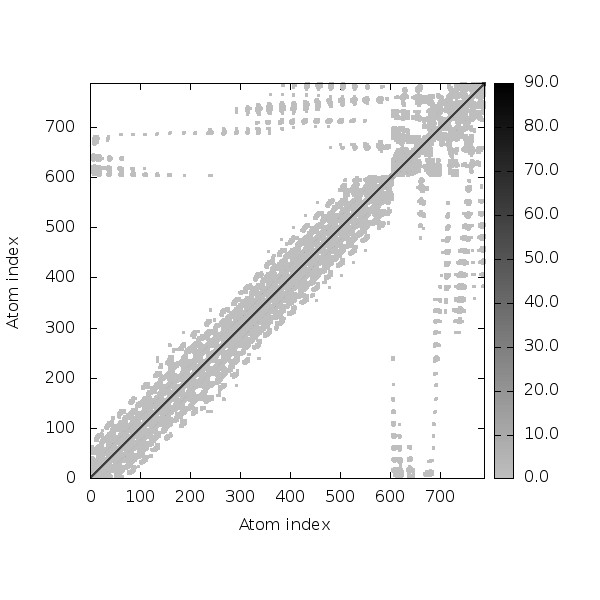

**Figure S12: SPVC_206(TLR4) simulation results [A) Deformability plot, B) B-factor plot, C) Eigenvalue plot , D) Covariance map [correlated (red), uncorrelated (white) or anti-correlated (blue) motions] , E) Elastic network (darker gray regions indicate more stiffer regions) of the complex]**

**A	B**

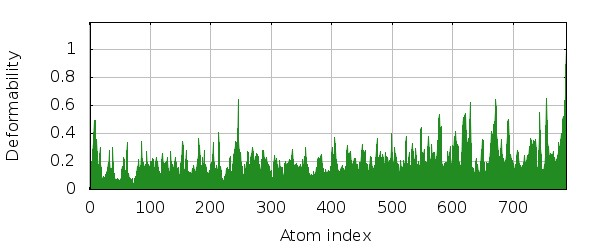

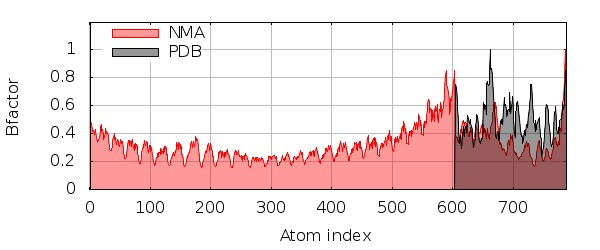

**C**

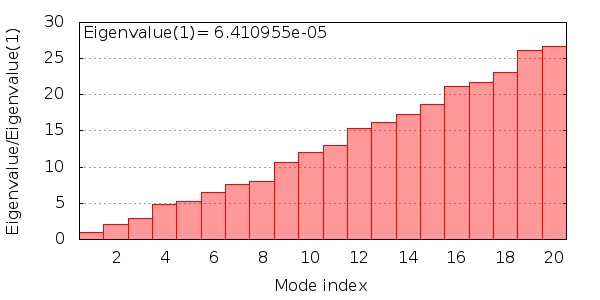

**D	E**

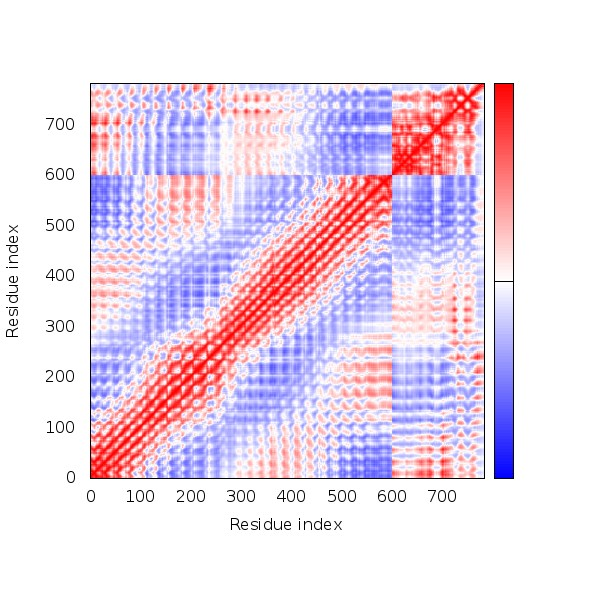

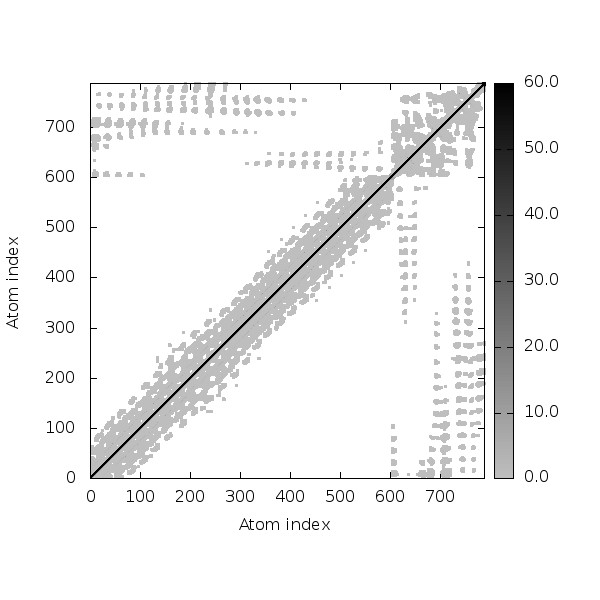

**Figure S13: SPVC_214(TLR4) simulation results [A) Deformability plot, B) B-factor plot, C) Eigenvalue plot , D) Covariance map [correlated (red), uncorrelated (white) or anti-correlated (blue) motions] , E) Elastic network (darker gray regions indicate more stiffer regions) of the complex]**

**A	B**

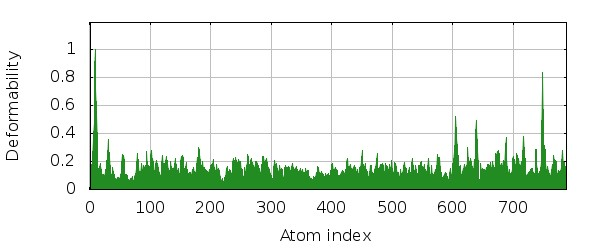

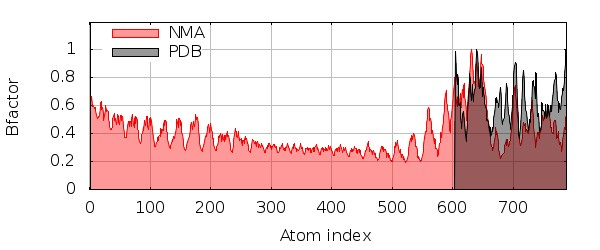

**C**

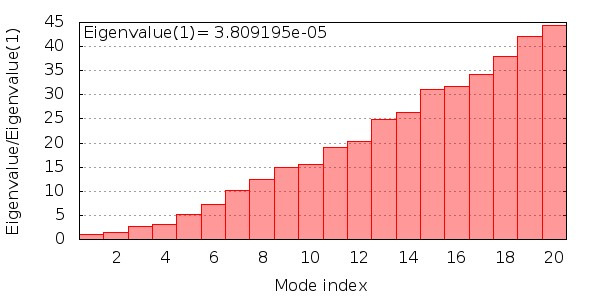

**D	E**

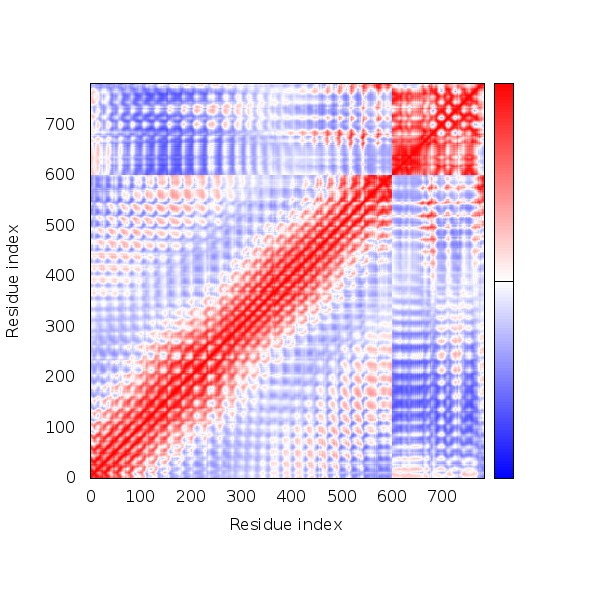

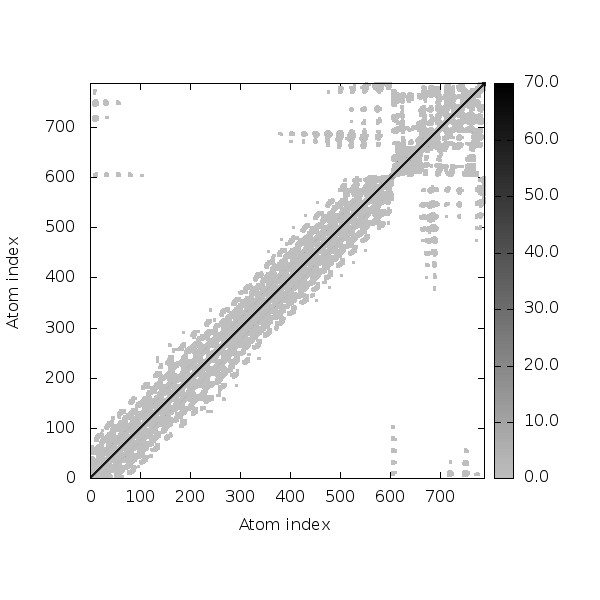

**Figure S14: SPVC_357(TLR4) simulation results [A) Deformability plot, B) B-factor plot, C) Eigenvalue plot , D) Covariance map [correlated (red), uncorrelated (white) or anti-correlated (blue) motions] , E) Elastic network (darker gray regions indicate more stiffer regions) of the complex]**

**A	B**

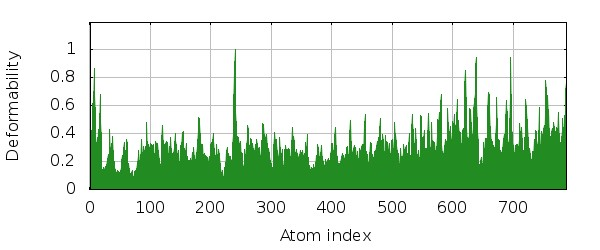

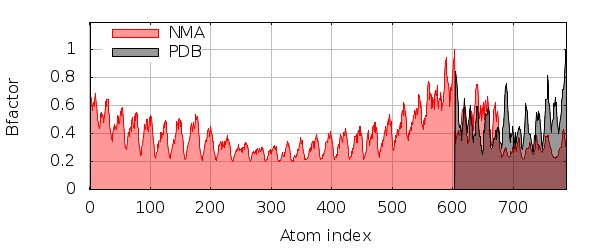

**C**

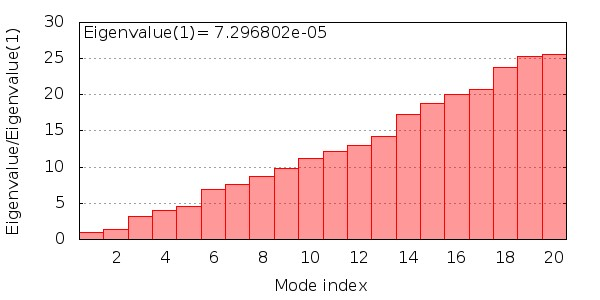

**D	E**

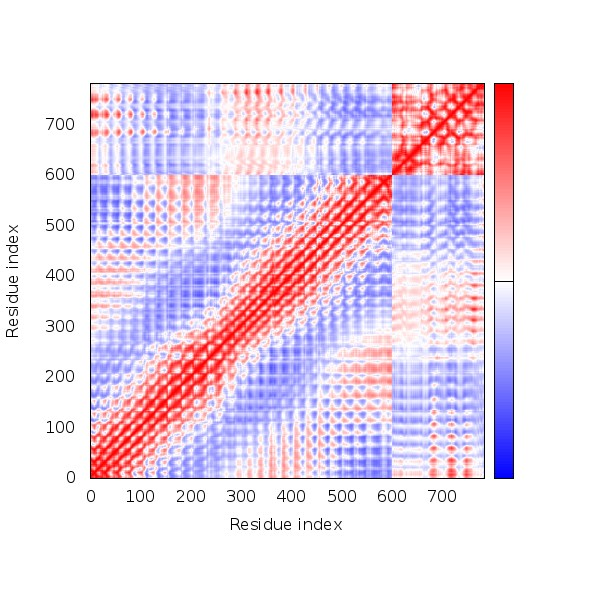

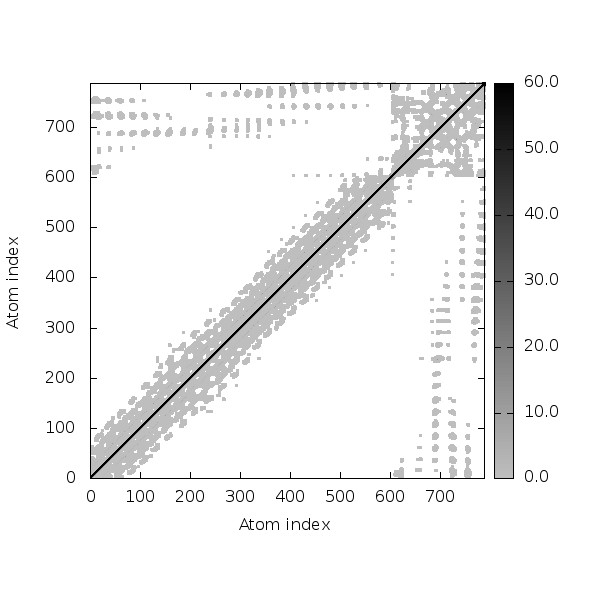

**Figure S15: SPVC_383(TLR4) simulation results [A) Deformability plot, B) B-factor plot, C) Eigenvalue plot , D) Covariance map [correlated (red), uncorrelated (white) or anti-correlated (blue) motions] , E) Elastic network (darker gray regions indicate more stiffer regions) of the complex]**

**A	B**

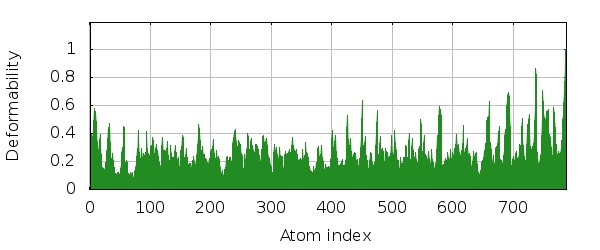

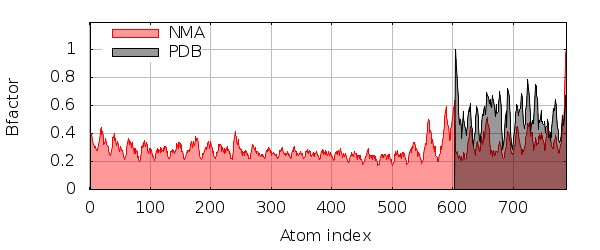

**C**

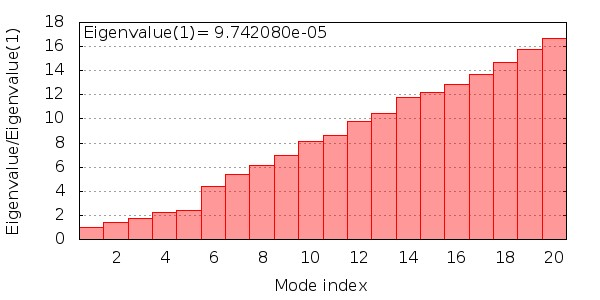

**D	E**

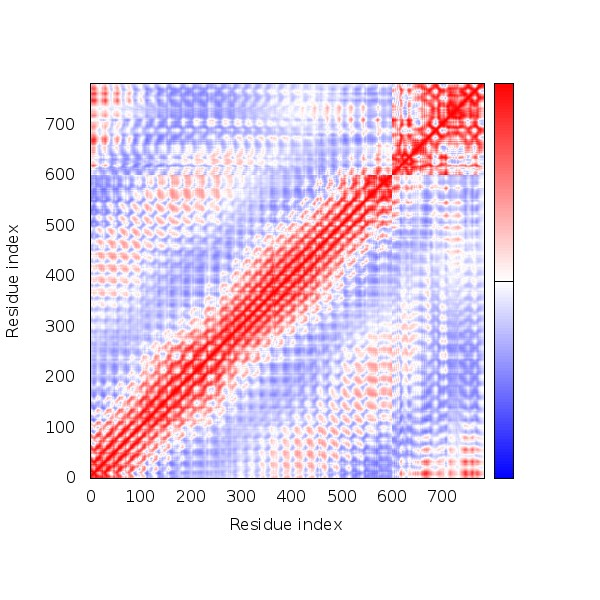

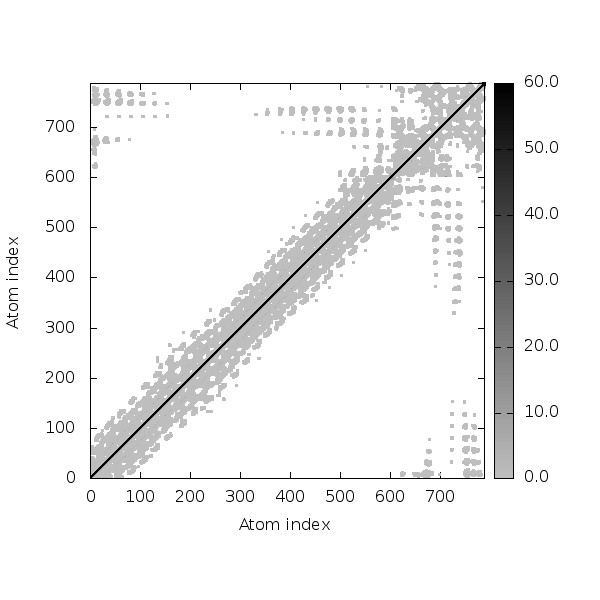

**Figure S16: SPVC_387(TLR4) simulation results [A) Deformability plot, B) B-factor plot, C) Eigenvalue plot , D) Covariance map [correlated (red), uncorrelated (white) or anti-correlated (blue) motions] , E) Elastic network (darker gray regions indicate more stiffer regions) of the complex]**

**A	B**

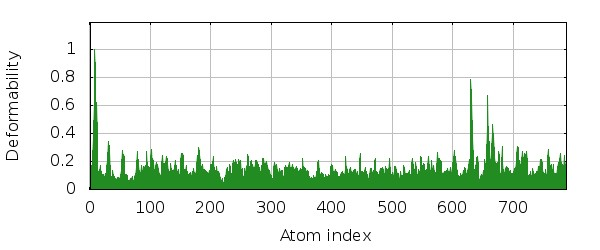

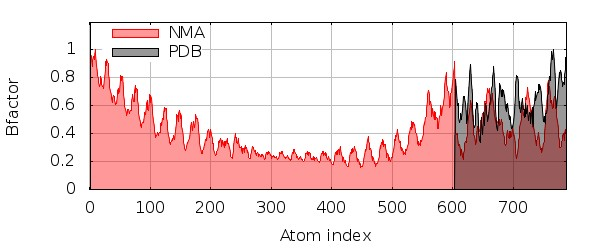

**C**

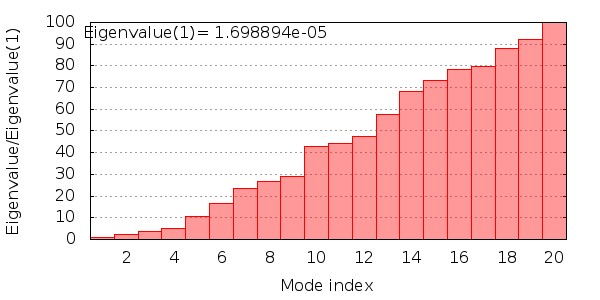

**D	E**

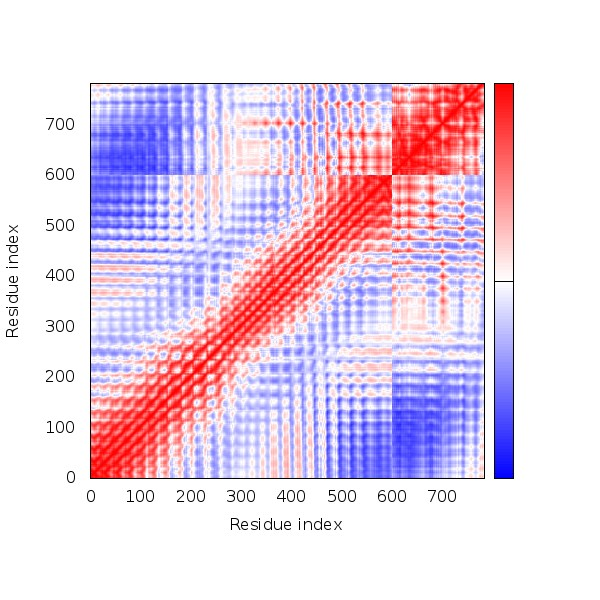

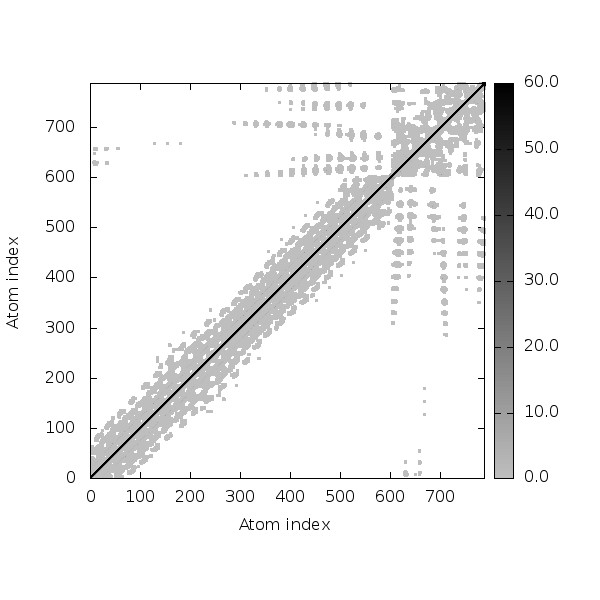

**Figure S17: SPVC_446(TLR4) simulation results [A) Deformability plot, B) B-factor plot, C) Eigenvalue plot , D) Covariance map [correlated (red), uncorrelated (white) or anti-correlated (blue) motions] , E) Elastic network (darker gray regions indicate more stiffer regions) of the complex]**

**A	B**

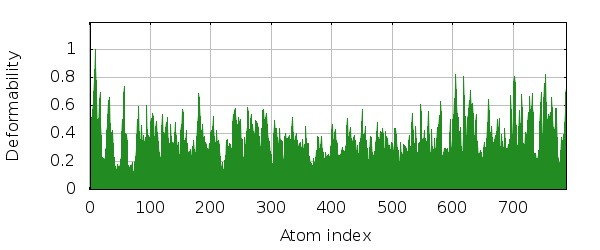

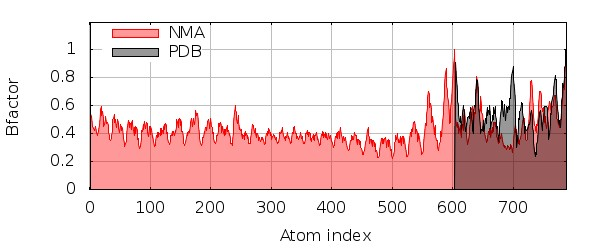

**C**

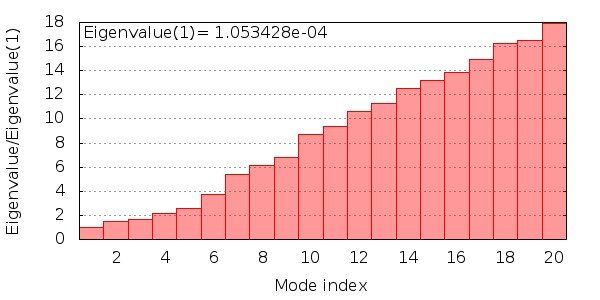

**D	E**

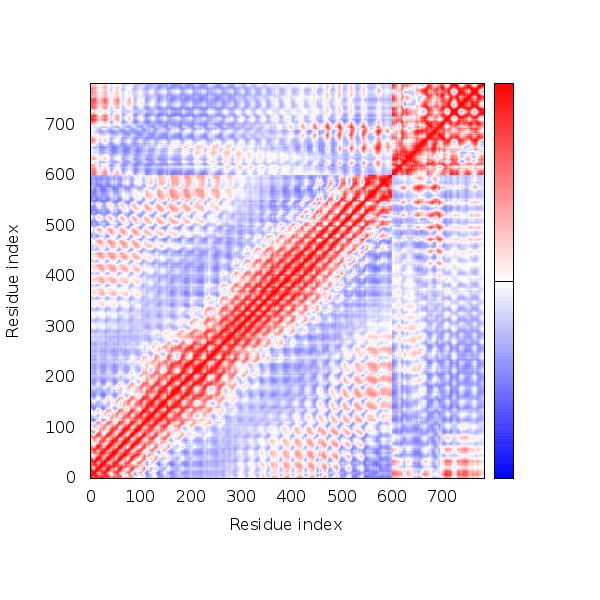

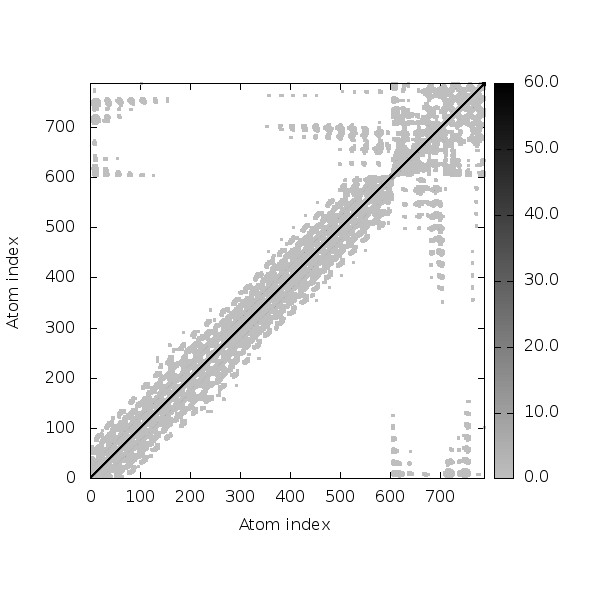

**Figure S18: SPVC_537(TLR4) simulation results [A) Deformability plot, B) B-factor plot, C) Eigenvalue plot , D) Covariance map [correlated (red), uncorrelated (white) or anti-correlated (blue) motions] , E) Elastic network (darker gray regions indicate more stiffer regions) of the complex]**

**A	B**

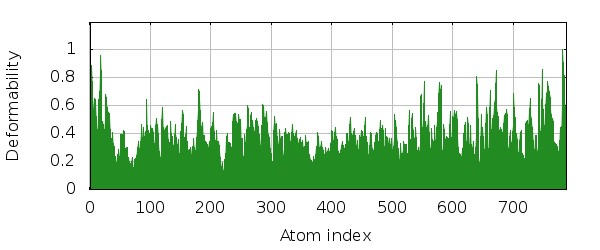

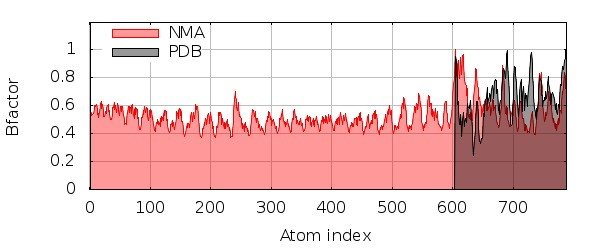

**C**

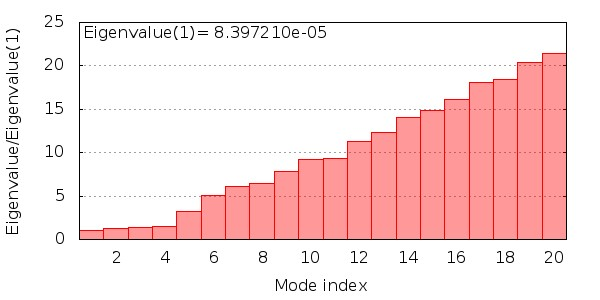

**D	E**

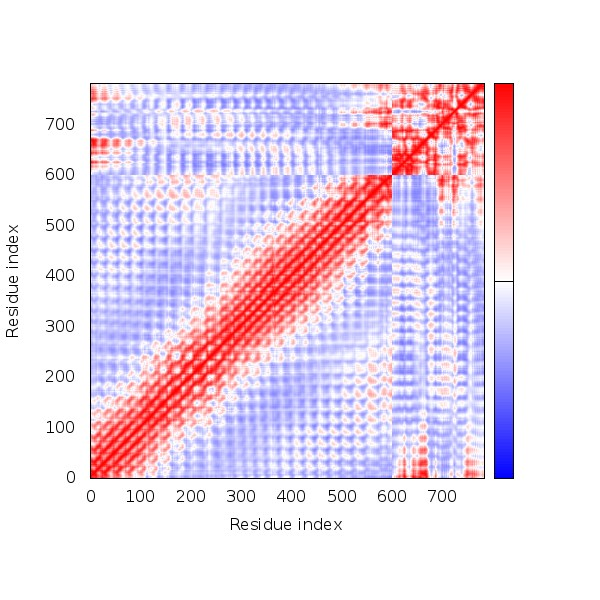

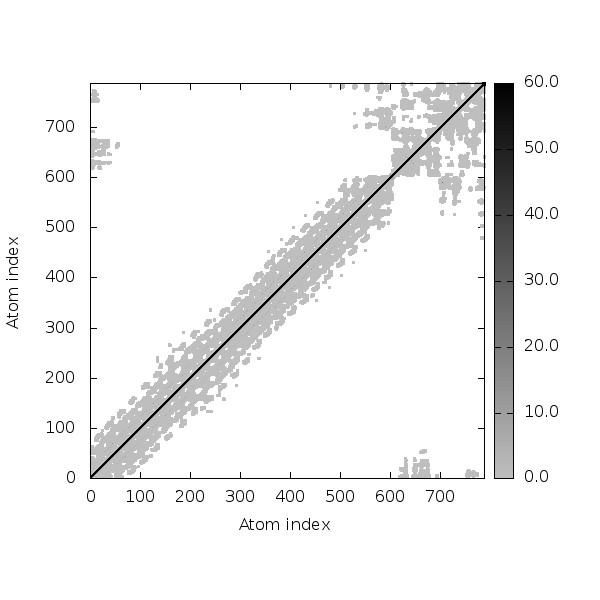

**Figure S19: SPVC_565(TLR4) simulation results [A) Deformability plot, B) B-factor plot, C) Eigenvalue plot , D) Covariance map [correlated (red), uncorrelated (white) or anti-correlated (blue) motions] , E) Elastic network (darker gray regions indicate more stiffer regions) of the complex]**

**A	B**

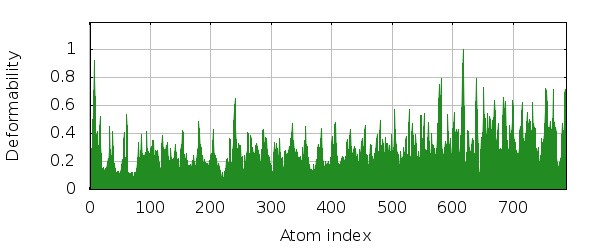

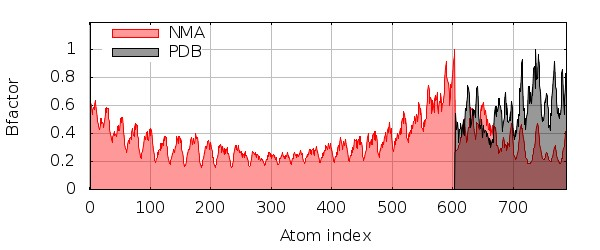

**C**

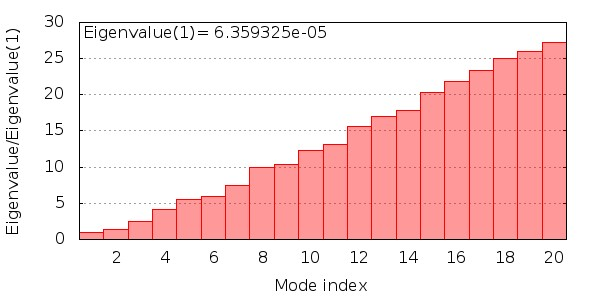

**D	E**

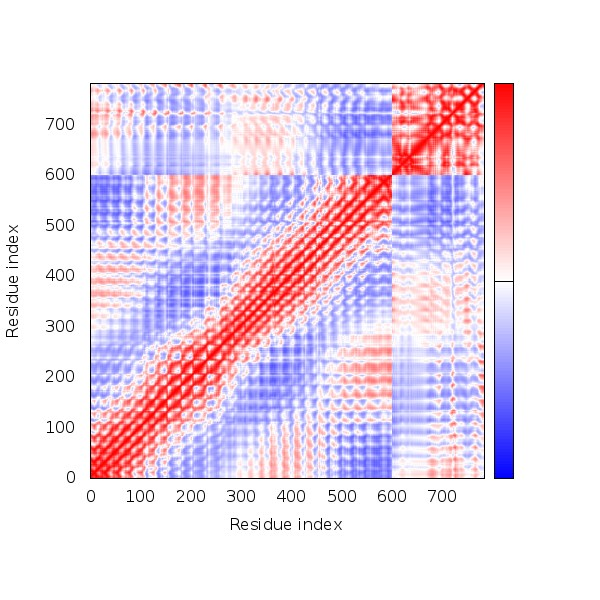

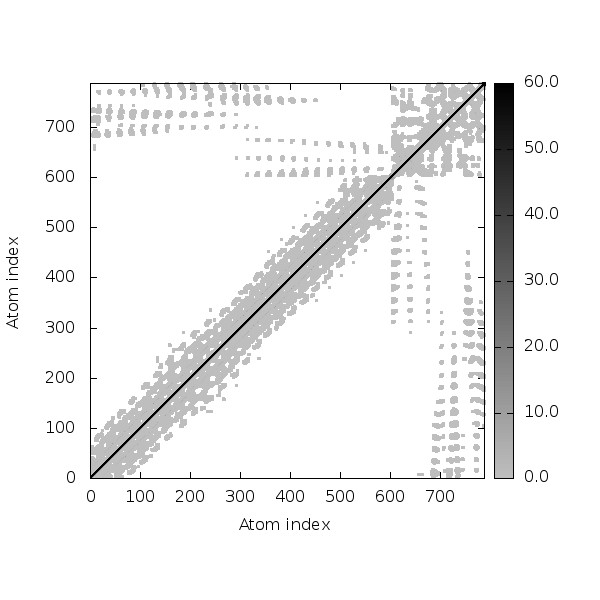

**Figure S20: REF_SEQ(TLR4) simulation results [A) Deformability plot, B) B-factor plot, C) Eigenvalue plot , D) Covariance map [correlated (red), uncorrelated (white) or anti-correlated (blue) motions] , E) Elastic network (darker gray regions indicate more stiffer regions) of the complex]**
