## Supplementary Material 3 for "Epitope order Matters in multi-epitope-based peptide (MEBP) vaccine design: An *in silico* study"

**C-Immsim results**

**A	B**

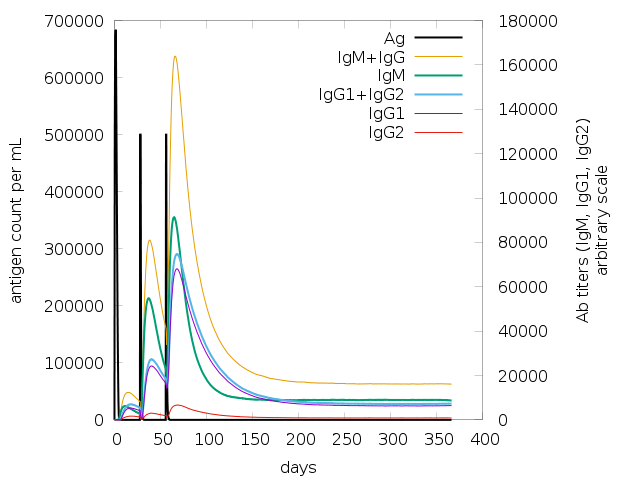

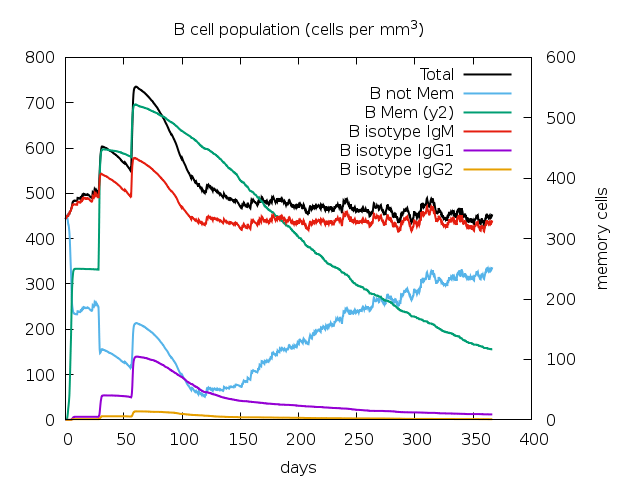

**C	D**

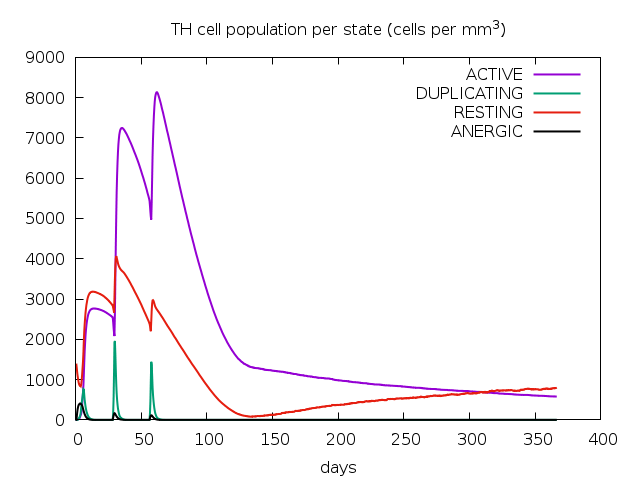

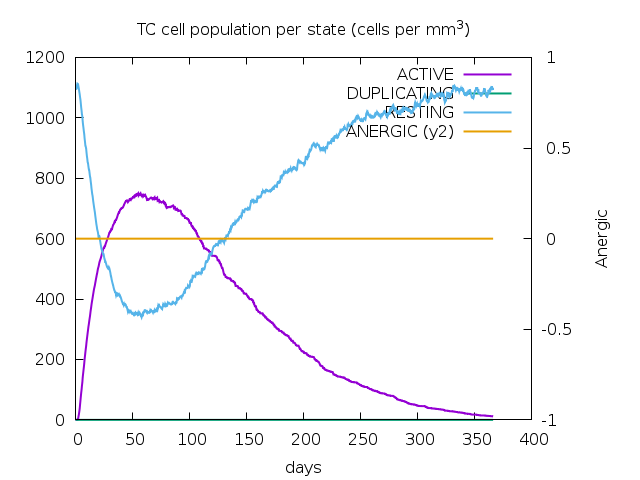

**Figure S1: SPVC_214 with adjuvants+HIS-tag:A)Antigen and immunoglobulin counts B)The changes observed in B-cell populations after given three injections, C) The development of T-helper, and D) T-cytotoxic cell populations per state after the injections**

**A	B**

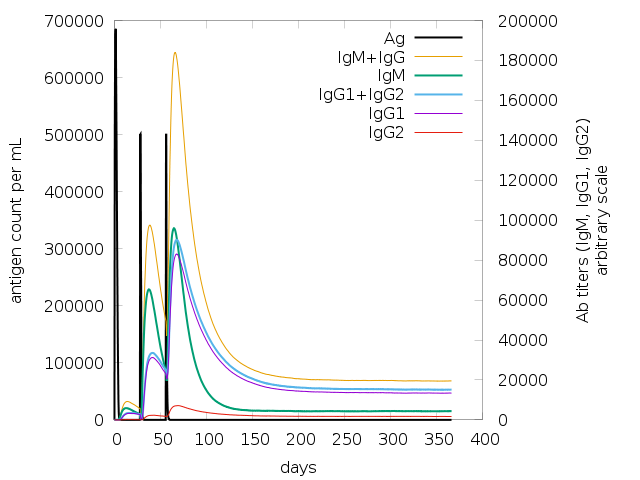

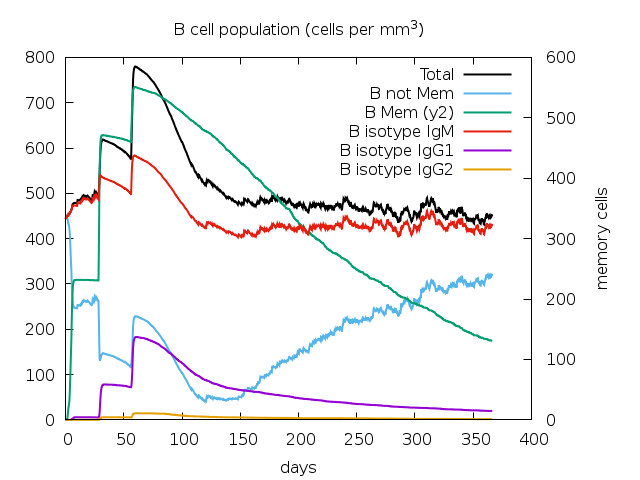

**C	D**

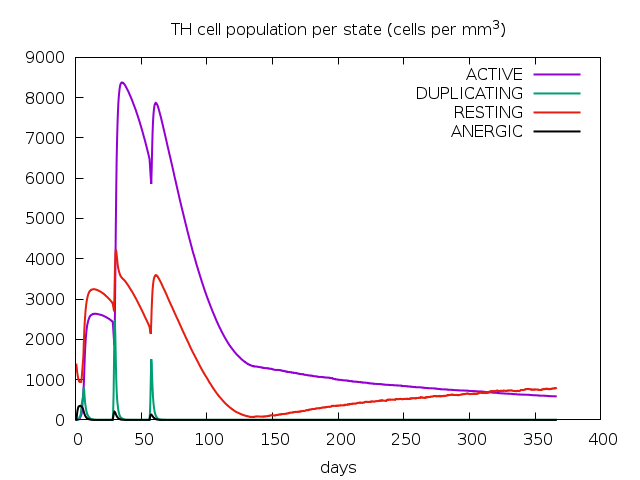

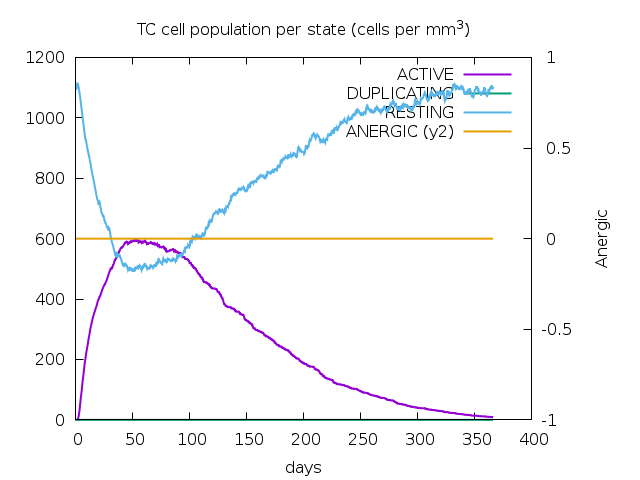

**Figure S2: SPVC_32 with adjuvants+HIS-tag:A)Antigen and immunoglobulin counts B)The changes observed in B-cell populations after given three injections, C) The development of T-helper, and D) T-cytotoxic cell populations per state after the injections**

**A	B**

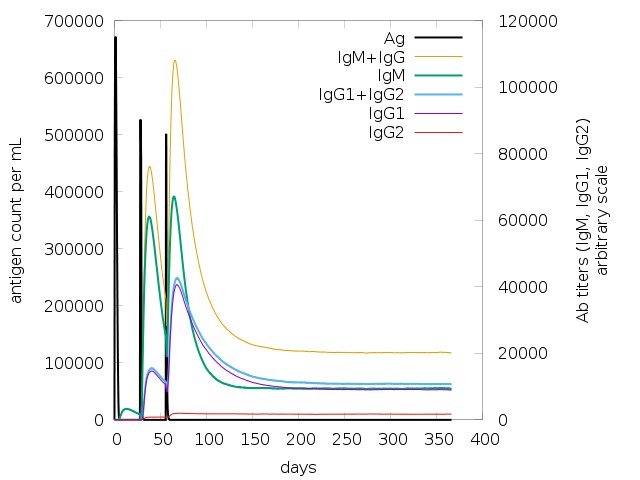

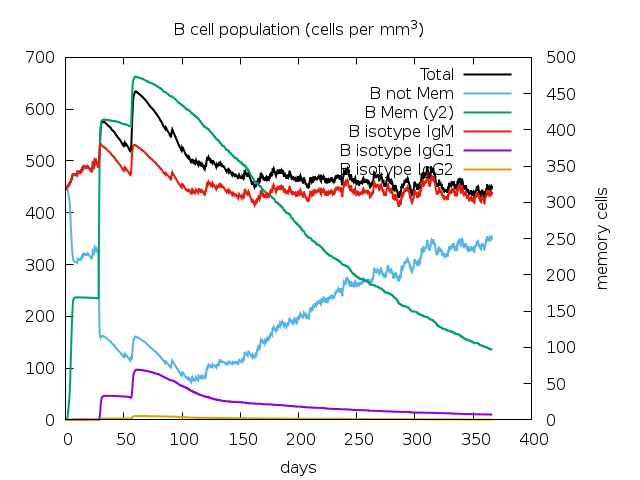

**C	D**

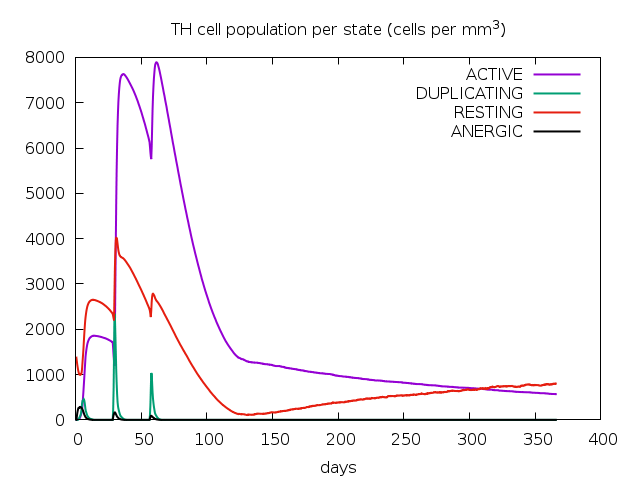

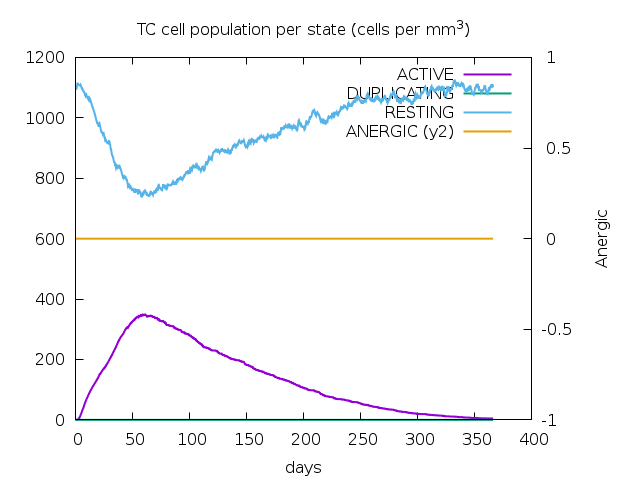

**Figure S3: SPVC_206 with adjuvants+HIS-tag:A)Antigen and immunoglobulin counts B)The changes observed in B-cell populations after given three injections, C) The development of T-helper, and D) T-cytotoxic cell populations per state after the injections**

**A	B**

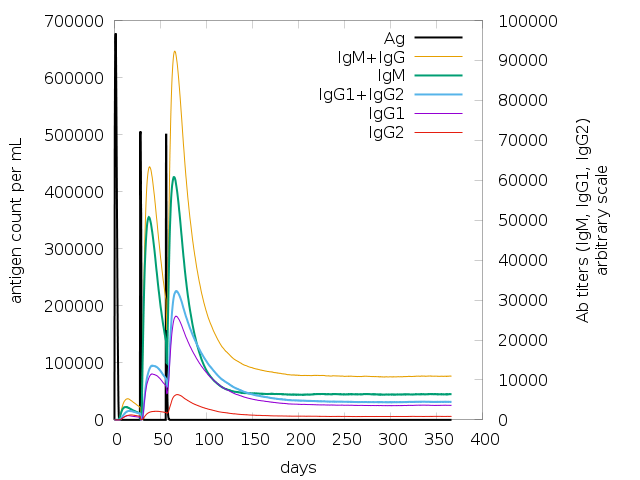

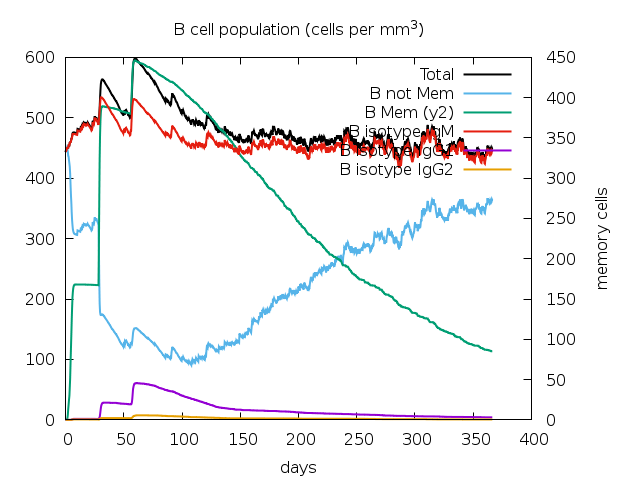

**C	D**

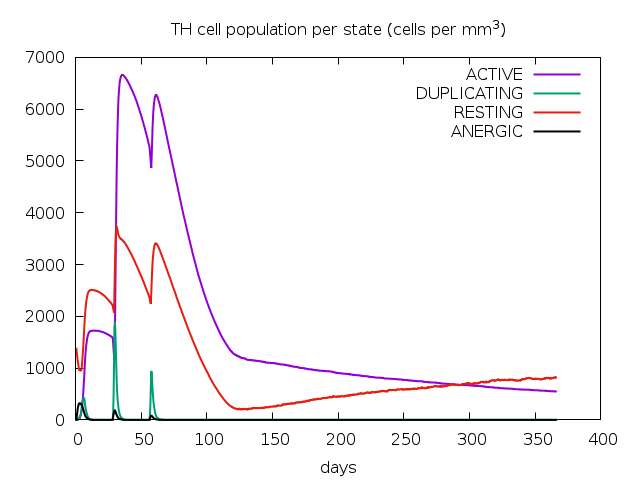

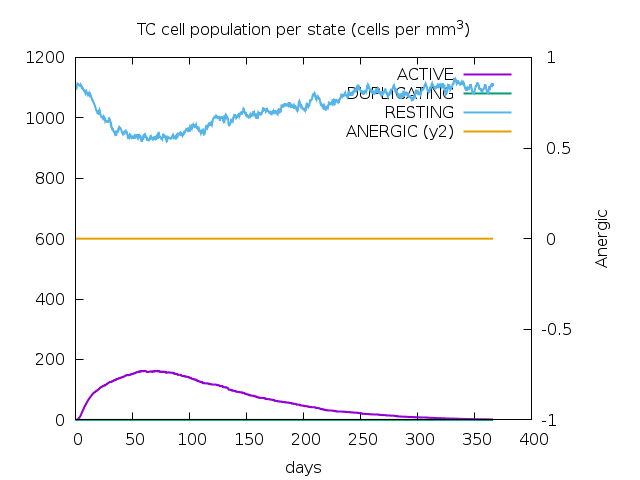

**Figure S4: SPVC_565 with adjuvants+HIS-tag:A)Antigen and immunoglobulin counts B)The changes observed in B-cell populations after given three injections, C) The development of T-helper, and D) T-cytotoxic cell populations per state after the injections**

**A	B**

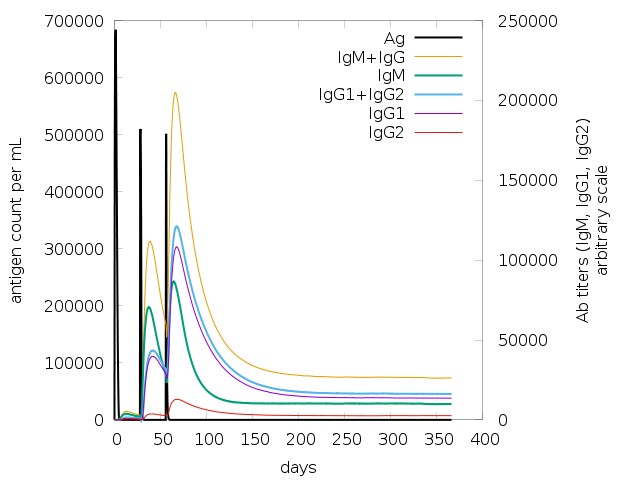

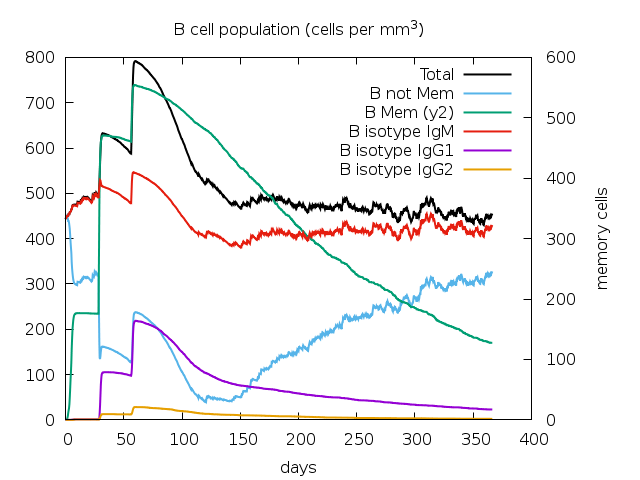

**D	E**

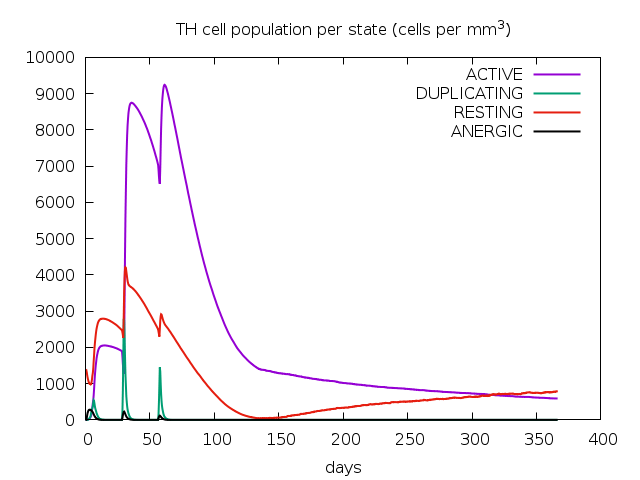

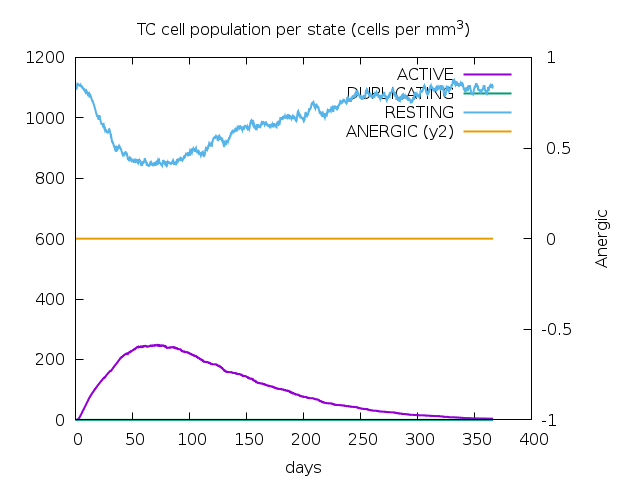

**Figure S5: SPVC_383 with adjuvants+HIS-tag:A)Antigen and immunoglobulin counts B)The changes observed in B-cell populations after given three injections, C) The development of T-helper, and D) T-cytotoxic cell populations per state after the injections**

**A	B**

**C**
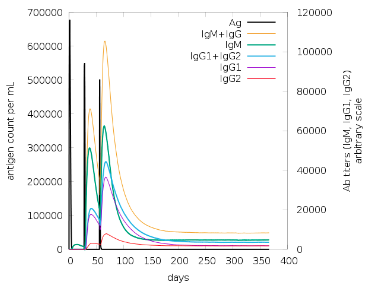
**D**
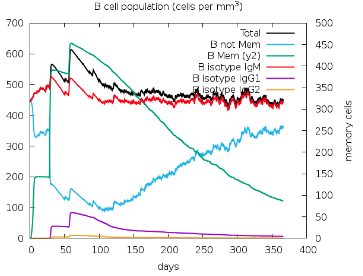

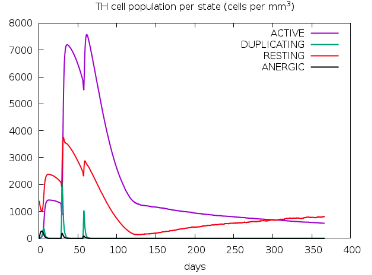

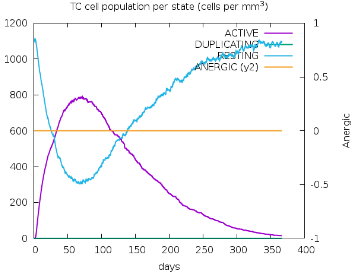

**Figure S6: SPVC_357 with adjuvants+HIS-tag:A)Antigen and immunoglobulin counts B)The changes observed in B-cell populations after given three injections, C) The development of T-helper, and D) T-cytotoxic cell populations per state after the injections**

**A	B**

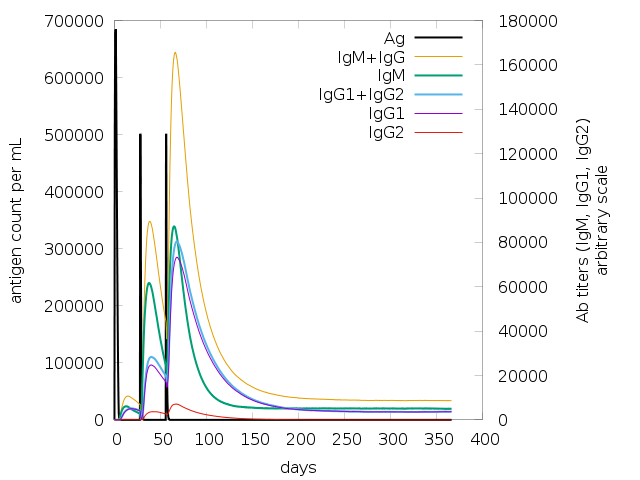

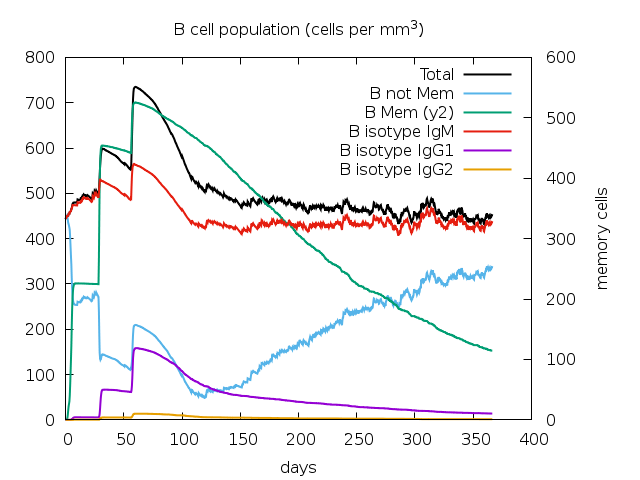

**C	D**

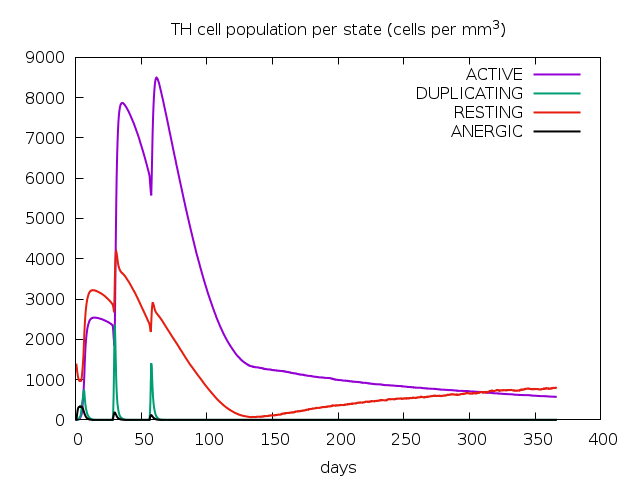

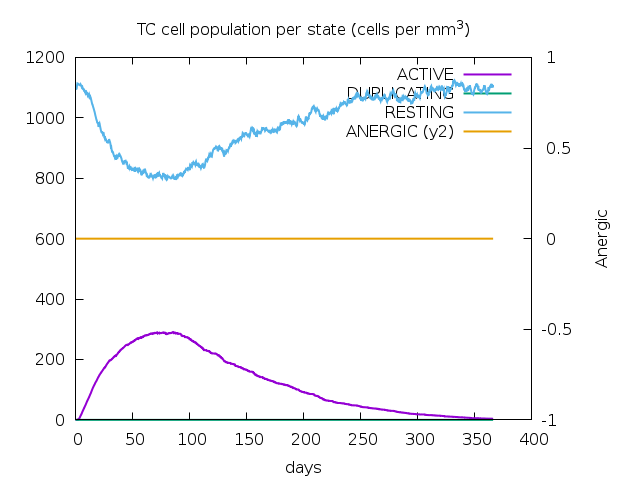

**Figure S7: SPVC_537 with adjuvants+HIS-tag:A)Antigen and immunoglobulin counts B)The changes observed in B-cell populations after given three injections, C) The development of T-helper, and D) T-cytotoxic cell populations per state after the injections**

**A	B**

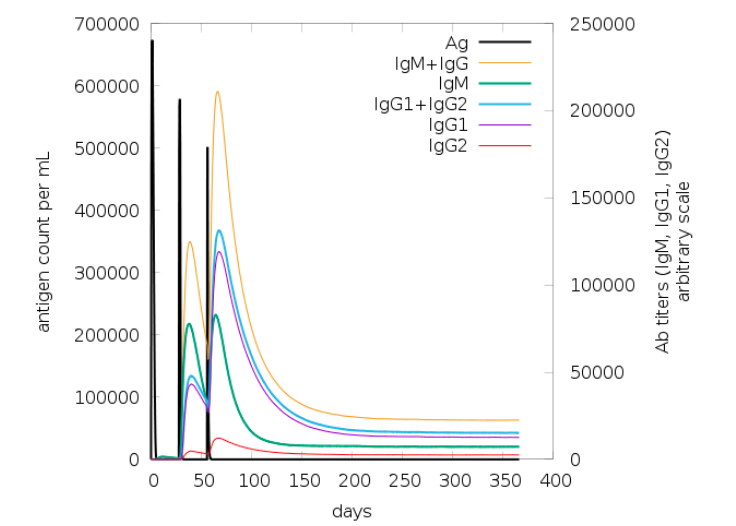

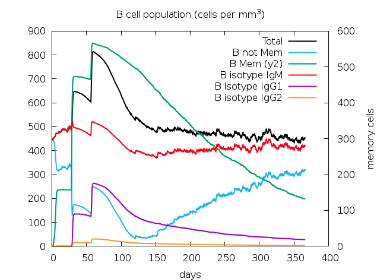

**C	D**

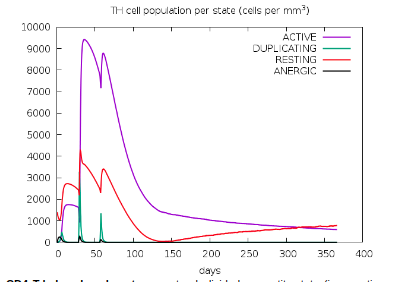

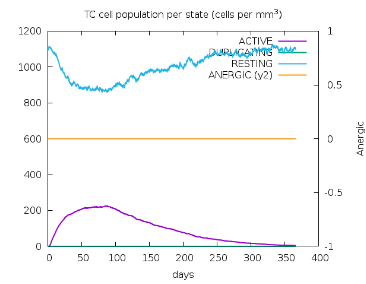

**Figure S8: REF_SEQ with adjuvants+HIS-tag: A)Antigen and immunoglobulin counts B)The changes observed in B-cell populations after given three injections, C) The development of T-helper, and D) T-cytotoxic cell populations per state after the injections**

**A	B**

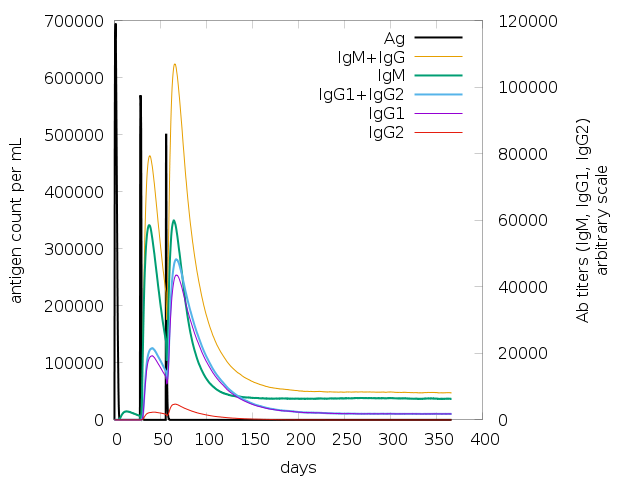

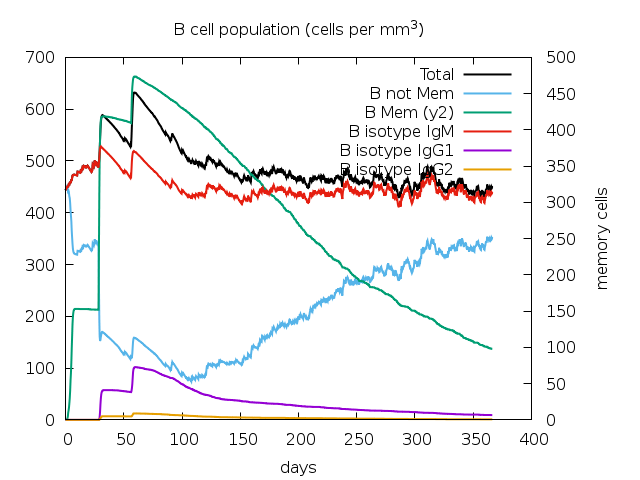

**C	D**

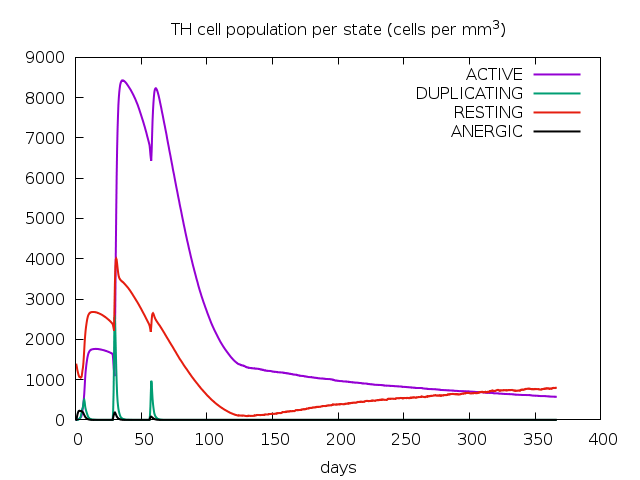

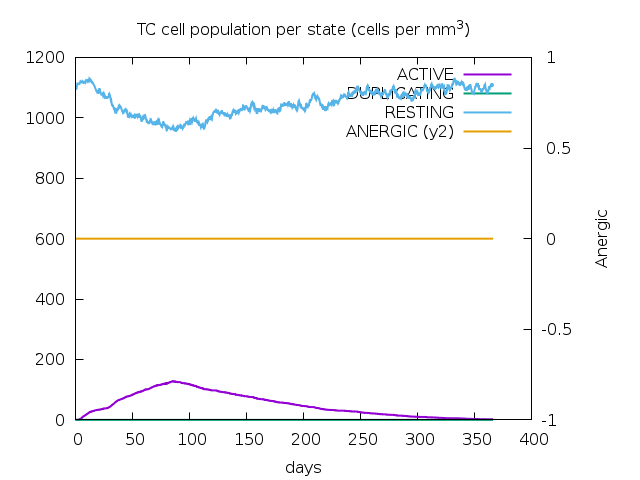

**Figure S9: SPVC_387 with adjuvants+HIS-tag:A)Antigen and immunoglobulin counts B)The changes observed in B-cell populations after given three injections, C) The development of T-helper, and D) T-cytotoxic cell populations per state after the injections**

**A	B**

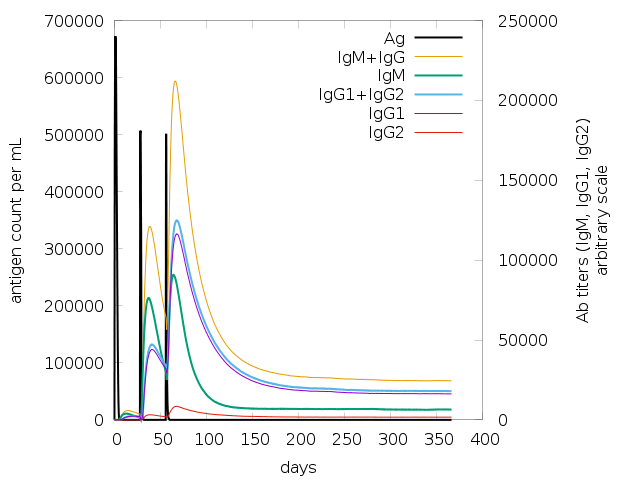

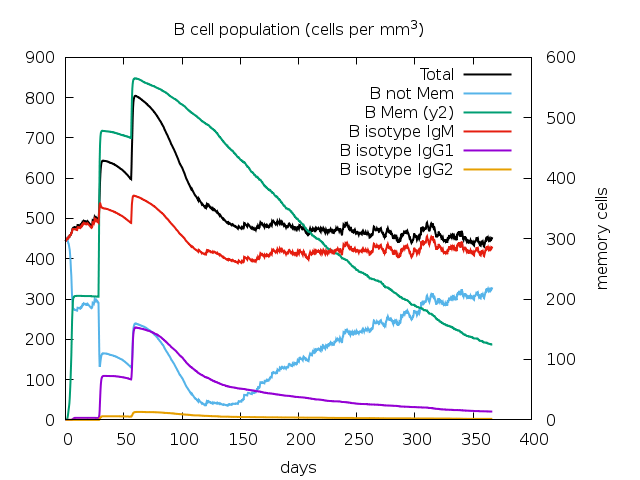

**C	D**

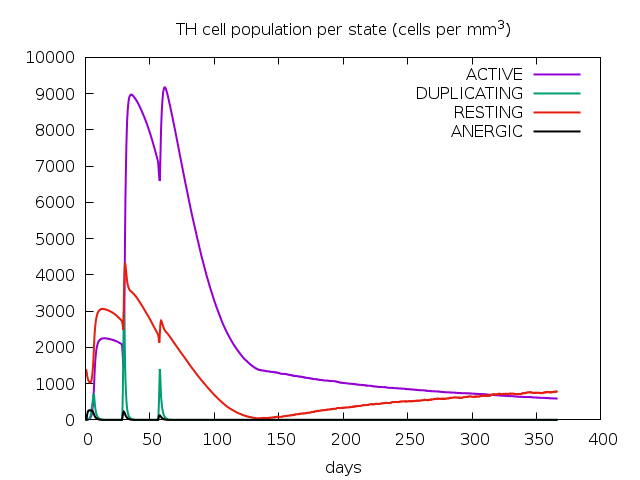

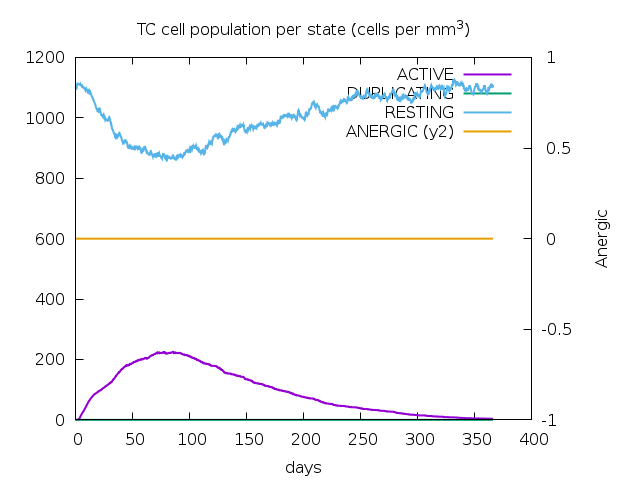

**Figure S10: SPVC_446 with adjuvants+HIS-tag:A)Antigen and immunoglobulin counts B)The changes observed in B-cell populations after given three injections, C) The development of T-helper, and D) T-cytotoxic cell populations per state after the injections**

**A	B**

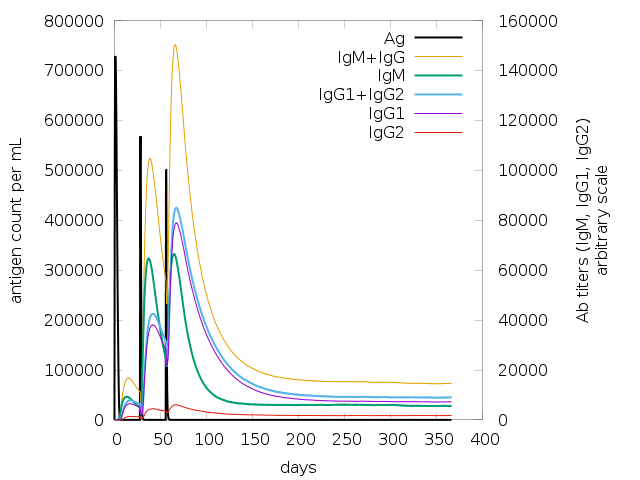

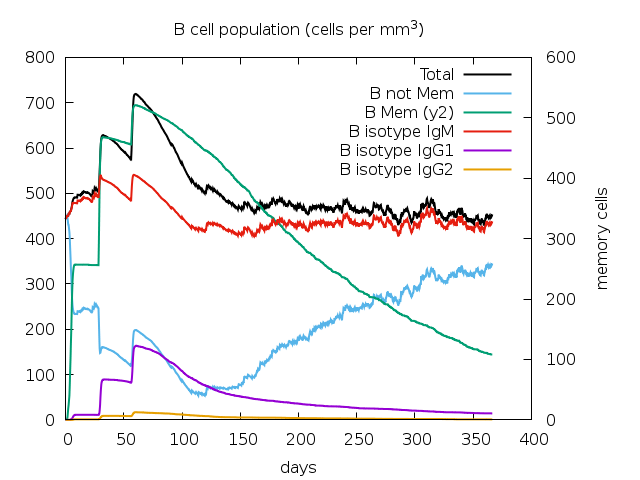

**C	D**

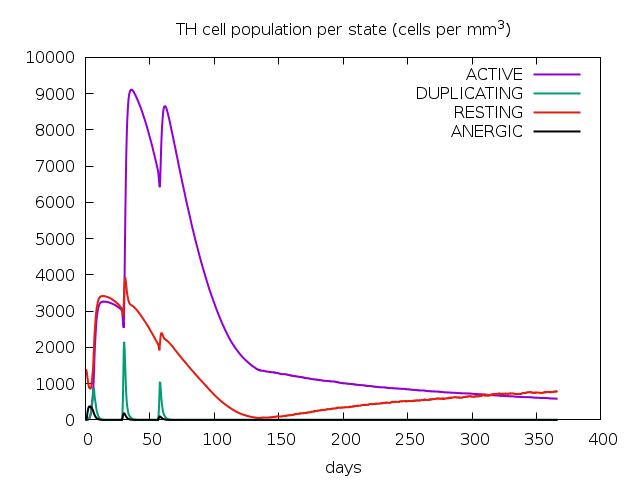

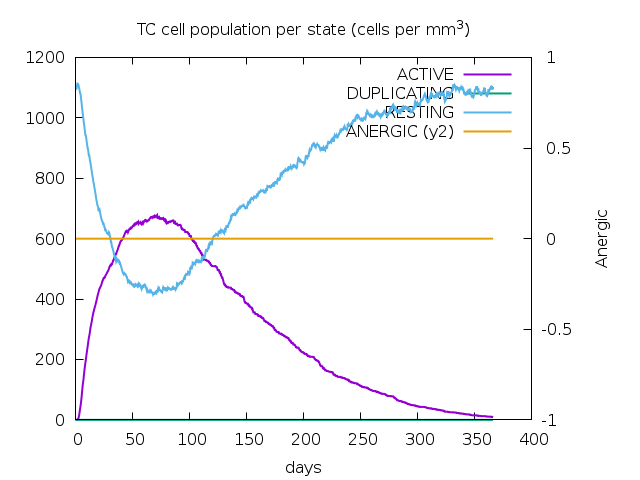

**Figure S11: SPVC_214 without adjuvants+HIS-tag:A)Antigen and immunoglobulin counts B)The changes observed in B-cell populations after given three injections, C) The development of T-helper, and D) T-cytotoxic cell populations per state after the injections**

**A	B**

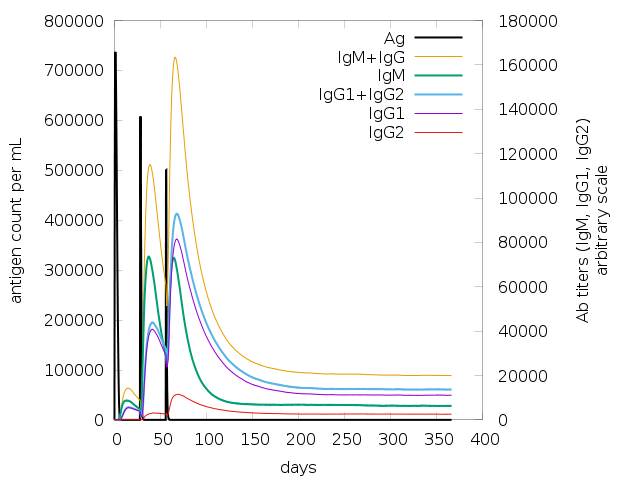

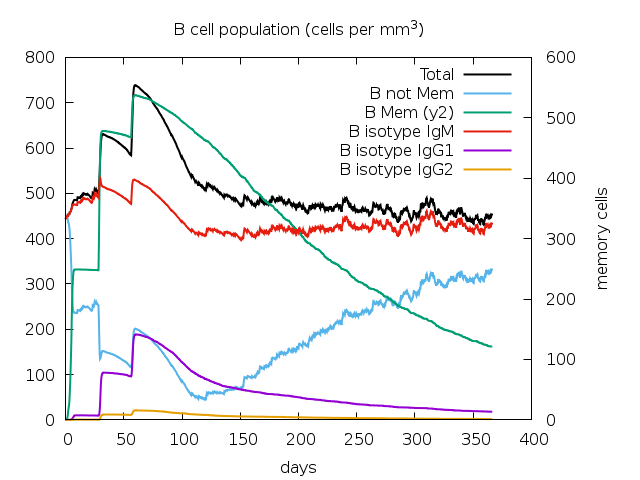

**C	D**

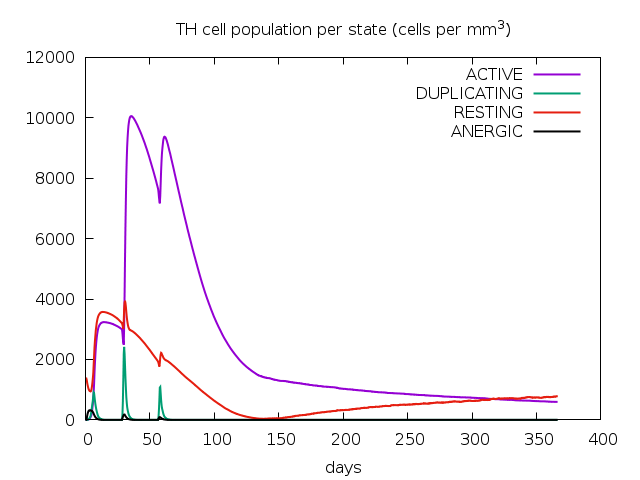

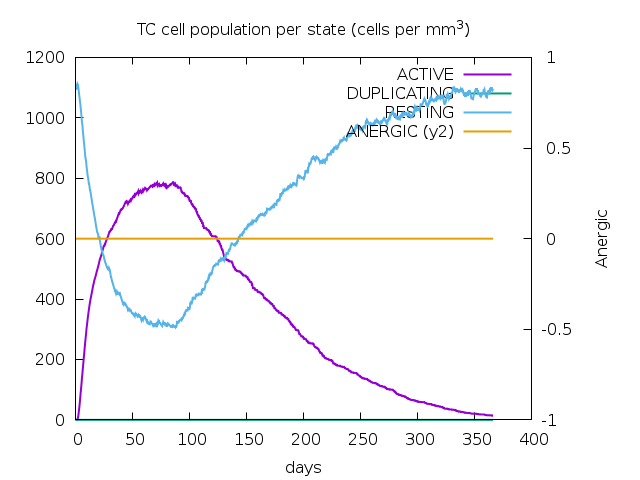

**Figure S12: SPVC_32 without adjuvants+HIS-tag:A)Antigen and immunoglobulin counts B)The changes observed in B-cell populations after given three injections, C) The development of T-helper, and D) T-cytotoxic cell populations per state after the injections**

**A	B**

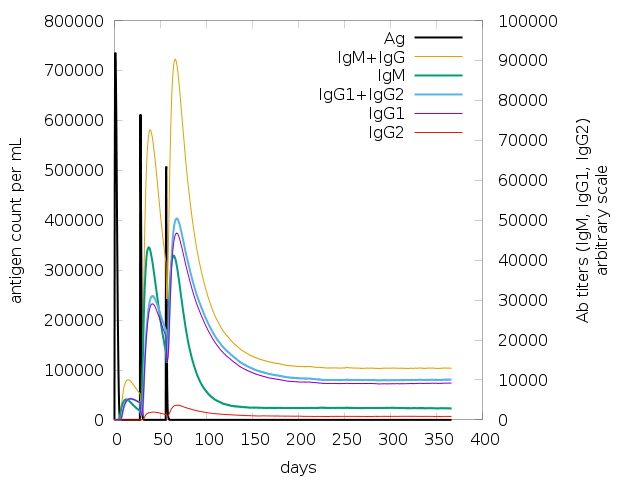

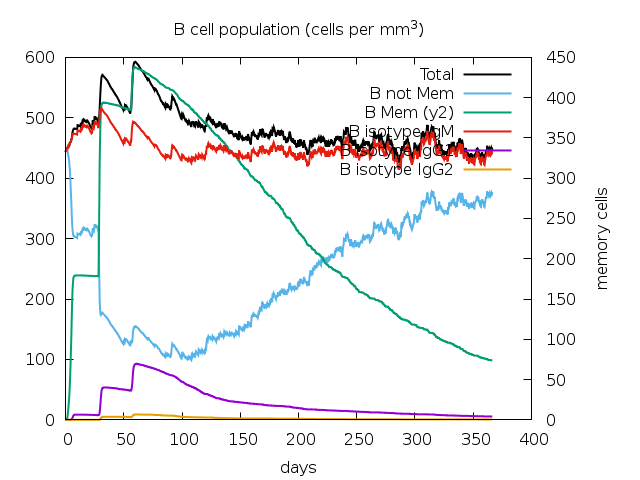

**C	D**

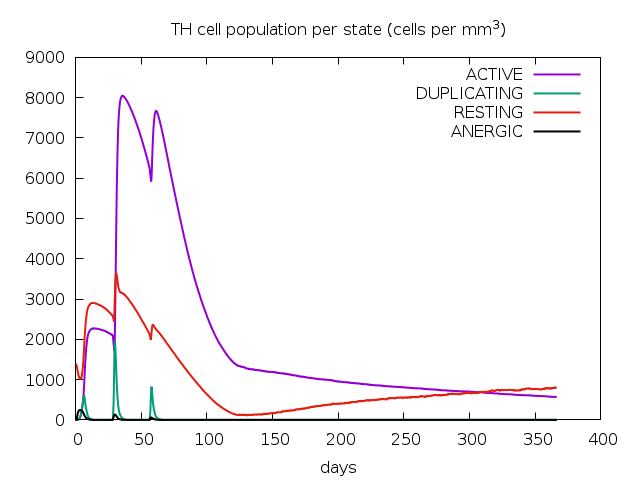

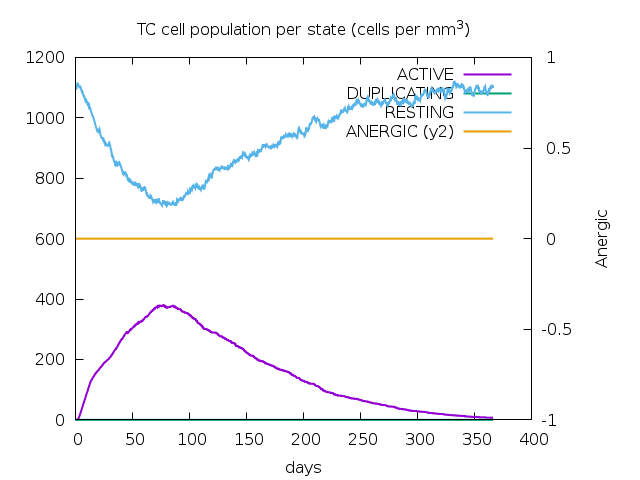

**Figure S13: SPVC_206 without adjuvants+HIS-tag:A) Antigen and immunoglobulin counts B)The changes observed in B-cell populations after given three injections, C) The development of T-helper, and D) T-cytotoxic cell populations per state after the injections**

**A	B**

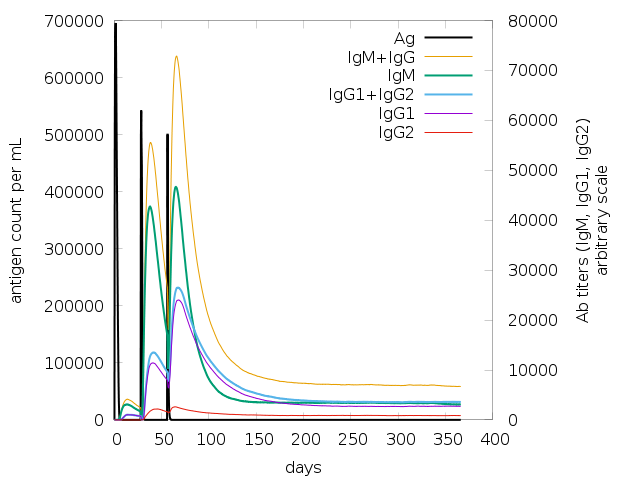

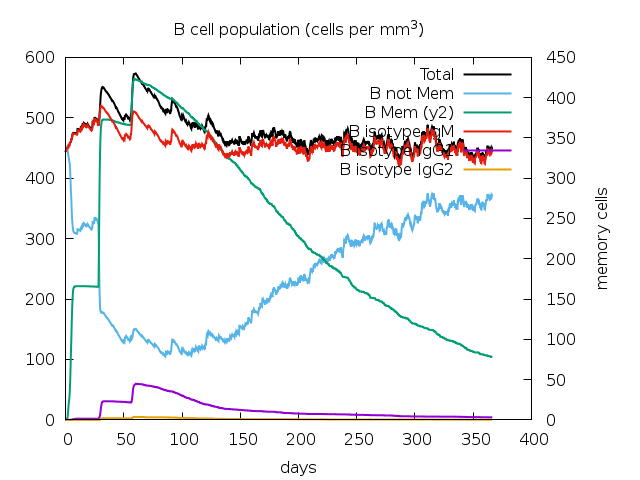

**C	D**

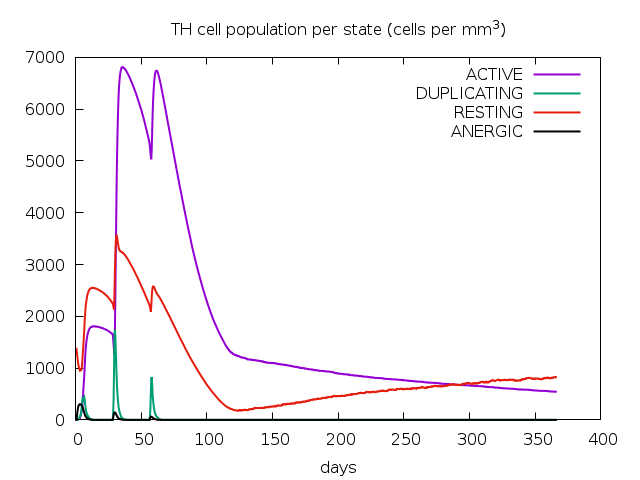

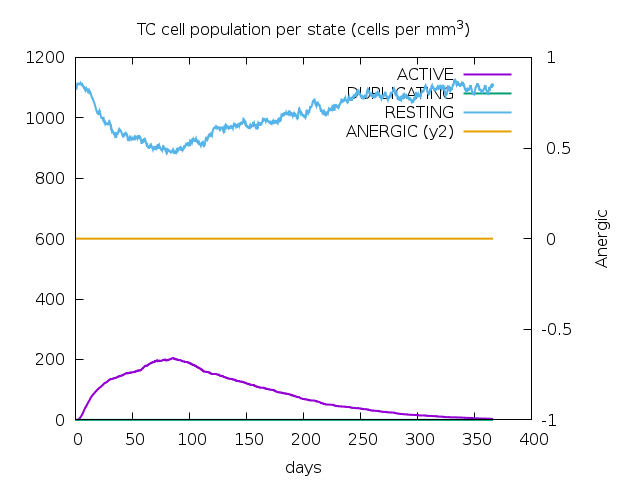

**Figure S14: SPVC_565 without adjuvants+HIS-tag:A)Antigen and immunoglobulin counts B)The changes observed in B-cell populations after given three injections, C) The development of T-helper, and D) T-cytotoxic cell populations per state after the injections**

**A	B**

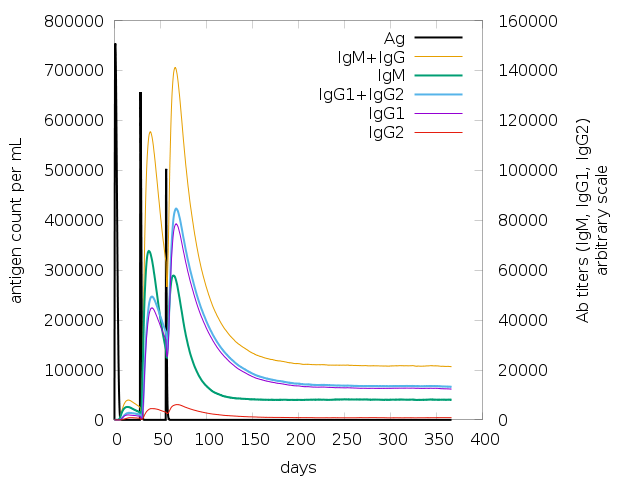

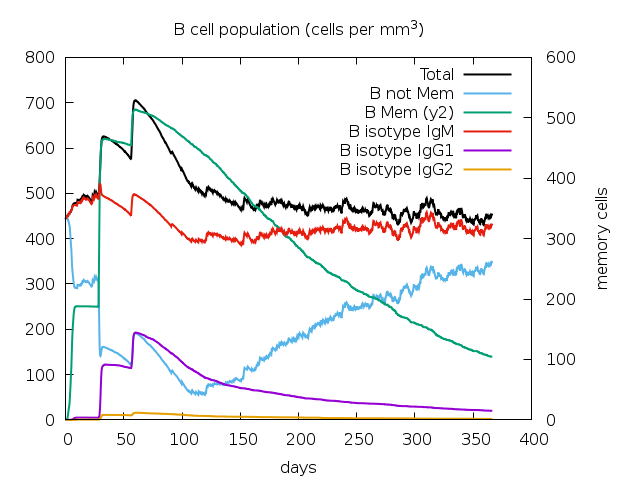

**C	D**

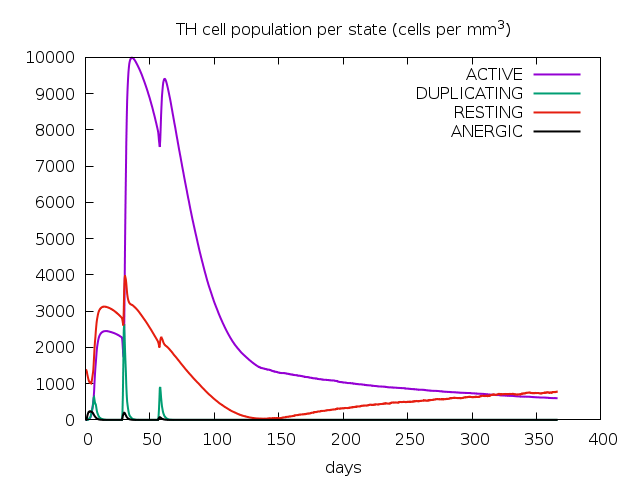

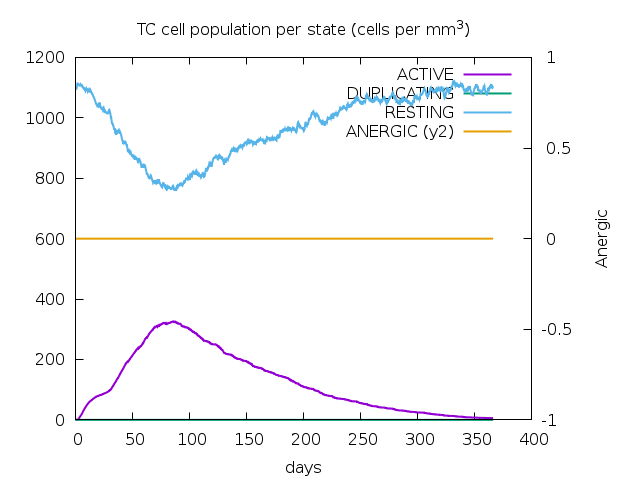

**Figure S15: SPVC_383 without adjuvants+HIS-tag:A)Antigen and immunoglobulin counts B)The changes observed in B-cell populations after given three injections, C) The development of T-helper, and D) T-cytotoxic cell populations per state after the injections**

**A	B**

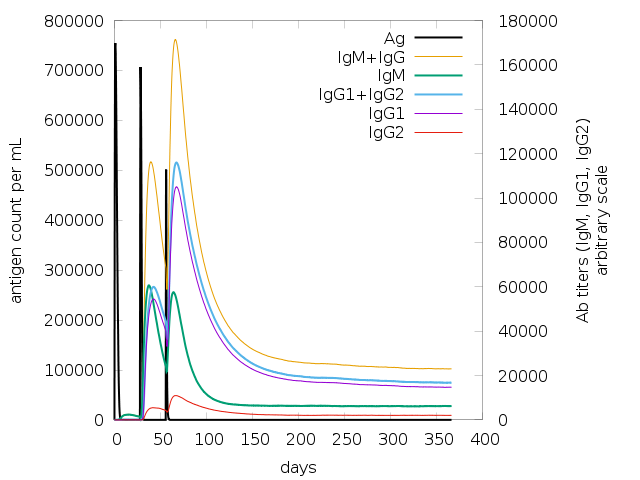

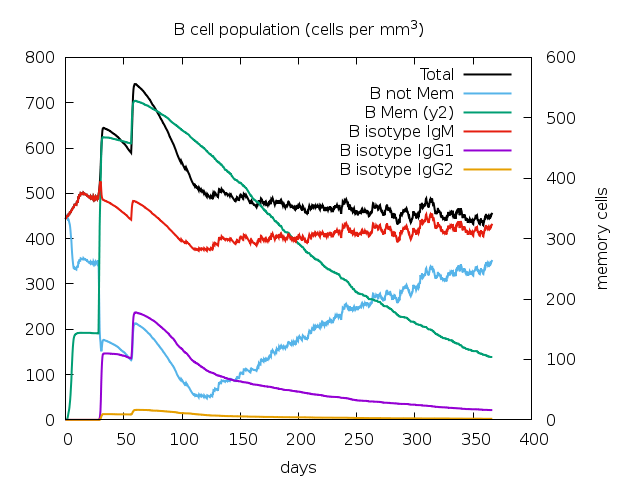

**C	D**

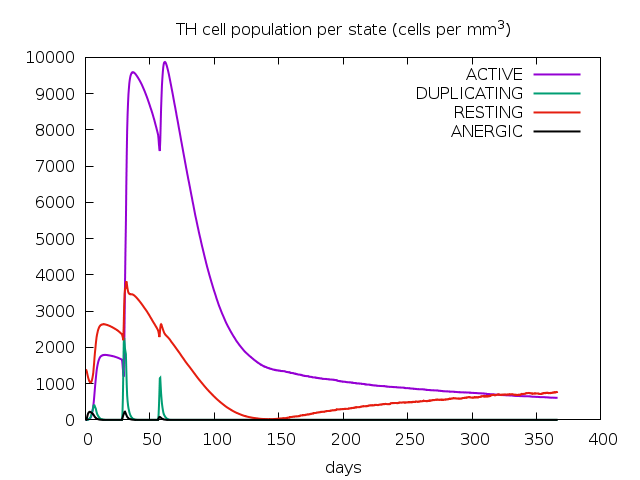

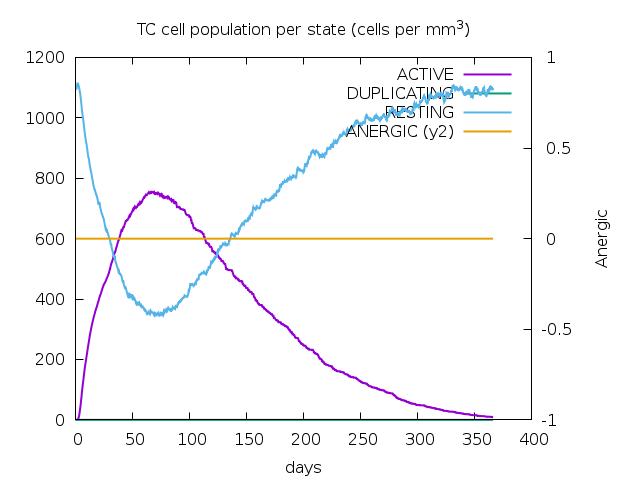

**Figure S16: SPVC_357 without adjuvants+HIS-tag:A)Antigen and immunoglobulin counts B)The changes observed in B-cell populations after given three injections, C) The development of T-helper, and D) T-cytotoxic cell populations per state after the injections**

**A	B**

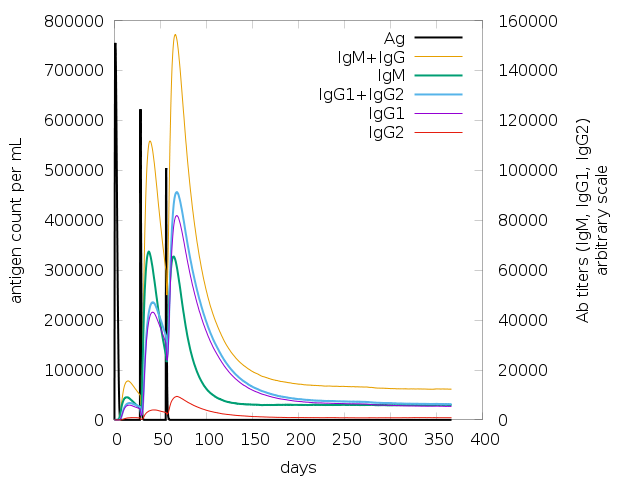

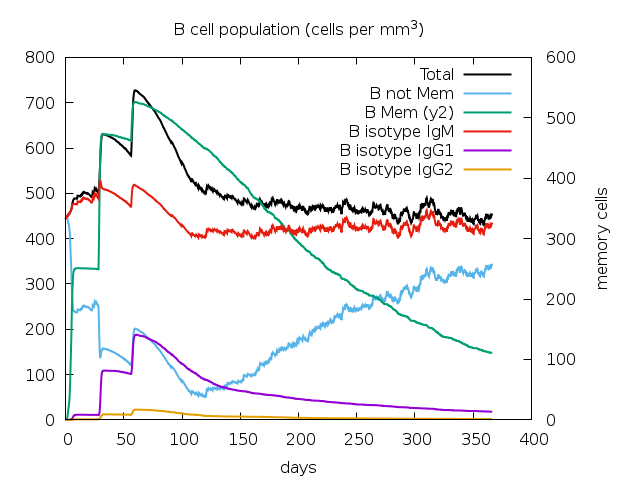

**C	D**

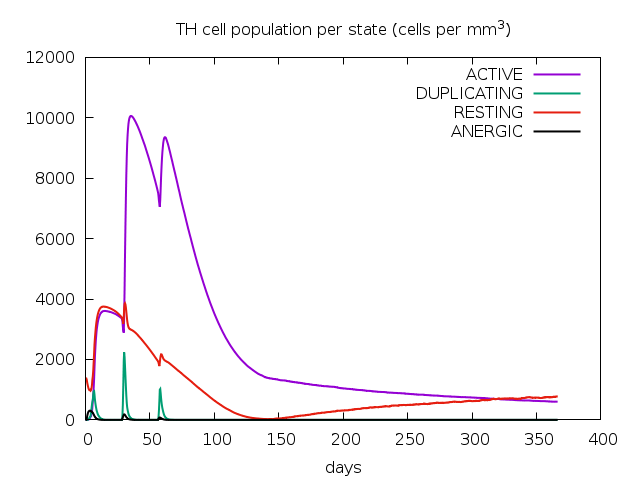

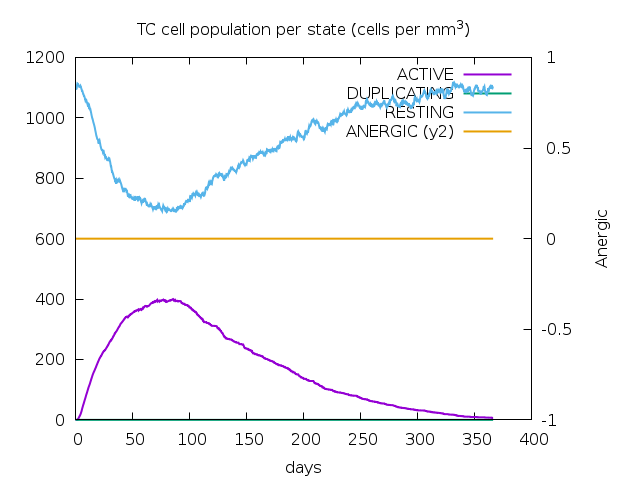

**Figure S17: SPVC_537 without adjuvants+HIS-tag:A)Antigen and immunoglobulin counts B)The changes observed in B-cell populations after given three injections, C) The development of T-helper, and D) T-cytotoxic cell populations per state after the injections**

**A	B**

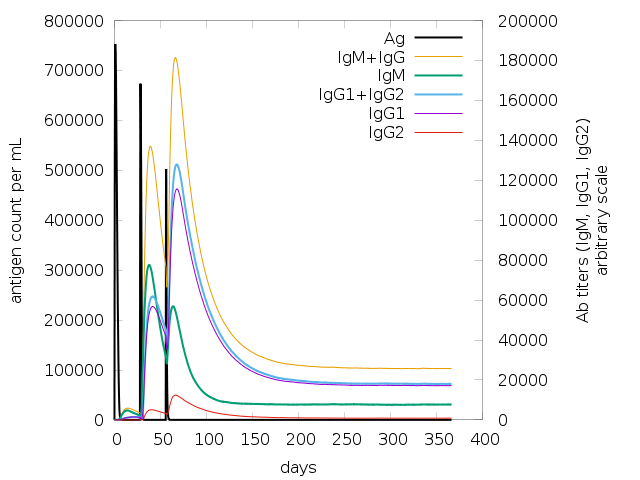

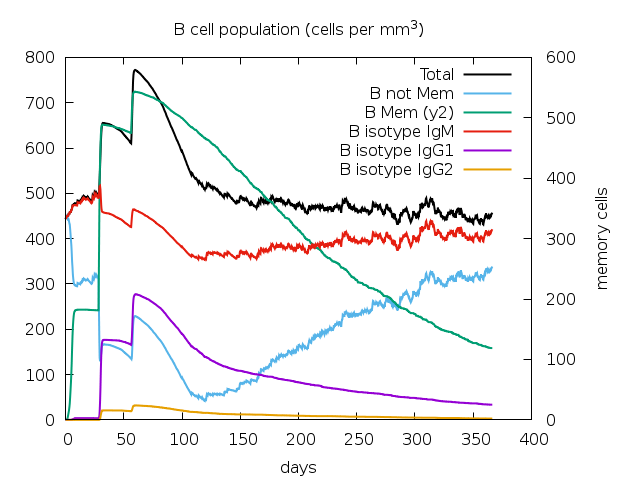

**C	D**

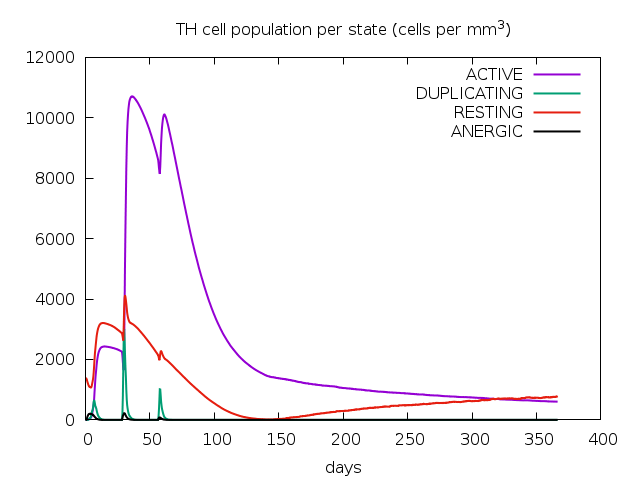

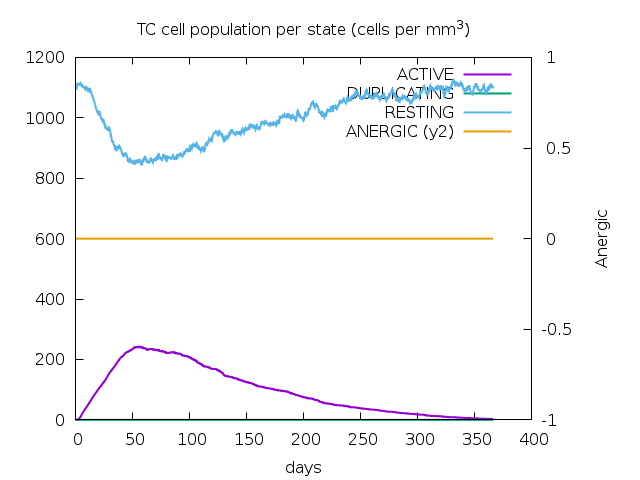

**Figure S18: REF_SEQ without adjuvants+HIS-tag:A)Antigen and immunoglobulin counts B)The changes observed in B-cell populations after given three injections, C) The development of T-helper, and D) T-cytotoxic cell populations per state after the injections**

**A	B**

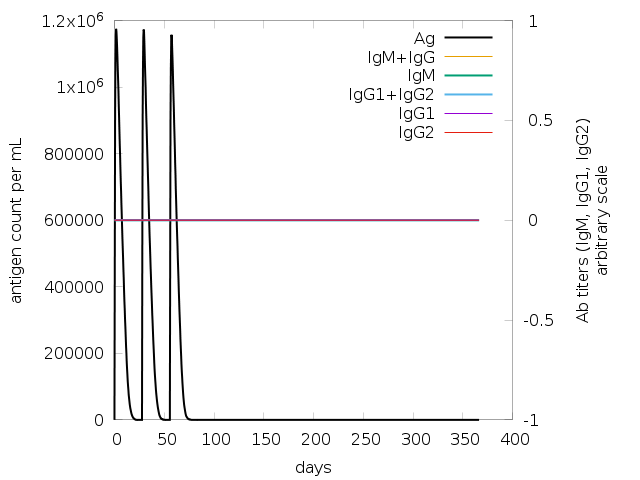

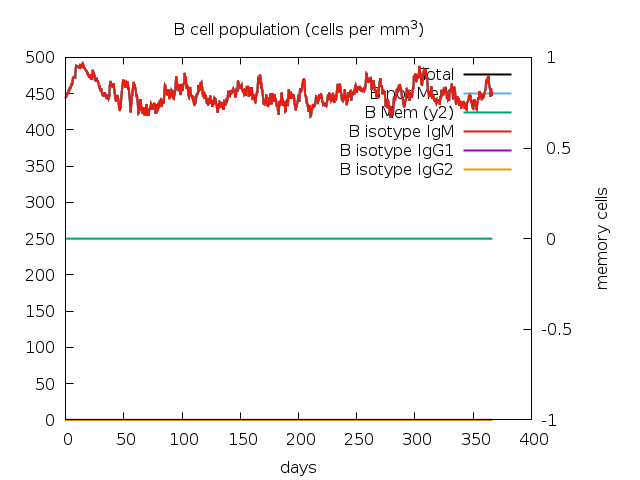

**C	D**

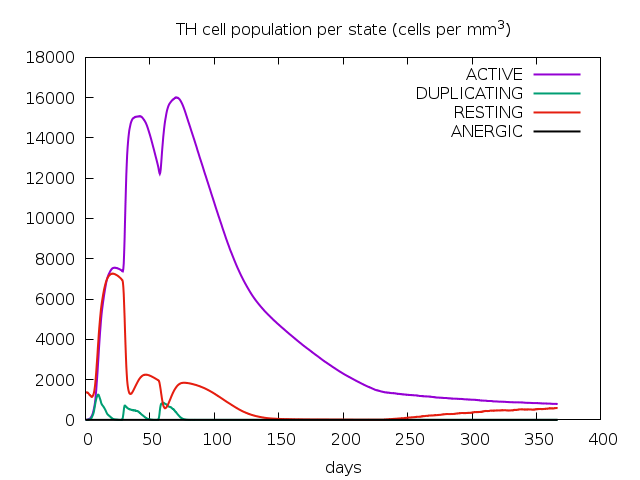

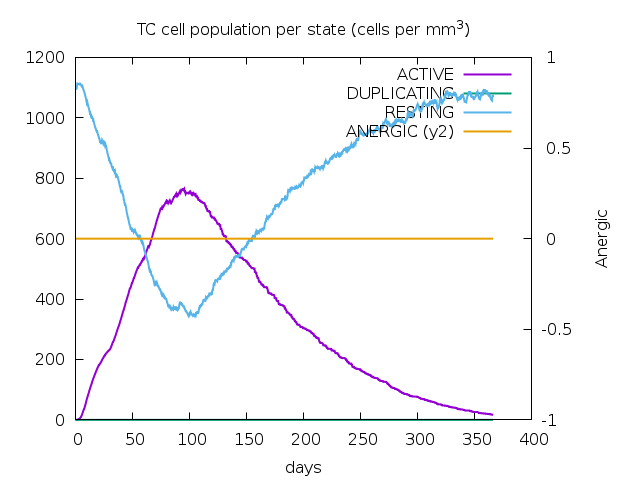

**Figure S19: SPVC_387 without adjuvants+HIS-tag:A)Antigen and immunoglobulin counts B)The changes observed in B-cell populations after given three injections, C) The development of T-helper, and D) T-cytotoxic cell populations per state after the injections**

**A	B**

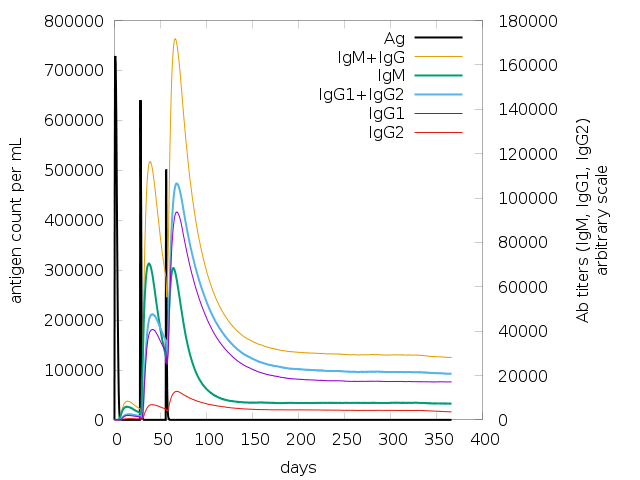

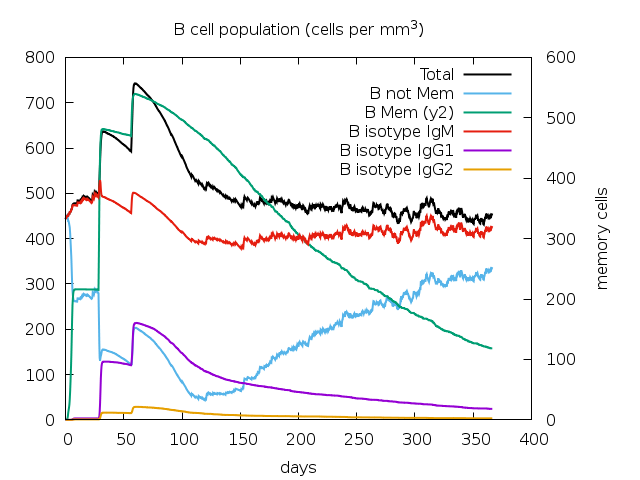

**C	D**

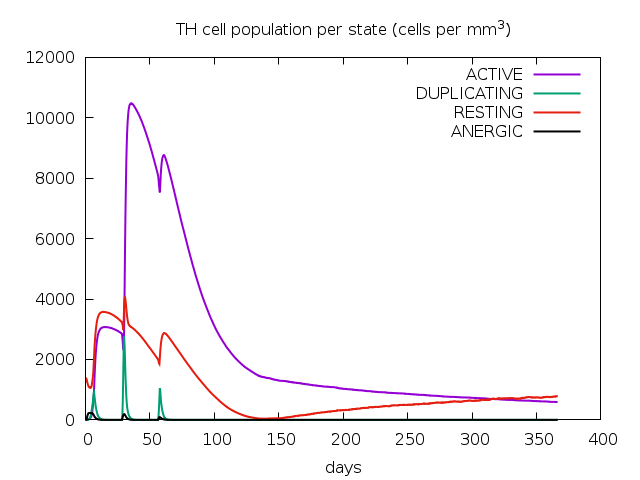

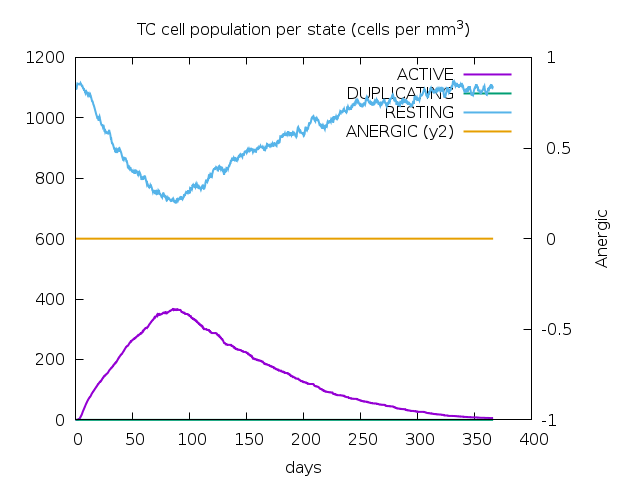

**Figure S20: SPVC_446 without adjuvants+HIS-tag:A)Antigen and immunoglobulin counts B)The changes observed in B-cell populations after given three injections, C) The development of T-helper, and D) T-cytotoxic cell populations per state after the injections**
